# High-throughput quantification of huntingtin mRNA expression and aggregation in mouse brain using automated RNAscope imaging

**DOI:** 10.64898/2026.02.09.704866

**Authors:** Pieter van Velde, Brian Tran, Sarah Allen, Eric Luu, Raymond Furgal, Ashley Summers, Jillian Belgrad, Emily Knox, Anastasia Khvorova, David Grunwald

## Abstract

Huntington’s disease (HD) is a repeat-associated neurodegenerative disorder traditionally characterized by toxic protein pathology resulting from expanded CAG repeats in the huntingtin (HTT) gene. In recent years, however, studies have identified repeat expansion–driven RNA pathology as an additional and potentially independent contributor to disease. In particular, mutant HTT transcripts containing expanded CAG repeats accumulate in the nucleus and form discrete RNA clusters, a feature shared with several other repeat-associated disorders. While protein aggregation and its downstream consequences have been extensively studied, our current understanding of the composition, organization, and dynamics of these nuclear mRNA clusters remains limited. Progress in this area has been constrained in part by the lack of robust methods to detect and quantify expanded HTT transcripts at single-molecule resolution within intact tissue. As a result, the contribution of RNA clustering to disease mechanisms, its relationship to repeat length, and its interaction with other pathological features of HD remain poorly defined. Here we present a high-throughput RNAscope™ pipeline that combines automated confocal imaging with rigorous microscope characterization to quantify both single mRNA molecules and multi-transcript clusters in fixed mouse brain tissue. Using 3D Gaussian point-spread function (PSF) fitting calibrated on 200 nm fluorescent beads and pointilistic image features from tissue data, we establish per-slide intensity thresholds from negative controls and normalize experimental signals to single-molecule reference intensities. The critical validation of our approach operates at two scales: for single molecules, the linear relationship between spot size and intensity (r^2^ > 0.90) reflects variable probe binding along transcripts; for clusters, the linear scaling between cluster volume and mRNA content (R^2^ > 0.98) confirms uniform probe accessibility and enables quantitative conversion of fluorescence intensity to absolute mRNA counts. Applied to HttQ111+/− knock-in mice across multiple ages, we analyzed thousands of fields of view (FOVs), detecting >900,000 single mRNA molecules and segmenting >1.9 million mRNA clusters using two probes targeting mouse huntingtin (Htt): one detecting the spliced transcript that uses early cryptic polyadenylation sites in intron 1 (HTT1a), and one detecting full-length Htt (fl-HTT). Our analysis reveals considerable heterogeneity in mRNA accumulation: 16-63% of Q111 FOVs are classified as “extreme” (exceeding the 95th percentile of wildtype clustered mRNA levels), with striatum showing higher prevalence than cortex for both probes (HTT1a: 63% striatum, 31% cortex; fl-HTT: 44% striatum, 16%cortex). Extreme FOVs are characterized by elevated cluster numbers (2-6× more clusters per nucleus) and higher cluster density (1.3-1.7× more mRNA per µm^3^). Cluster localization shows nuclear bias (∼68%) in normal FOVs, but extreme FOVs exhibit a shift toward cytoplasmic localization, particularly for fl-HTT (48% nuclear vs 68% in normal FOVs), though the interpretation of this shift requires further investigation. Despite the large dataset at the cellular level, our study included only 11 mice (9 Q111, 2 wildtype), and this limited sample size precluded robust statistical inference at the animal level. Nevertheless, these quantitative metrics provide a framework for investigating disease mechanisms and evaluating therapeutic interventions using RNAscope in future studies with larger cohorts.

## 1 Introduction

RNAscope™ is an in situ hybridisation (ISH) method that addresses the long-standing trade-off between sensitivity and specificity in RNA detection. Conventional ISH often produces high background and limited sensitivity, whereas PCR-based methods lose spatial context. RNAscope pairs a stringent probe architecture with a branched amplification cascade: each “double Z” probe comprises two adjacent oligonucleotides that must bind contiguously to the target RNA before a pre-amplifier can attach, preventing amplification from partial or non-specific binding [1]. With this design, researchers visualise individual RNA molecules as discrete puncta using chromogenic or fluorescent readouts; the assay works on formalin-fixed tissue sections and supports multiplex detection of three to four targets per section [1, 2]. According to the manufacturer, each RNAscope punctate dot represents a single copy of an mRNA molecule; however, there is variation in dot intensity and size due to differences in the number of ZZ probes bound to each target molecule [3]. The manufacturer therefore recommends that “the number of dots is critical and not the intensity/size of the dot(s).” This guidance has shaped how the field uses R NAscope: current practice focuses on counting individual, well-separated puncta rather than measuring their intensities, which works well for single-molecule quantification but fails when multiple mRNAs cluster together. For mRNA clusters, ranging from just a few transcripts to dozens or even hundreds of co-localised molecules, simple dot counting cannot distinguish a single bright punctum from a cluster of many molecules, creating a fundamental measurement gap for pathologies driven by transcript clustering. In this work, we explicitly deviate from this manufacturer recommendation: our calibration framework validates that fluorescence intensity, when properly normalized, provides quantitative mRNA counts. This validation rests on three elements: (1) per-slide calibration using negative controls to establish detection thresholds, (2) modal intensity normalization using isolated single molecules as an internal standard, and (3) the resulting linear relationship between cluster volume and mRNA content (*R*^2^ > 0.98), which would not hold if intensity were not quantitative. This calibrated intensity-based approach enables quantification of multi-transcript clusters that simple dot counting cannot resolve.

We apply RNAscope to Huntington disease (HD) models to demonstrate this approach. In HD, expanded CAG repeats in *HTT* transcripts cause aberrant splicing: the expanded repeat impairs normal splicing at the exon 1intron 1 junction, leading to production of a truncated transcript that retains part of intron 1 and encodes a highly pathogenic exon 1 protein fragment [5, 19]. These mutant transcripts also form stable secondary structures that contribue to nuclear retention and foci formation [28]. Related RNA-mediated toxic-ity occurs in *C9orf72*-linked amyotrophic lateral sclerosis/frontotemporal dementia (ALS/FTD), where expanded G_4_C_2_ repeats form nuclear RNA foci [6]. Our model is the HttQ111+/− knock-in mouse, which carries a single expanded allele and exhibits progressive striatal-specific transcriptional dysregulation and neuronal intranuclear inclusions despite minimal neurodegeneration up to 12 months [17]. Beyond genotype, HD pathology varies by anatomical location: in human post-mortem studies, striatal medium spiny neurons show early vulnerability while cortical pyramidal cells are relatively spared, and within each hemisphere, neurodegeneration progresses along a dorsal-to-ventral gradient within the striatum [7]. We therefore multiplex DARPP-32 labelling to mark medium spiny neurons (MSNs), the principal striatal neuron type, and combine it with DAPI nuclear counterstain to assess whether transcripts localize to the nucleus or cytoplasm. This cell-level and region-level heterogeneity demands comprehensive spatial sampling. Traditional approaches (manually selecting a handful of “representative” fields) risk systematic bias and cannot capture gradients of pathology across anatomical structures. Slide-scanner microscopy allows us to image entire sections, but this raises a practical question: is whole-brain coverage necessary, or can a few well-chosen slices suffice? By registering our data to the Allen Brain Atlas coordinate framework, we can test whether sub-sampling introduces artefacts and determine the minimal imaging strategy needed to reliably map regional vulnerability.

Here we present an end-to-end pipeline that integrates semi-automated slide-scanner imaging with a rigorous, quality-controlled analysis workflow to quantify mRNA clustering in HttQ111+/− mouse brain tissue using RNAscope. The primary contribution of this work is methodological: we describe a complete framework for quantifying both single mRNA molecules and multi-transcript clusters, with rigorous calibration and quality control at each stage. Each slide contains three adjacent brain sections: one hybridized with the experimental probe set targeting mouse *Htt* transcripts (a probe for the aberrantly spliced transcript that uses early cryptic polyadenylation sites in intron 1 [HTT1a] and a probe for full-length *Htt* [fl-HTT], along with DARPP-32 RNAscope probe for anatomical segmentation), one with positive control probes (POLR2A and UBC), and one with a negative control probe (non-mammalian bacterial DapB). This design enables within-slide technical validation, where positive controls establish assay sensitivity and dynamic range, while negative controls empirically determine detection thresholds for filtering background fluorescence. DARPP-32 staining in the experimental sections enables precise anatomical segmentation, distinguishing striatum from cortex and allowing region-specific analysis within each hemisphere along the rostro-caudal axis using atlas coordinates. Through high-throughput imaging of thousands of fields of view across anatomically defined cortical and striatal subregions, we first establish a single-molecule intensity calibration from isolated puncta, then characterize the transition from diffuse single molecules to high-intensity clusters using automated spot detection and spatial clustering. Our multi-tiered quality control (QC) framework computes slide-specific detection thresholds from negative control intensity distributions, cross-validates experimental measurements against positive control expression on the same slide, and traces outliers to distinguish technical failures from biological variation. We applied this pipeline to tissue from 9 HttQ111/+ mice (3 per age group at 2, 6, and 12 months) and 2 wildtype controls (at 2 and 12 months). We processed 30 slides total (with multiple non-continuous 40 *µ*m-thick sections per animal), which were used for calibration and quality control analyses. After quality control filtering (described in section 2.1), we acquired 1,617 fields of view, each imaged in four color channels (DAPI nuclear stain, DARPP-32 striatal marker, HTT1a, and fl-HTT), detected more than 900,000 single mRNA molecules, and segmented nearly 2 million mRNA clusters. We observe age-dependent increases in mRNA clustering for both fl-HTT and the pathogenic HTT1a splice variant, with substantial heterogeneity across tissue regions. However, we emphasize that while the dataset is large at the cellular level, the number of animals (particularly wildtype controls, n = 2) limits the strength of biological conclusions; the findings presented here should be considered preliminary observations that demonstrate the utility of the method rather than definitive characterization of HD progression. This approach combines statistical power from large-scale imaging with single-molecule resolution to map regional vulnerability and progression of huntingtin mRNA clustering in disease-relevant brain structures.

## 2 Results

### 2.1 Experimental design and data acquisition

We imaged HttQ111+/− mice across multiple ages and anatomical regions. These mice were derived from control groups of a therapeutic study [50] and received either no treatment (2-month-old) or control injections (aCSF or NTC siRNA; see section 4.1 and Table 1). Each slide contained three coronal sections: one with experimental RNAscope probes, one with positive controls, and one with negative controls. The experimental probe configuration consisted of: C1: mm-HTT-intron1 conjugated to Alexa Fluor 488 (this probe targets intron 1 sequences, detecting both the aberrantly spliced HTT1a transcript and nascent pre-mRNA), C2: DARPP-32 conjugated to Alexa Fluor 647, C3: full-length mm-HTT (hereafter fl-HTT) conjugated to Alexa Fluor 546 (this probe targets the 3’ UTR and detects full-length transcripts but not HTT1a), and a DAPI nuclear stain (probe target sequences in table S5 and Supplementary Note 4). Throughout this work, we refer to the 488 nm channel as the green channel and the 546 nm channel as the orange channel. These probes detect both wild-type (*mmHtt*) and mutant (*mmHtt1a*) transcripts, while DARPP-32 RNAscope signal provided anatomical segmentation. We observed that single-molecule detection was challenging in the Alexa 647 channel due to minimal intensity differences between specific signal and non-specific binding events in the negative control; therefore, we used DARPP-32 in this channel as a qualitative rather than quantitative measure. Positive control sections employed POLR2A in channel C1 (a low-abundance housekeeping gene), PPIB in channel C2 (a medium-abundance housekeeping gene), and UBC in channel C3 (a high-abundance housekeeping gene). Negative control sections utilized probes targeting the bacterial *dapB* gene, which has no mammalian targets. Each detection channel had its own negative control probe: dapB-C1, dapB-C2, and dapB-C3. We processed tissue from 9 Q111 mice (3 at 2 months, 3 at 6 months, 3 at 12 months) and 2 wildtype controls (1 at 2 months, 1 at 12 months), totaling 30 slides across all animals (1-4 slides per mouse). Each slide contained adjacent sections hybridized with different probe sets. All 30 slides had intact negative control sections, which were used to establish detection thresholds. Only 26 slides had intact experimental probe sections (HTT1a, fl-HTT, and DARPP-32), as tissue had fallen off during processing on 4 slides; these 26 slides were used for PSF calibration and single-molecule characterization (fig. 1). For biological analysis of mRNA expression and clustering (figs. 2 to 5), additional quality control filtering removed slides with insufficient tissue integrity (torn tissue, scratched tissue, etc.), leaving 15 Q111 slides (from 8 mice) and 3 WT slides (from 2 mice), generating 1,334 Q111 and 283 wildtype unique imaging positions.

**Table 1:** Mouse ID mapping. Anonymized IDs used in Figures 3F and 4F correspond to individual mice. Q_111_#1-Q_111_#9 denote HttQ111+/− mice; WT#1-WT#2 denote wildtype controls. Treatment abbreviations: UNT = untreated, NTC = non-targeting control siRNA, aCSF = artificial cerebrospinal fluid (see text for details). Multiple slides per mouse are denoted with decimal notation (e.g., Q_111_#1.1, Q_111_#1.2).

| ID | Sample | ID | Sample |
| --- | --- | --- | --- |
| Q<sub>111</sub> #1 | Q111 2mo #1 UNT | Q<sub>111</sub> #6 | Q111 6mo #3 NTC |
| Q<sub>111</sub> #2 | Q111 2mo #2 UNT | Q<sub>111</sub> #7 | Q111 12mo #1 aCSF |
| Q<sub>111</sub> #3 | Q111 2mo #3 UNT | Q<sub>111</sub> #8 | Q111 12mo #2 aCSF |
| Q<sub>111</sub> #4 | Q111 6mo #1 NTC | Q<sub>111</sub> #9 | Q111 12mo #3 aCSF |
| Q<sub>111</sub> #5 | Q111 6mo #2 NTC |  |  |
| WT#1 | WT 2mo #2 UNT | WT#2 | WT 12mo #3 aCSF |

**Fig. 1:**
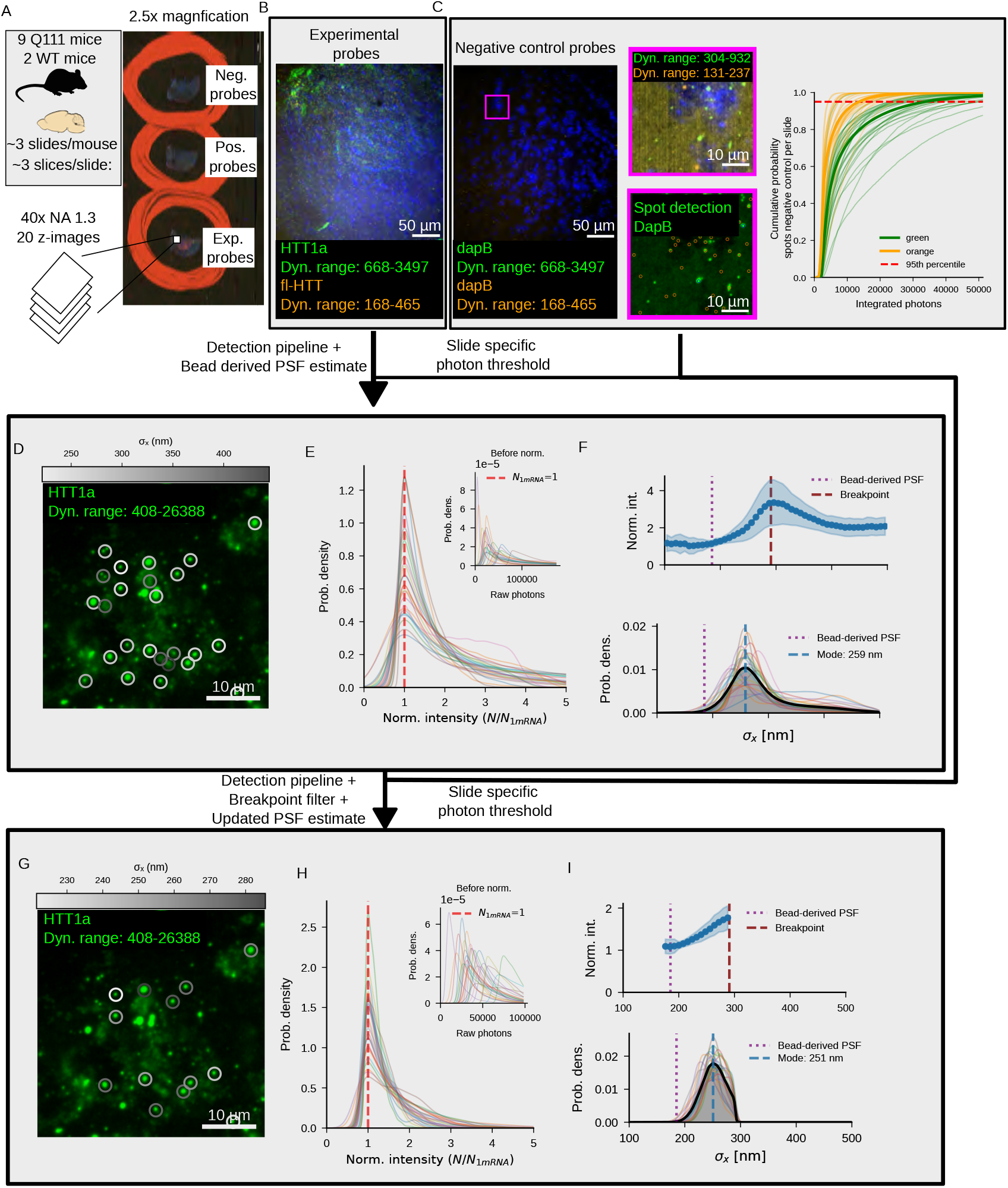
Establishing detection thresholds and single-molecule criteria through multi-stage calibration. **(A)**Experimental design overview. Left: 9 Q111 and 2 WT mice, 1 to 4 slides per mouse, 3 tissue sections per slide, imaged at 40× (NA 1.3) with 20 z-planes per FOV (500 nm spacing). Right: 2.5× magnification overview showing slide layout with three adjacent coronal sections stained for negative control probes (bacterial DapB), positive control probes (housekeeping genes), and experimental probes (HTT). **(B)** Representative FOV from experimental probe section. HTT1a (green, 488 nm) and fl-HTT (orange, 548 nm) with DAPI nuclear counterstain (blue). Dynamic ranges indicated for each channel. Scale bar: 50 *µ*m. **(C)** Negative control threshold determination. Left: FOV from DapB section displayed at same dynamic range as (B), showing minimal specific signal. Top-right inset: zoom at reduced dynamic range to visualize low-level background fluorescence. Bottom-right inset: same region with spot detection overlay; these detected spots are used to generate the CDF. Far right: CDFs of integrated photon counts from DapB sections (n = 727,329 spots, 30 slides). Faint lines: individual slides; bold lines: channel means (green = 488 nm; orange = 548 nm). Red dashed line: 95th percentile threshold. Thresholds computed per slide-channel to account for local imaging conditions. **(D-F)** *Stage 1: Initial detection using bead PSF*. Spots detected using bead-derived PSF (*σ*_*x*_ = 185 nm) as initial parameters, with minimal filtering (PFA < 0.05, intensity > threshold) and no size restrictions. **(D)** Representative FOV showing detected spots color-coded by fitted *σ*_*x*_ (250-400 nm range). Circles indicate all detected spots before size filtering, including both single molecules and unresolved clusters. Scale bar: 10 *µ*m. **(E)** Normalized intensity distributions (n = 1,819,916 spots, 26 slides, green channel). Main: distributions with modal intensity set to *N*_1mRNA_ = 1.0. Inset: raw distributions showing 2-3× slide-to-slide variation before normalization. **(F)** Size-intensity relationship. Top: mean normalized intensity vs. *σ*_*x*_ showing increase from bead PSF to breakpoint, then plateau (model mismatch). Purple dotted line: bead PSF (185 nm); red dashed line: breakpoint. Bottom: *σ*_*x*_ probability density with mode at 259 nm. **(G-I)** *Stage 2: Detection with breakpoint filtering*. Spots detected with breakpoint-based size filtering (*σ*_*x*_ < 290.8 nm) to isolate single molecules. **(G)** Same FOV as (D) after applying size filter. Color scale now spans 230-280 nm, showing uniform single-molecule population. Larger clusters excluded. Scale bar: 10 *µ*m. **(H)** Normalized intensity distributions (n = 249,526 spots, 26 slides, green channel). Successful collapse around *N*_1mRNA_ = 1.0. Inset: raw distributions before normalization. **(I)** Size-intensity relationship validates single-molecule identification. Top: flat intensity profile below breakpoint confirms homogeneous population. Bottom: narrower *σ*_*x*_ distribution (mode 251 nm) after filtering. Technical parameters: pixel size 162.5 nm, slice depth 500 nm, size lower bounds 80% of bead-derived PSF for all dimensions (*σ*_*x*_ ≥ 148 nm, *σ*_*y*_ ≥ 150 nm, *σ*_*z*_ ≥ 458 nm), PFA < 0.05. Figure S1 and Supplementary Note 1: per-slide vs. pooled threshold comparison. Figures S2 to S3: complete breakpoint analysis for both channels (green and orange) across all three dimensions (*σ*_*x*_, *σ*_*y*_, *σ*_*z*_). Figures S4 to S5: detailed size and intensity distributions for single-molecule filtering in both channels.

**Fig. 2:**
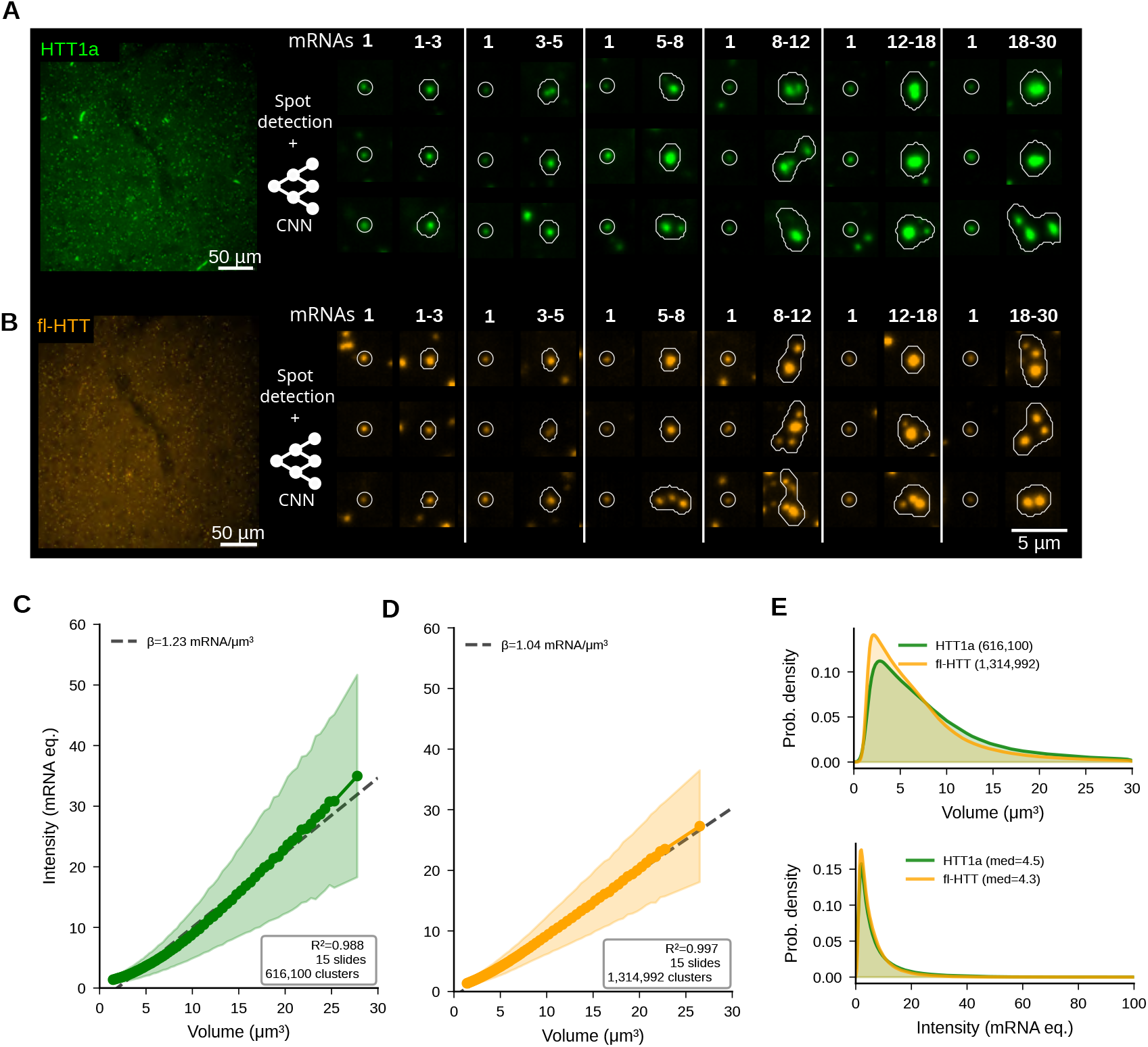
Single mRNA detection and CNN-based cluster segmentation capture the full range of transcript aggregation states. **(A)** HTT1a probe (488 nm, green). Left: representative FOV overview showing spatial distribution of mRNA signal. Scale bar: 50 *µ*m. Right: paired comparisons of single mRNAs (left column of each pair, labeled “1”) versus clusters of increasing mRNA content (right column, labeled by mRNA equivalents: 1-3, 3-5, 5-8, 8-12, 12-18, 18-30). Two rows show spot detection (circles) and CNN segmentation (white outlines). Single spots appear diffraction-limited while clusters show extended morphologies. Scale bar for zooms: 5 *µ*m. **(B)** fl-HTT probe (548 nm, orange). Same layout as (A). **(C)** Green channel (HTT1a) cluster intensity (in mRNA equivalents) versus cluster volume. Each point represents the mean intensity of all clusters within a volume bin; shading shows ±1 SD across slides. Dashed line: linear fit with slope *β* = 1.23 mRNA/*µ*m^3^, *R*^2^ = 0.988 (n = 616,100 of 640,320 clusters within 0 to 30 *µ*m^3^ volume range, 15 slides). Linear scaling validates quantitative mRNA counting and confirms probe accessibility throughout clusters. **(D)** Orange channel (fl-HTT) cluster intensity versus volume, same format as (C). Linear fit: *β* = 1.04 mRNA/*µ*m^3^, *R*^2^ = 0.997 (n = 1,314,992 of 1,337,908 clusters within 0 to 30 *µ*m^3^, 15 slides). Similar slopes between channels suggest comparable mRNA packing density. **(E)** Distribution panels. Top: cluster volume distributions (KDE). Bottom: cluster intensity distributions in mRNA equivalents; medians of 4.5 (HTT1a) and 4.3 (fl-HTT). Data from Q111 animals only (15 slides). Cluster filtering: pruning removed 16.7% green (140,619/843,092) and 26.6% orange (499,176/1,875,933) clusters overlapping single-molecule spots; intensity/CV threshold removed 8.8% green (62,153/702,473, mean threshold 33,932 photons) and 2.8% orange (38,849/1,376,757, mean threshold 13,210 photons). Volume scaling: 0.0132 *µ*m^3^/voxel. Extended analyses in figs. S8 to S12.

### 2.2 Establishing detection thresholds through negative control calibration

Quantifying mRNA at single-molecule resolution in fixed tissue presents unique challenges: probe hybridization efficiency varies between tissue types and preparation batches, autofluorescence contributes variable background, and the transition from isolated transcripts to multi-molecule clusters must be objectively defined. We address these challenges through a well-characterized, multi-stage pipeline that establishes per-slide control signals, distinguishes single molecules from clusters based on empirical size criteria, and controls for non-specific binding using negative controls (fig. 1).

The foundation of our quantitative approach is rigorous background characterization, based on detector characterization and conversion of grey levels to photons (see section 4.2.1). We computed per-slide, per-channel photon thresholds as the 95th percentile of integrated intensities from negative-control detections, i.e., spots detected on tissue sections hybridized with bacterial DapB probes that have no mammalian targets (n = 727,329 spots across 30 slides; see section 4.4 for detection pipeline details; fig. 1C; fig. S1). This threshold represents the intensity below which detected signals cannot be distinguished from non-specific binding and autofluorescence. Critically, we observed substantial slide-to-slide variation in these thresholds: coefficient of variation (CV) = 39.8% for the green channel (mean ± SD: 32,955 ± 13,117 photons) and CV = 41.9% for the orange channel (12,384 ± 5,194 photons). This variation motivated our per-slide normalization strategy: applying a global threshold would systematically bias results across slides. We validated that negative control thresholds showed no significant correlation with experimental mRNA expression levels across all four channel-region combinations: HTT1a in cortex (*r* = −0.273, *p* = 0.345), HTT1a in striatum (*r* = −0.283, *p* = 0.307), fl-HTT in cortex (*r* = +0.255, *p* = 0.379), and fl-HTT in striatum (*r* = +0.284, *p* = 0.305; mean |*r*| = 0.274, all *p* > 0.3; fig. S7). This weak/absent correlation is critical: it demonstrates that the biological variation we observe in HTT expression across slides is not driven by differences in negative control thresholds. Slides with higher background thresholds do not systematically show higher or lower HTT expression, confirming that observed differences in huntingtin transcript levels reflect true biological variation rather than technical artifacts of threshold determination.

For each slide, we imaged a subregion within the cortex and a subregion within the striatum, allowing us to examine whether background thresholds differed systematically between these brain regions. Comparing 58 paired slide-channel combinations across cortex and striatum revealed no significant regional difference: cortex thresholds averaged 23, 037 ± 14, 661 photons versus striatum 21, 938 ± 15, 334 photons (mean difference +1, 099 ± 6, 469 photons; paired *t*-test: *t* = 1.294, *p* = 0.201; fig. S14). Per-channel analysis confirmed this pattern: green channel cortex vs striatum *p* = 0.481; orange channel *p* = 0.100. Age showed no effect on thresholds (all correlations |*r*| < 0.1, *p* > 0.5). Rostrocaudal position along the anterior-posterior axis also showed negligible correlations with thresholds in either region individually (cortex: *r* = −0.052, *p* = 0.70; striatum: *r* = +0.087, *p* = 0.52), indicating that the precise anatomical placement of the negative control imaging region does not meaningfully affect threshold determination. Given the absence of significant regional or positional effects, we merged the cortex and striatum measurements to compute a single per-slide threshold for subsequent analyses. An alternative cost-saving strategy (imaging negative controls on only a subset of slides and applying the pooled threshold to all slides) is evaluated in Supplementary Note 1 and figs. S22 to S24.

### 2.3 PSF calibration and single-molecule identification

Initial PSF calibration from 200 nm fluorescent TetraSpeck microspheres yielded point-spread function widths of *σ*_*x*_ = 185.0 nm, *σ*_*y*_ = 187.0 nm laterally, and *σ*_*z*_ = 573.0 nm axially for the green channel (40× oil objective, NA 1.3, spinning disk confocal; values for both green and orange channels in table S3). These values exceed the theoretical diffraction limit (*σ*_*x,y*_ ≈ *λ*/4NA ≈ 99 nm for 515 nm emission [30]), reflecting the finite bead size and non-optimal optical characteristics of the spinning disk system. However, RNAscope signals arise from branched amplification trees attached to ∼20 ZZ probe pairs that bind along the target mRNA sequence. Unlike point-like fluorophores, these extended signal sources may produce spots with different characteristics than bead-derived PSFs. We therefore employed a two-stage empirical recalibration strategy, focusing on the experimental probe set (HTT1a and fl-HTT), to determine appropriate size criteria for single-molecule detection.

In the following, we illustrate the calibration procedure using the green channel (HTT1a); fig. 1D-I shows this channel only, with equivalent orange channel (fl-HTT) analysis in figs. S2 to S3. In the first stage, we used the bead-derived PSF (*σ*_*x*_ = 185 nm, *σ*_*y*_ = 187 nm, *σ*_*z*_ = 573 nm) as initial parameters for spot detection, then fitted each detected spot with a 3D Gaussian using flexible sigma values (see section 4.4 for detection and fitting details). Minimal filtering was applied at this stage: probability of false alarm (PFA) < 0.05 from the generalized likelihood ratio test (GLRT), and intensity above the per-slide negative control threshold. No size restrictions were applied. Raw integrated photon counts varied by 2-3× across slides due to differences in imaging parameters between sessions (tables S1 and S2), tissue autofluorescence, and probe hybridization efficiency (fig. 1E, inset). We normalized intensities per slide by identifying the modal intensity via kernel density estimation (KDE) with Scott’s rule bandwidth selection [43, 44], then scaling all intensities such that this mode equals *N*_1mRNA_ = 1.0 (fig. 1E, main panel). Formally, the normalized intensity *Ñ* for each spot is:

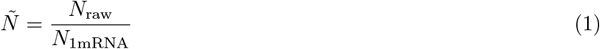

where *N*_raw_ is the integrated photon count from the 3D Gaussian fit, and *N*_1mRNA_ is the per-slide modal intensity representing a single mRNA molecule. This normalization assumes that the most common detected signal corresponds to a single mRNA molecule, a reasonable assumption given that the detection pipeline is optimized for single spots.

Plotting normalized intensity versus fitted spot size (fig. 1F, shown for the green channel using *σ*_*x*_), we observed that intensity increases linearly from the bead PSF size toward larger spots, up to a certain point, which we term the breakpoint (*σ*_*x*_ = 290.8 nm for the green channel). Beyond this breakpoint, further size increases yield no additional photon counts, a plateau reflecting model mismatch, where the Gaussian PSF model can no longer accurately describe larger structures. The breakpoint exceeds the bead PSF by 57%, reflecting the extended nature of RNAscope signal sources compared to point-like fluorophores. Figures S2 to S3 provide the complete breakpoint analysis for both channels (green and orange) across all three dimensions (*σ*_*x*_, *σ*_*y*_, *σ*_*z*_).

In the second stage, we used the modal PSF sizes from Stage 1 as the new detection parameters, with the empirically-determined breakpoints as size upper bounds (*σ*_*x*_ < 290.8 nm, *σ*_*y*_ < 291.1 nm, *σ*_*z*_ < 698.6 nm) and 80% of the bead-derived PSF as size lower bounds for all dimensions (spots smaller than this are likely noise or fitting artifacts). Applying these criteria to experimental tissue data (fig. 1H-I) yields intensity distributions that converge around *N*_1mRNA_ = 1.0 across all 26 slides (fig. 1H), confirming that our normalization procedure effectively removes technical variation. The size-intensity relationship (fig. 1I) shows a linear trend within the single-molecule regime, validating the breakpoint-based filtering. Detailed size and intensity distributions for single-molecule filtering are shown in fig. S4 (green channel) and fig. S5 (orange channel). Notably, the modal spot sizes after filtering differ between channels: the green channel (HTT1a) yields modes of *σ*_*x*_ = 252 nm, *σ*_*y*_ = 258 nm, *σ*_*z*_ = 674 nm, while the orange channel (fl-HTT) produces smaller modes of *σ*_*x*_ = 239 nm, *σ*_*y*_ = 238 nm, *σ*_*z*_ = 667 nm (5-8% smaller laterally). Since bead-derived PSF values are nearly identical between channels (table S3), this difference is not attributable to optical properties. Instead, it may reflect the shorter probe target region for fl-HTT (966 bp) compared to HTT1a (1,150 bp), with both target sequences detailed in table S5 and Supplementary Note 4, as 20 ZZ probe pairs distributed over a shorter mRNA segment would produce more spatially compact fluorescent signals. Other biological factors may also contribute, including differences in mRNA secondary structure, local folding environment, or probe binding efficiency along the target sequence. These refined single-molecule detections and the associated calibration parameters were used for all subsequent analyses.

This calibration framework enables quantitative intensity analysis of RNAscope data, a departure from conventional practice, which relies exclusively on counting discrete puncta. Within the single-molecule regime, we observe that larger spots exhibit proportionally higher fluorescence intensity, consistent with variable numbers of probe pairs binding to each target mRNA (the RNAscope amplification tree accommodates up to 20 double-Z probe pairs per target, but not all binding sites may be occupied on every transcript). Notably, the size distribution before breakpoint filtering (fig. 1F, bottom panel) exhibits a long tail extending toward larger sizes, reflecting unresolved clusters and model mismatch; after filtering (fig. 1I, bottom panel), the distribution becomes approximately normal, as expected for a homogeneous population of individual transcripts with stochastic variation in probe binding efficiency, further supporting our single-molecule assignment. By establishing per-slide control signals and empirically defining the boundary between single molecules and unresolved clusters, we can now convert integrated photon counts into absolute mRNA equivalents. This capability is essential for studying RNA clustering pathologies, where the biologically relevant signal lies not in the number of puncta but in the total transcript content of each cluster.

### 2.4 Quantifying mRNA clusters: linear intensity-volume scaling

Signals exceeding single-molecule size criteria cannot be resolved by PSF-based fitting and instead were captured by a 3D convolutional neural network trained to segment extended fluorescent regions (section 4.6). Crucially, single-molecule detection and cluster segmentation operate on the same images with overlapping sensitivity ranges, ensuring that no fluorescent signal is missed: isolated transcripts are captured by PSF fitting while extended clusters are captured by CNN segmentation. This overlap necessitates careful handling to prevent double-counting. We therefore implemented a conservative pruning strategy that serves as a double-safeguard: first, we catalog all potential signals comprehensively (including ambiguous cases during CNN training to maximize detection sensitivity), then we prune clusters that overlap with validated single-molecule coordinates. This two-stage approach ensures that single molecules are never double-counted as clusters while maintaining high sensitivity for genuine cluster detection. The pruning step removed 16.7% of green channel clusters (140,619/843,092) and 26.6% of orange channel clusters (499,176/1,875,933), indicating substantial spatial overlap between the detection modalities and validating the necessity of this safeguard. We then applied a two-stage intensity filter: (1) the same negative control intensity threshold used for single-molecule filtering, and (2) a CV threshold on voxel intensities within each cluster (≥ 0.5) to remove homogeneous low-intensity regions likely arising from background fluorescence rather than true mRNA signal. This filtering discarded an additional 8.8% of green (62,153/702,473) and 2.8% of orange (38,849/1,376,757) clusters. The final dataset comprised 640,320 green channel clusters and 1,337,908 orange channel clusters across 15 Q111 slides (see section 2.1 for slide exclusion criteria; fig. 2).

For each retained cluster 𝒞, we computed background-corrected integrated intensity using a local annular background estimate:

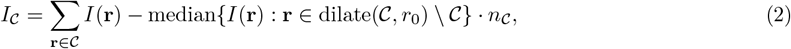

where **r** is the spatial coordinate of a voxel, *I*(**r**) is the fluorescence intensity at voxel **r**, *n*_𝒞_ is the number of voxels in the cluster, dilate(𝒞, *r*_0_) is the morphological dilation of 𝒞 by radius *r*_0_, and dilate(𝒞, *r*_0_)\ 𝒞 represents the annular region surrounding the cluster. This local background correction is critical in tissue, where fluorescence intensity can vary substantially over spatial scales of 10-20 *µ*m due to cellular and subcellular structures. We then expressed cluster signal in single-molecule equivalents using the per-slide normalization factor from eq. (1):

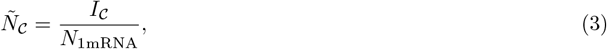

where *Ñ*_𝒞_ represents the mRNA content of cluster 𝒞 in units of single-molecule equivalents. This normalization places clusters on the same quantitative scale as single mRNAs and enables direct comparison across slides despite photon-yield variations.

The critical validation of our quantitative approach is the relationship between cluster volume and mRNA content. If our single-molecule normalization correctly converts fluorescence intensity to mRNA equivalents, and if RNAscope probes uniformly access mRNA within clusters, we expect cluster mRNA content (*Ñ*_𝒞_) to scale linearly with volume. Deviation from linearity would indicate either (1) probe penetration limitations at larger cluster sizes (causing saturation), or (2) size-dependent variation in mRNA packing density (for example, if larger clusters were systematically more or less densely packed than smaller ones). We observed strong linear relationships for both channels: *β* = 1.23 mRNA/*µ*m^3^ for the green channel (*R*^2^ = 0.988 on binned means) and *β* = 1.04 mRNA/*µ*m^3^ for the orange channel (*R*^2^ = 0.997 on binned means) (fig. 2C-D).

This linearity has two important implications. First, it demonstrates that RNAscope probe amplification trees can access mRNA molecules throughout clusters regardless of cluster size; if probe penetration were limiting, we would observe saturation (sublinear scaling) at larger volumes. The consistent linearity across volumes spanning two orders of magnitude (from sub-micron to >20 *µ*m^3^) indicates that even large clusters remain fully accessible to probe hybridization and branched amplification. Second, the similar slopes between channels ( ∼1.1 mRNA/*µ*m^3^ average) suggest comparable mRNA packing density regardless of transcript type (exon 1 vs full-length), providing internal consistency validation.

Cluster volume and intensity distributions are shown in fig. 2E: both channels showed similar right-skewed distributions with intensity medians of 4.5 (HTT1a) and 4.3 (fl-HTT) mRNA equivalents.

We observed per-slide variation in the volume-intensity slope *β* (CV = 36% green, 26% orange), which represents mRNA packing density within clusters (fig. S6). This variation could potentially be explained by our choice of *N*_1mRNA_ (the per-slide modal intensity we select as the single-molecule reference for normalization). If so, this would be problematic, as we would be introducing artificial variability through our normalization procedure rather than measuring true biological differences. To test this, we examined whether *N*_1mRNA_ correlates with *β*. We observed no significant correlation between per-slide peak intensity and slope for either channel (green: *r* =− 0.37, *p* = 0.17; orange: *r* = 0.04, *p* = 0.89). The observed inter-slide variability in *β* therefore reflects genuine biological or statistical variance across different tissue sections and animals, rather than artifacts of the normalization procedure.

### 2.5 Regional expression patterns and individual variation

Our analysis pipeline establishes a two-tier quantification strategy that captures the full spectrum of mRNA expression states. Single mRNAs are counted directly when they conform to tissue-calibrated PSF dimensions and pass stringent statistical filters (*p*_FA_ < 0.05, size within breakpoint thresholds, intensity above negative-control threshold). Clusters, detected through 3D segmentation and expressed in single-molecule equivalents via the per-slide normalization factor *N*_1mRNA_, capture signal from densely packed or spatially unresolved transcript clusters (see section 4.6 for complete pipeline details). This slide-specific normalization ensures that both single molecules and clusters are quantified on a common, biologically interpretable scale that is robust to technical variation.

Throughout our analysis, we quantify three complementary measures of mRNA expression. For each region of interest ℛ containing *n* single molecules and *m* clusters, we define:

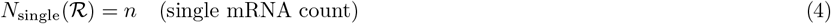

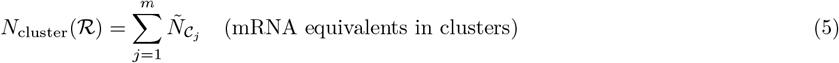

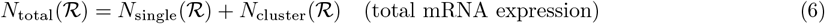

where 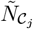 is the intensity of cluster *j* expressed in single-molecule equivalents (eq. (3)). This framework enables direct comparison across experimental conditions, brain regions, and time points, as all measurements are normalized to a common molecular scale.

To convert total mRNA expression to per-nucleus values, we segment nuclei from DAPI-stained images using a residual 3D U-Net architecture (section 4.6). Direct counting of individual nuclei is challenging in brain tissue due to the densely packed cellular architecture, where overlapping nuclei prevent accurate instance segmentation. We therefore estimate the number of nuclei per field of view 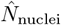 by dividing the total DAPI-stained volume by a mean nuclear volume of 716 *µ*m^3^, derived from manual diameter measurements (mean ±SEM: 10.6 ± 0.2 *µ*m; SD = 2.3 *µ*m; n = 120 nuclei; fig. S25; Supplementary Note 2). We then express mRNA per nucleus as:

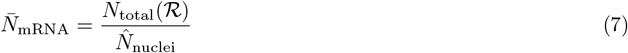

This normalization controls for variation in cell density across FOVs and enables comparison of expression levels across tissue regions.

Having established this quantitative framework, we examined mRNA expression across brain regions (fig. 3). We analyzed expression across 9 anatomical subregions: 4 striatal quadrants (dorsomedial, dorsolateral, ventrolateral, ventromedial) definitively identified via DARPP-32 marker expression, and 5 cortical areas (motor, somatosensory upper limb, somatosensory nose, gustatory/insular, and piriform) assigned based on anatomical landmarks alone. Unlike the striatal subregions, where DARPP-32 immunoreactivity provides unambiguous molecular boundaries, cortical area assignments relied on visual comparison to the mouse brain atlas without cell-type-specific markers; consequently, there is inherent uncertainty in the precise cortical area designations, and some FOVs may be misclassified. One-way ANOVA revealed no significant differences between subregions within cortex or striatum, nor between cortex and striatum as major regions, for either transcript (all *p* > 0.05; fig. 3B; fig. S17). Given this uniformity, we averaged all FOVs within each region for subsequent analyses. Rostrocaudal variation is examined separately below (fig. 3E).

**Fig. 3:**
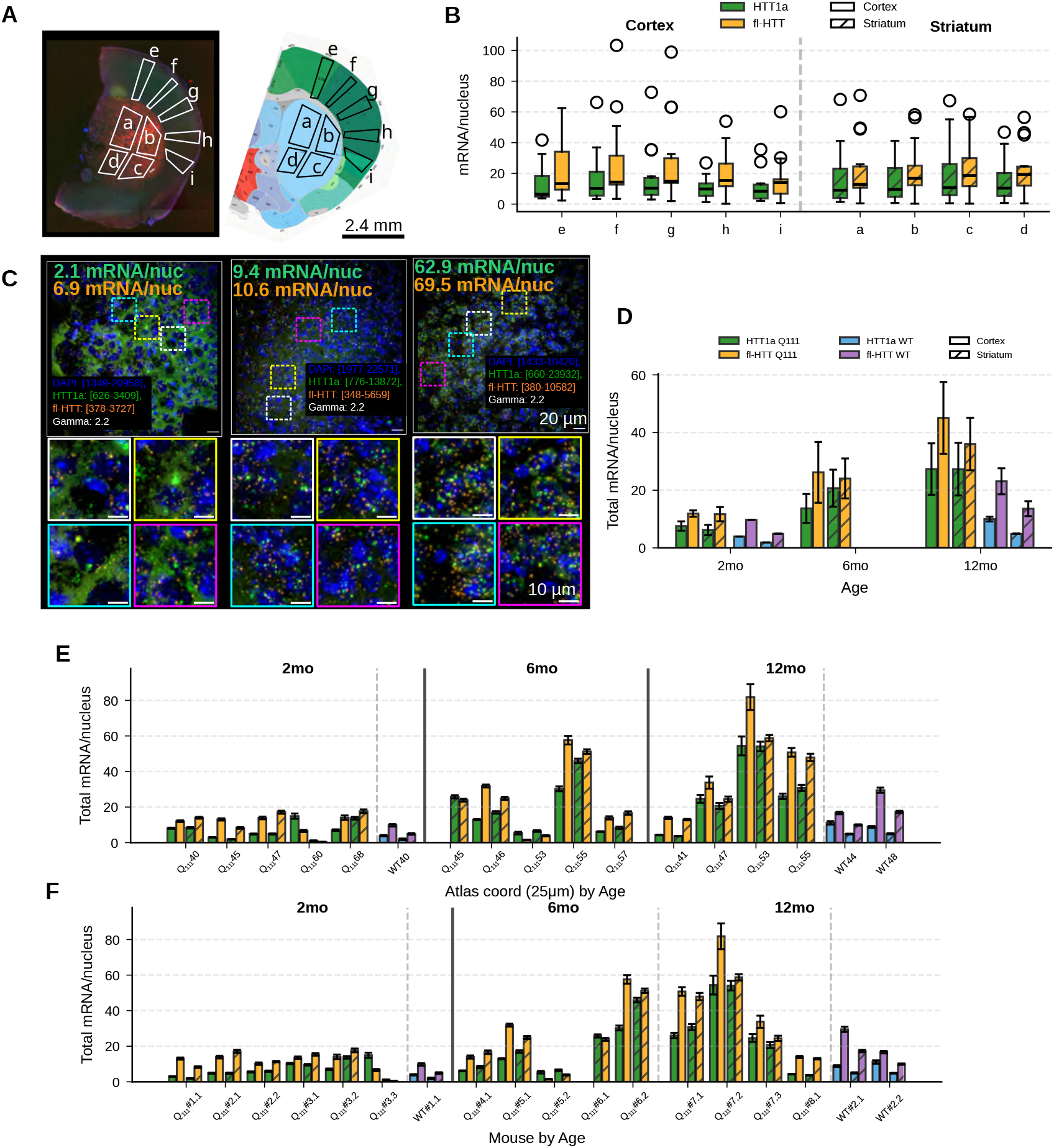
Regional expression patterns and total mRNA quantification in Q111 and WT mice. **(A)** Anatomical overview. Left: striatum (red) identified via DARPP-32 marker; yellow box shows imaging FOV locations. Right: schematic with subregion labels (a-i), adapted from the Allen Mouse Brain Atlas [32]. Striatal (DARPP-32 identified): a = dorsomedial, b = dorsolateral, c = ventrolateral, d = ventromedial. Cortical (approximate, no cell markers): e = motor (M1/M2), f = somatosensory upper limb (S1), g = somatosensory nose (S1), h = gustatory/insular, i = piriform. **(B)** Sub-regional expression (n = 250 measurements). X-axis: cortical (e-i, left of dashed line) and striatal (a-d, right) subregions. Box plots: median, IQR, 1.5×IQR whiskers. Green: HTT1a; orange: fl-HTT. Solid: cortex; hatched: striatum. **(C)** Representative FOV images: low, medium, high expression. Expression values annotated (green = HTT1a, orange = fl-HTT, mRNA/nucleus). DAPI nuclear stain in blue. Images displayed with gamma correction 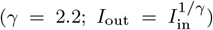 to enable simultaneous visualization of dim single molecules and bright clusters; the intensity scale is therefore non-linear, and bright spots are substantially brighter in the raw data than they appear. Dynamic ranges (raw image intensity values, not photon-corrected) shown at bottom of each panel. **(D)** Total mRNA by age (2, 6, 12 months). Each point represents one slide (per-slide mean of FOVs); bars show group median. Q111: green/orange; WT: blue/purple. Solid: cortex; hatched: striatum. Mann-Whitney U tests: no significant differences between ages, regions, or genotypes (all *p* > 0.05), likely due to limited sample sizes (n = 4-6 slides per group). **(E)** Expression by approximate atlas coordinate (assigned by visual comparison to Allen Mouse Brain Atlas coronal sections spaced at 100 *µ*m intervals [32]), grouped by age. **(F)** Per-mouse breakdown, grouped by age. IDs: Q_111_#1-Q_111_#9, WT#1-WT#2; see Table 1 for mapping. Data: 1,334 Q111 FOVs (2,668 including both channels), 283 WT FOVs (566 including both channels). Extended analysis in figs. S12 to S13, figs. S17 to S19.

We examined total mRNA expression across ages (2, 6, and 12 months) in Q111 mice (fig. 3D; fig. S18). To avoid pseudoreplication, we first averaged FOVs within each slide, then compared per-slide means across conditions. Although median expression values increased consistently with age across all conditions (e.g., Q111 striatum HTT1a: 5.4 mRNA/nucleus at 2 months, 17.0 at 6 months, 25.7 at 12 months), pairwise Mann-Whitney U tests between consecutive ages did not reach statistical significance (all *p* > 0.05), likely due to limited statistical power with small sample sizes (n = 4-6 slides per age group). Similarly, comparisons between cortex and striatum within each age and genotype, and between Q111 and wildtype within each region and age, showed no significant differences (Mann-Whitney U tests, all *p* > 0.26). Median total mRNA levels in Q111 at 12 months: cortex HTT1a 25.3, fl-HTT 42.3 mRNA/nucleus; striatum HTT1a 25.7, fl-HTT 36.2 mRNA/nucleus.

Critically, examination of individual animals revealed substantial inter-mouse variability (fig. 3F; fig. S19): some Q111 mice showed expression levels comparable to wildtype, while others exhibited dramatically elevated mRNA accumulation (>50 mRNA/nucleus vs <10 in low-expressing mice). This variability suggests that mean expression values may obscure biologically important heterogeneity. The majority of HTT signal arose from clustered rather than single mRNA molecules: single mRNA contributed <5% of total expression (mean single mRNA: 0.39-0.49 mRNA/nucleus for HTT1a, 1.77-1.84 for full-length; figs. S12 to S13), indicating that most HTT signal arises from clusters, defined as any fluorescent signal above threshold that did not qualify as a single molecule according to our size criteria (section 4.5).

Given the limited number of slides per condition (n = 4-6) and even fewer mice, we lacked statistical power to draw firm conclusions about age or genotype effects at the animal level. To gain additional insight into HTT expression heterogeneity in Q111 mice, we therefore examined variation at the level of individual fields of view, focusing on clustered mRNA abundance and cluster properties.

### 2.6 FOV-level analysis reveals heterogeneous mRNA accumulation

To move beyond mean expression values and characterize the distribution of mRNA accumulation at finer spatial resolution, we analyzed individual FOVs and compared Q111 distributions to wildtype (fig. 4A-C). We defined “extreme” FOVs as those with clustered mRNA per nucleus exceeding the 95th percentile (P95) of the wildtype distribution, a biologically grounded threshold that accounts for natural variation in healthy tissue. By definition, only ∼5% of wildtype FOVs exceed this threshold, so any substantial deviation in Q111 mice indicates disease-associated mRNA accumulation.

**Fig. 4:**
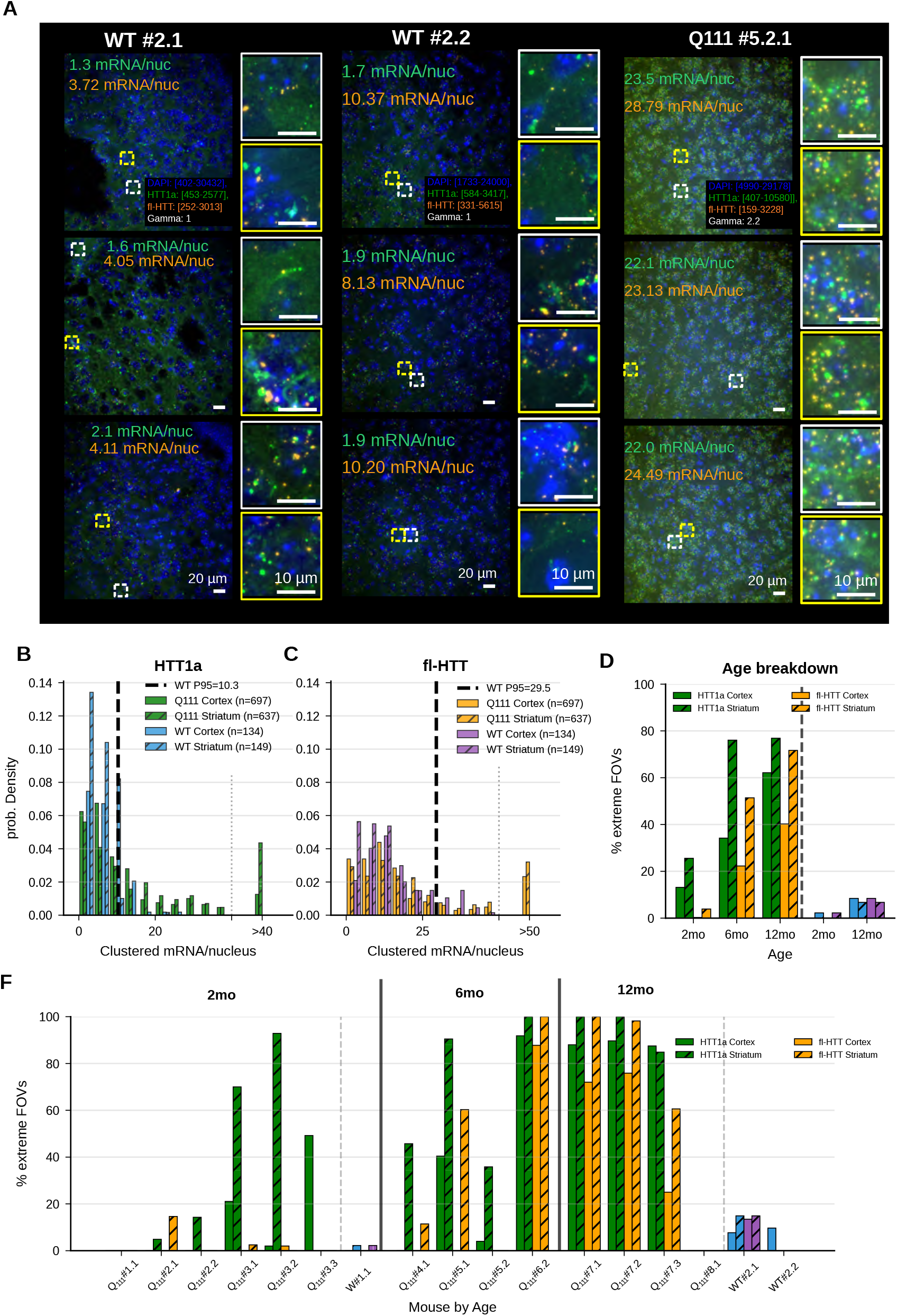
FOV-level variance and extreme outlier analysis reveals heterogeneous mRNA accumulation. **(A)** Representative FOV images comparing WT and extreme Q111 expression. Left two columns: WT mice (#2.1, #2.2) showing typical low-to-moderate expression (1-10 mRNA/nucleus). Right column: Q111 mouse (#5.2.1) showing an extreme FOV with elevated clustered mRNA (>20 mRNA/nucleus). Each column displays three FOVs with per-nucleus mRNA values annotated (green = HTT1a, orange = fl-HTT); insets show higher magnification of clusters. DAPI nuclear stain in blue. Q111 images displayed with gamma correction (*γ* = 2.2; 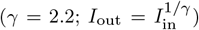) to enable simultaneous visualization of dim single molecules and bright clusters; the intensity scale is therefore non-linear, and bright spots are substantially brighter in the raw data than they appear. WT images displayed with linear scaling (*γ* = 1.0) as signal intensity is lower and does not require compression. Dynamic ranges (raw image intensity values, not photon-corrected) shown at bottom of each panel. Scale bars: 10 *µ*m (main), 20 *µ*m (insets). **(B)** HTT1a: clustered mRNA/nucleus distributions. Histograms with tail bin (>40) compressing outliers. Black dashed: WT P95 threshold (cortex: 12.80; striatum: 7.76 mRNA/nucleus). Q111 cortex: 31.1% extreme (217/697 FOVs), median 7.47; Q111 striatum: 62.6% extreme (399/637 FOVs), median 11.84. WT cortex: n = 134, median 7.47; WT striatum: n = 149, median 3.67. Mann-Whitney: cortex *p* = 7.58 × 10^−4^, striatum *p* = 4.57 × 10^−31^. **(C)** fl-HTT: same analysis. WT P95 thresholds: cortex 38.43, striatum 20.57. Q111 cortex: 16.1% extreme (112/697), median 13.52; striatum: 43.6% extreme (278/637), median 17.68. WT cortex median 13.47; WT striatum median 9.12. Mann-Whitney: cortex *p* = 7.99 × 10^−1^, striatum *p* = 3.70 × 10^−19^. **(D)** Extreme FOV % by age. Q111 left, WT right of dashed line. **(E)** Extreme FOV % by approximate atlas coordinate. **(F)** Per-mouse extreme FOV %, grouped by age. IDs consistent with Fig. 3F. Extreme = clustered mRNA/nucleus > WT P95 (per channel-region). Extended analysis in figs. S15 to S16.

We compared clustered mRNA per nucleus between Q111 and wildtype FOVs using Mann-Whitney U tests (fig. 4B-C). Wildtype P95 thresholds (dashed lines) were: HTT1a cortex 12.80, striatum 7.76 mRNA/nucleus; fl-HTT cortex 38.43, striatum 20.57 mRNA/nucleus. The lower striatal thresholds reflect lower baseline HTT expression in wildtype striatum.

This analysis revealed substantial heterogeneity in Q111 tissue. For HTT1a, 31.1% of cortical FOVs (217/697; *p* = 7.58 × 10^−4^) and 62.6% of striatal FOVs (399/637; *p* = 4.57 × 10^−31^) exceeded wildtype P95 thresholds, far exceeding the 5% expected by chance. For fl-HTT, 16.1% of cortical FOVs (112/697) showed elevated clustering, though this did not reach significance (*p* = 0.80), while 43.6% of striatal FOVs (278/637; *p* = 3.70 × 10^−19^) showed significant extreme clustering. The higher prevalence of extreme FOVs in striatum is consistent with known striatal vulnerability in HD, where medium spiny neurons show selective susceptibility compared to cortical neurons [16, 29]. We examined factors contributing to this heterogeneity (fig. 4D-F). The proportion of extreme FOVs increased with age in Q111 mice, while remaining low in wildtype controls (fig. 4D). Rostrocaudal position (atlas coordinate) showed no clear gradient, with extreme FOVs distributed throughout the sampled range (fig. 4E). Individual mouse analysis revealed substantial animal-to-animal variation (fig. 4F): some Q111 mice showed >80% extreme FOVs while others remained near wildtype levels. However, given our limited sample size (8 Q111 and 2 wildtype mice, 15 slides after quality filtering), these trends should be interpreted with caution; larger cohorts will be needed to determine whether age, anatomical position, or other factors reliably predict mRNA accumulation phenotypes.

### 2.7 Cluster properties distinguish extreme from normal FOVs

Having identified tissue regions with abnormal mRNA accumulation, we asked what distinguishes clusters in extreme FOVs from those in normal FOVs (fig. 5; figs. S20 to S21). Do extreme FOVs simply contain more clusters, or are individual clusters also different in size, density, or subcellular localization?

**Fig. 5:**
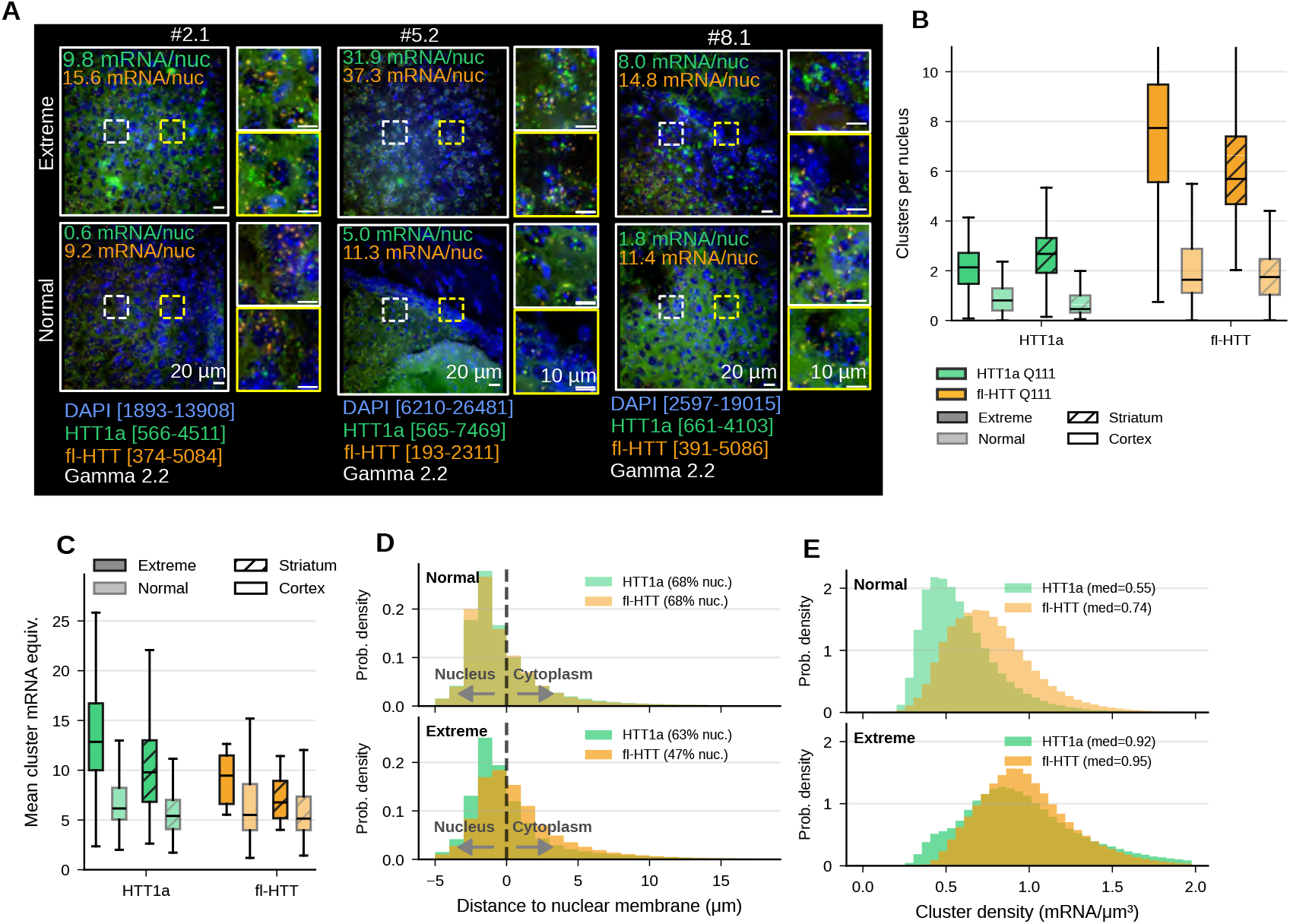
Cluster properties in extreme vs normal FOVs reveal mechanistic insights. **(A)** Representative FOV images comparing extreme (top row) vs normal (bottom row) Q111 expression across three mice (#2.1, #5.2, #8.1). Per-nucleus mRNA values annotated (green = HTT1a, orange = fl-HTT); insets show higher magnification of clusters. DAPI nuclear stain in blue. Images displayed with gamma correction (*γ* = 2.2; 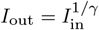) to enable simultaneous visualization of dim single molecules and bright clusters; the intensity scale is therefore non-linear, and bright spots are substantially brighter in the raw data than they appear. Dynamic ranges (raw image intensity values, not photon-corrected) shown at bottom of each column. Scale bars: 20 *µ*m (main), 10 *µ*m (insets). **(B)** Clusters per nucleus. Box plots: extreme (dark, left) vs normal (light, right) Q111 FOVs. Sample sizes: HTT1a cortex normal n = 480, extreme n = 217; striatum normal n = 238, extreme n = 399; fl-HTT cortex normal n = 585, extreme n = 112; striatum normal n = 359, extreme n = 278. Mann-Whitney U test: HTT1a cortex extreme median = 2.137, normal = 0.807, *p* = 2.86 × 10^−54^; striatum extreme = 2.678, normal = 0.458, *p* = 9.03 × 10^−91^. Cluster number is the primary driver of elevated mRNA. **(C)** Mean cluster size (mRNA equivalents). Extreme FOVs have larger clusters. **(D)** Cluster localization (distance to DAPI edge, *µ*m). Negative = nuclear; positive = cytoplasmic. Top: normal FOVs; bottom: extreme FOVs. Legend shows nuclear % (HTT1a: 67.9% normal, 63.0% extreme; fl-HTT: 67.9% normal, 47.5% extreme). **(E)** Cluster density (mRNA/*µ*m^3^). Top: normal; bottom: extreme. HTT1a median normal = 0.55, extreme = 0.92 (1.69×); fl-HTT median normal = 0.74, extreme = 0.95 (1.29×). Quality control: FOVs with <40 nuclei were excluded to ensure reliable per-nucleus normalization, removing 3,253 HTT1a clusters and 9,735 fl-HTT clusters relative to the unfiltered totals in fig. 2. Total clusters after filtering: HTT1a 637,067 (normal 216,482, extreme 420,585); fl-HTT 1,328,173 (normal 670,520, extreme 657,653). Extended analysis in figs. S20 to S21.

Extreme FOVs showed increased cluster abundance compared to normal FOVs (fig. 5B): for HTT1a, extreme FOVs had median 2.12 clusters/nucleus in cortex (IQR 1.26) vs 0.81 in normal FOVs (IQR 0.90), representing a 2.6× fold change (Mann-Whitney U: *p* = 7.47 × 10^−54^); striatum showed median 2.69 vs 0.47, a 5.7 × fold change (*p* = 1.58 × 10^−90^). For fl-HTT, cortex showed 4.7× increase (median 7.67 vs 1.64, *p* = 2.24 × 10^−55^); striatum 3.2× (5.69 vs 1.76, *p* = 2.80 × 10^−97^). This 2-6-fold increase in cluster number is the primary driver of elevated mRNA levels in extreme FOVs. Individual clusters in extreme FOVs also contained more mRNA (fig. 5C): mean cluster mRNA was 12.9 vs 6.1 mRNA equivalents in cortex (*p* = 6.02 × 10^−52^) and 9.8 vs 5.4 in striatum for HTT1a, indicating that both cluster number and mRNA content per cluster contribute to elevated expression.

Beyond abundance and mRNA content, clusters in extreme FOVs differed in their mRNA packing density (fig. 5E): median cluster density was 1.3-1.7-fold higher in extreme compared to normal FOVs (HTT1a: 0.92 vs 0.55 mRNA/*µ*m^3^, 1.69× fold change; fl-HTT: 0.95 vs 0.74 mRNA/*µ*m^3^, 1.29× fold change). This indicates that clusters in extreme FOVs are not simply bigger; they contain more mRNA per unit volume.

Clusters showed nuclear bias in both extreme and normal FOVs (fig. 5D), with 67-68% located within the DAPI-stained nuclear region in normal FOVs (median distance to DAPI: −1.05 to −1.12 *µ*m). However, extreme FOVs showed a significant shift toward cytoplasmic localization, particularly for fl-HTT: extreme FOVs showed 45-49% nuclear (median distance +0.12 to +0.34 *µ*m, i.e., cytoplasmic) vs 68% in normal FOVs (*p* = 1.69 × 10^−32^ cortex, *p* = 2.12 × 10^−34^ striatum). HTT1a showed a more modest shift (61-66% nuclear in extreme vs 67-75% in normal, *p* = 1.11 × 10^−5^ to 1.99 × 10^−14^).

### 2.8 Technical validation and control analyses

Control analyses validated that experimental signals represent genuine biological variation rather than technical artifacts. Negative control thresholds showed no significant correlations with experimental mRNA expression (Pearson |*r*| < 0.3, *p* > 0.3 for all channel-region combinations; fig. S7). Positive control analysis quantified housekeeping gene expression: POLR2A (low-abundance) showed 16 ± 15 mRNA/nucleus in cortex and 17 ± 20 in striatum (paired *t*-test: *t* = −0.91, *p* = 0.38), while UBC (high-abundance) showed 89 ± 48 and 177 ± 105 mRNA/nucleus respectively (*t* = −4.81, *p* = 0.0002) (fig. S8). POLR2A showed strong correlations with experimental HTT probes (cortex *r* = 0.81, striatum *r* = 0.98 for HTT1a; fig. S9). A potential concern is that these correlations could be spurious: tissues with better RNA preservation might show higher counts for all transcripts, creating apparent correlations between any gene pair. To investigate this, we performed partial correlation analysis, statistically removing the effect of UBC (used here as a proxy for tissue quality) from both POLR2A and HTT measurements. If the original correlations were driven by tissue quality, partial correlations would be substantially reduced. Instead, correlations remained virtually unchanged (HTT1a striatum: original *r* = 0.975, partial *r* = 0.977; <1% reduction; fig. S10). While this analysis has caveats (UBC may not perfectly capture all technical variation), the persistence of strong correlations after controlling for tissue quality is consistent with genuine biological coupling rather than a pure technical artifact. Comparison with bulk RNA-seq data from an independent Q111 mouse cohort (GSE65774 [10]) showed broadly consistent patterns: the unit-independent UBC/POLR2A ratio was comparable between methods (44 ± 68 RNAscope vs 54 ± 10 RNA-seq; *p* = 0.48), validating our absolute quantification (fig. S11). RNAscope showed higher variance (CV 154% vs 18% for the ratio), likely reflecting preserved cell-to-cell heterogeneity that bulk RNA-seq averages out [9]; sampling differences may also contribute, as our RNAscope approach examines thin tissue sections (40 *µ*m) from specific anatomical locations, while the RNA-seq dataset used dissected striatum of unspecified volume.

## 3 Discussion

We have developed and validated a high-throughput RNAscope imaging pipeline that quantifies both single mRNA molecules and their clusters in fixed brain tissue at single-molecule resolution. The key technical innovation is the establishment of a quantitative relationship between fluorescence intensity and absolute mRNA content, validated at two scales: for single molecules, the linear relationship between spot size and intensity (*r*^2^ > 0.90) is consistent with variable probe binding along the transcript; for clusters, the linear scaling between cluster volume and mRNA equivalents (*R*^2^ > 0.98) confirms uniform probe accessibility. Applied to HttQ111+/− knock-in mice, our analysis reveals several notable features of mutant huntingtin transcript biology: a substantial fraction of HTT mRNA exists in clusters rather than as single molecules, these clusters typically contain 4-5 transcripts, and considerable heterogeneity exists across tissue regions.

### 3.1 Technical advances: quantitative mRNA counting through validated linearity

Our pipeline addresses key limitations of standard RNAscope analysis by establishing objective, reproducible methods for quantifying both single mRNA molecules and multi-transcript clusters. The critical validation is the linear relationship between cluster volume and mRNA content. This linearity is essential for three reasons: (1) it confirms that our per-slide modal normalization correctly converts arbitrary intensity units to absolute mRNA counts; (2) it demonstrates that RNAscope probe amplification trees access mRNA uniformly throughout clusters, with no saturation at larger cluster sizes; and (3) it provides a quantitative foundation for comparing mRNA accumulation across samples, slides, and experimental conditions.

For single-molecule detection, we use empirical PSF calibration that accounts for the ∼57% larger effective spot dimensions arising from RNAscope probe cluster geometry compared to 200 nm fluorescent beads. The two-stage calibration (first establishing the size-intensity relationship on high-quality data to identify the breakpoint, then applying size-based filtering to isolate single molecules) provides an objective, data-driven approach to single-molecule identification that does not rely on arbitrary intensity thresholds.

For cluster quantification, we developed a neural network-based 3D segmentation approach combined with intensity normalization using the per-slide single-molecule reference. The two-stage cluster filtering (pruning overlaps with single molecules, then applying intensity and CV thresholds) prevents double-counting while removing background signals. The resulting intensity-volume relationship validates this approach: deviations from linearity would indicate probe penetration limitations, normalization errors, or systematic biases in cluster segmentation. The per-slide normalization strategy is essential given the 2-3× variation in raw photon counts across slides (fig. 1E, inset). By identifying the modal intensity (the most common signal, corresponding to single mRNA molecules) via KDE and scaling each slide such that this mode equals *N*_1mRNA_ = 1.0, we center the intensity distributions around 1.0 (fig. 1E, main panel). After applying size-based filtering to isolate single molecules, the distributions sharpen further around *N*_1mRNA_ = 1.0 (fig. 1H), confirming that the filtering step successfully isolates a homogeneous population of single-molecule signals.

This quantitative intensity approach explicitly departs from manufacturer guidance, which emphasizes dot counting over intensity measurement. Our departure is justified by empirical validation: the strong linear relationship between cluster volume and normalized intensity (*R*^2^ > 0.98) demonstrates that our calibration procedure successfully converts fluorescence intensity to absolute mRNA counts. Without this calibration framework, particularly the per-slide modal normalization and negative control thresholds, intensity-based quantification would indeed be unreliable due to the substantial slide-to-slide variation we observe (2-3× in raw photon counts). The manufacturer’s conservative guidance reflects the challenges of intensity quantification without proper calibration, which our pipeline addresses.

#### 3.1.1 Sources of variance and inference framework

Our measurements contain both technical and biological sources of variation. Technical variation arises from slide-to-slide differences in imaging parameters (2-3× variation in raw photon counts) and negative control thresholds (CV ≈ 40%). We address this through per-slide calibration: modal normalization removes photon-yield differences, and per-slide negative control thresholds account for local background conditions.

Biological variation occurs at multiple levels. Between animals, we observe substantial differences in mRNA accumulation (fig. 3F), with some Q111 mice showing expression levels comparable to wildtype while others exhibit dramatically elevated clustering. Within animals, different tissue regions show heterogeneous mRNA accumulation: the fraction of FOVs classified as “extreme” (exceeding wildtype P95 thresholds) varies by brain region and probe, ranging from 16% (fl-HTT in cortex) to 63% (HTT1a in striatum). Age effects trend upward but do not reach statistical significance with our current sample sizes.

Our study design samples at three nested levels: multiple FOVs per slide, multiple slides per mouse, and multiple mice per condition. For statistical comparisons, we average FOVs within each slide first, then compare slides across conditions. This avoids treating FOVs as independent observations (pseudoreplication) and ensures that the high FOV-to-FOV variability ( ∼30% CV) is appropriately averaged rather than inflating our apparent statistical power. The biological variance between animals, not technical noise between FOVs, is what limits our ability to detect genotype and age effects.

### 3.2 Biological insights: heterogeneous mRNA accumulation

A notable observation is not the prevalence of clustering per se, but rather the heterogeneous distribution of elevated clustering across tissue. Independent work using FISH has demonstrated widespread formation of mutant HTT mRNA clusters in HD mouse models, with 54-96% of striatal and cortical neurons containing clusters in YAC128 and BACHD-97Q-ΔN17 mice [26]; similar nuclear clustering was confirmed in YAC128 and BAC-CAG models [13]. Our FOV-level analysis reveals that 16-63% of Q111 FOVs exceed the 95th percentile of wildtype clustered mRNA, with considerable regional and individual variation. In striatum, 62.6% of Q111 FOVs showed extreme HTT1a clustering compared to only 31.1% in cortex; for fl-HTT, the corresponding values were 43.6% and 16.1%. This heterogeneity (where a substantial subset of tissue regions shows elevated mRNA accumulation) may be relevant to understanding disease pathogenesis and assessing therapeutic mechanisms, though larger sample sizes will be needed to confirm these patterns.

Although our per-slide analysis lacked statistical power to detect significant age effects (n = 4-6 slides per group), median mRNA expression values increased consistently with age across all conditions (e.g., Q111 striatum HTT1a: 5.4 at 2 months to 25.7 mRNA/nucleus at 12 months). This trend is consistent with the progressive pathology reported in Q111 mice, where neuronal intranuclear inclusions (NIIs) of mutant huntingtin protein increase from near-zero at 3 months to 28% of striatal neurons by 12 months, with inclusion size increasing over 40% between 9 and 12 months [17], and motor phenotypes emerge at 9 months and worsen progressively [18]. Future studies with larger sample sizes will be needed to determine whether the observed mRNA trends reach statistical significance. Our quantitative framework enables characterization of this heterogeneity. Extreme FOVs are characterized by: (1) elevated cluster numbers per nucleus (2-6× increase), indicating that cluster number is a primary driver of elevated total mRNA; (2) higher cluster density, i.e., more mRNA molecules packed per unit volume (1.3-1.7× higher); and (3) a shift toward cytoplasmic localization in extreme FOVs, particularly for fl-HTT (48% vs 68% nuclear). The interpretation of this cytoplasmic shift remains unclear. While mutant huntingtin has been shown to disrupt nuclear pore complex integrity [27], HD models typically show nuclear *retention* of mRNA rather than increased cytoplasmic accumulation [26]. The shift we observe could reflect altered nucleocytoplasmic transport dynamics, differences in cluster detection sensitivity between compartments, or other factors that will require further investigation. These specific metrics provide quantitative readouts that could potentially be used to assess therapeutic interventions targeting HTT expression or processing.

The mRNA clusters we characterize here are consistent with previous observations of mutant huntingtin mRNA forming nuclear clusters in HD mouse models, where these structures have been shown to exhibit phase-separation-like properties and resist therapeutic silencing [13, 26]. More broadly, nuclear mRNA foci containing repeat-expanded transcripts form through RNA-driven phase separation in a repeat length-dependent manner [46], and mutant huntingtin protein inclusions themselves behave as dynamic, phase-separated compartments [47]. Our 3D z-stack imaging enables direct volumetric density measurements (0.55-0.95 mRNA/*µ*m^3^), a metric not commonly reported in condensate studies. For example, Cochard et al. used smFISH with maximum intensity projections to quantify RNA recruitment to artificial protein condensates, reporting surface densities of 2-16 RNA/*µ*m^2^ and ∼25-32 total RNAs per condensate (0.8-2.0 *µ*m diameter) [48]. The total mRNA content per structure is in the same order of magnitude as our measurements (median 3.8-4.5, mean 6-9 mRNA equivalents per cluster), with the tail of our distribution extending beyond 20 mRNA equivalents (fig. 2E). However, converting their 2D surface density to volumetric density (by dividing total surface RNA by the sphere volume: 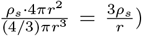 yields ∼ 6-120 RNA/*µ*m^3^, substantially higher than our values. This difference may reflect distinct biology (native disease-associated mRNA clusters vs. engineered protein condensates), but also highlights that volumetric density depends critically on how cluster boundaries are defined. Defining boundaries consistently is non-trivial: we use local background correction, which makes total mRNA counts robust to boundary placement (a slightly larger boundary still yields accurate mRNA counts because background signal is subtracted), but the volume is directly affected by boundary choice, and thus so is the density. Consistent methodology, as applied here, is essential for meaningful comparisons across conditions. The observation that clusters in extreme FOVs show higher mRNA packing den-sity could reflect concentration-dependent effects, where elevated local mRNA levels promote further condensation, though distinguishing active concentration mechanisms from passive crowding will require additional experiments. Several mechanisms could contribute to this heterogeneity. First, transcription from the mutant allele may be stochastically variable, with some cells experiencing transcriptional bursts that generate high local mRNA concentrations [5, 8]. Second, CAG repeat-containing transcripts form stable hairpin secondary structures that may impair degradation or export; in-cell studies show these hairpins remain largely folded under physiological conditions [28]. Critically, HTT1a arises from CAG repeat length-dependent incomplete splicing: the expanded repeat causes a block in splicing between exon 1 and exon 2, followed by cleavage and polyadenylation at a cryptic site in intron 1, producing a short mRNA encoding only exon 1 [5, 19]. The resulting exon 1 HTT protein fragment (containing the expanded polyglutamine tract) forms aggregates more readily than full-length protein and causes aggressive pathology; the R6/2 mouse, which expresses only this fragment, is the fastest-progressing HD model [20]. The uniform distribution of this pathogenic HTT1a species across cortex and striatum may be particularly relevant for understanding why striatal neurons (despite not having higher HTT1a levels) show selective vulnerability, supporting the hypothesis that intrinsic cellular factors rather than transcript abundance determine susceptibility [16].

Ideally, RNAscope imaging would be combined with immunofluorescence to assess protein localization and concentration in parallel with mRNA measurements. Such combined approaches would enable direct correlation between transcript accumulation and protein aggregation at the single-cell level, providing insight into whether mRNA clusters precede, coincide with, or follow protein inclusion formation. This would be particularly valuable for understanding the relationship between HTT1a mRNA clusters and the huntingtin protein inclusions that are hallmarks of HD pathology.

### 3.3 Implications for therapeutic development

The predominance of clustered mutant HTT mRNA raises important questions for ASO and siRNA therapeutics currently in clinical development for HD [11]. Recent studies have shown that huntingtin mRNA nuclear localization is cell-type specific, with approximately 50% of HTT mRNA localizing to the nucleus in neurons [12]. While ASOs can reduce both cytoplasmic and nuclear HTT mRNA (achieving ∼90% and ∼60% reduction respectively), siRNAs primarily eliminate cytoplasmic transcripts with minimal effect on nuclear mRNA [12]. Critically, nuclear mRNA clusters–which are specifically enriched in the aberrantly spliced HTT1a transcript [26]–are resistant to both siRNA and GapmeR ASO-based silencing [13], possibly explaining reduced efficacy in cases with ultralong CAG repeats where nuclear clusters are more prevalent. Our finding that the majority of mutant HTT signal exists in clusters and ∼48-68% of these are nuclear therefore presents a therapeutic challenge, particularly for HTT1a which preferentially forms these resistant nuclear clusters. If nuclear clusters are resistant to current approaches, measuring cluster reduction (in addition to total mRNA knockdown) may be important for predicting clinical efficacy.

### 3.4 Generalizability to other RNAscope applications

Several features of this pipeline should generalize beyond huntingtin mRNA to other RNAscope applications. The core principle, using isolated single molecules as an internal intensity standard for each slide, requires only that some fraction of target transcripts exist as resolvable single molecules, which is expected for most transcripts under most conditions. The key factors determining applicability include: (1) *Probe density* : our probes contain 20 ZZ pairs spanning ∼1000 bp target regions, providing sufficient fluorescence for reliable single-molecule detection; transcripts with shorter target regions or fewer probe pairs may require sensitivity adjustments. (2) *Transcript structure*: the extended nature of RNAscope amplification trees (reflected in the ∼57% larger PSF compared to point-like beads) may vary with target mRNA secondary structure; transcripts with extensive secondary structure might show different effective PSF dimensions. (3) *Clustering behavior* : our cluster quantification relies on the linear volume-intensity relationship, which requires uniform probe accessibility throughout clusters; transcripts forming dense phase-separated condensates with limited probe penetration might show saturation at large cluster sizes. (4) *Expression level* : transcripts expressed at very low levels may lack sufficient single molecules for reliable modal intensity estimation, while very high expression may cause extensive clustering that obscures single-molecule signals. For applications meeting these criteria, our calibration framework provides a validated approach to absolute mRNA quantification that extends RNAscope beyond simple dot counting.

### 3.5 Limitations and future directions

Several limitations should be noted. Our wildtype sample size (n = 2) was insufficient for robust genotype comparisons. A major limitation is that we segmented DAPI signal per FOV and normalized by mean nuclear volume rather than performing individual nuclear segmentation. While we used DARPP-32 staining to identify the striatal region containing MSNs, this information was not incorporated into per-cell analysis; our approach cannot differentiate individual MSNs from other cell types (interneurons, glial cells) within the DARPP-32-positive region, which would require individual nuclear segmentation combined with cell-type-specific markers. Cell-type-specific analysis through individual nuclear segmentation combined with morphological or molecular markers would greatly enhance the biological interpretation of our findings. While we quantified nuclear vs cytoplasmic localization, we did not resolve sub-nuclear compartments. Sample preparation requires improvement: of 30 slides, 12 (∼40%) were excluded due to poor tissue integrity, substantially reducing statistical power. Additionally, our slide selection did not systematically sample along the anterior-posterior axis, precluding rigorous analysis of spatial heterogeneity along this dimension. The substantial heterogeneity observed (both between animals and across tissue regions) suggests that sparse sampling may lead to unreliable estimates, motivating future studies to image large tissue areas per brain. Finally, our 3D Gaussian PSF model assumes a point-like emitter, but RNAscope probe binding regions span ∼1000 bp of the target sequence, creating a spatially extended fluorescent source; replacing the point-source assumption with a model that accounts for this finite emitter size would enable deconvolution-based estimation of actual emitter dimensions and improve single-transcript classification. For example, a vectorial PSF model [37–39], which can also account for non-optimal optics, could be convolved with geometric primitives representing the probe binding region–a sphere for isotropic binding, or a rod-like structure aligned along the transcript to better approximate the ∼1000 bp target sequence orientation. While individual FOV-level deviations can be large, we do not propagate this variability directly into mouse-level conclusions. When aggregating to the slide or mouse level (fig. 3), we observe consistent trends but lack statistical power with only 9 Q111 and 2 wildtype mice to detect significant differences. However, at the FOV level, significant differences emerge: Q111 FOVs show elevated mRNA accumulation compared to wildtype (fig. 4), and FOVs classified as extreme based on wildtype thresholds exhibit distinct cluster properties including elevated cluster numbers, higher packing density, and altered subcellular localization (fig. 5).

Future directions based on these limitations include: expanding sample sizes; implementing individual nuclear segmentation to enable cell-type-specific analysis; registering sections to a standardized mouse brain atlas; processing whole brains to capture full spatial heterogeneity; developing PSF models that incorporate finite emitter size; and applying the pipeline to evaluate therapeutic interventions using the cluster metrics identified here.

### 3.6 Conclusion

We present a high-throughput RNAscope imaging and analysis pipeline for quantifying mutant huntingtin mRNA in mouse brain at single-molecule resolution. The pipeline’s key innovation is the establishment and validation of a linear relationship between cluster volume and mRNA content, enabling quantitative conversion of fluorescence intensity to absolute mRNA counts. Applied to HttQ111+/− mice, we observe considerable heterogeneity in mRNA accumulation: 16-63% of Q111 tissue regions show elevated clustering exceeding wildtype thresholds (HTT1a: 31% cortex, 63% striatum; fl-HTT: 16% cortex, 44% striatum). Extreme FOVs are characterized by elevated cluster numbers per nucleus (2-6× increase), higher cluster density (1.3-1.7×), and a shift toward cytoplasmic localization for fl-HTT (48% nuclear vs 68% in normal FOVs). While larger sample sizes will be needed to confirm these patterns, these quantitative metrics provide a framework for investigating disease mechanisms and evaluating therapeutic interventions. Importantly, having a validated analysis pipeline now justifies the investment in RNAscope probes and large-scale imaging: systematic studies across disease models, timepoints, and therapeutic interventions are now feasible. Combined with improvements in sample preparation to reduce tissue loss, this approach enables quantitative, high-throughput mRNA profiling for HD and other repeat expansion disorders characterized by RNA foci formation.

## 4 Materials and Methods

### 4.1 Samples and Treatment

#### 4.1.1 Mouse Models

Brain tissue samples were obtained as described previously [50]. All mouse experiments were performed in alignment with the Institutional Animal Care and Use Committee at the University of Massachusetts Chan Medical School (protocol 202000010). Mice were bred and born at Jackson Labs (Farmington, CT) and transferred at 6 weeks of age to pathogen-free UMass Chan animal facilities, where they were housed with a 12h light:12h dark cycle at 23 ± 1^°^C and 50 ± 20% humidity with free access to food and water. We used heterozygous HttQ111+/− mice (JAX Strain ID 370624, Mc Q111 KI; C57BL/6J background) and wild-type C57BL/6J littermate controls. Tissue was collected after perfusion/fixation and processed identically across conditions to minimize batch effects. Mice in this study were derived from a separate therapeutic study investigating siRNA-mediated silencing of MSH3 and HTT [50]. That study administered treatments via intracerebroventricular (ICV) injection at 2 months of age. The mice used here were from control groups only: 2-month-old mice were untreated (UNT, prior to injection), while 6- and 12-month-old mice received either artificial cerebrospinal fluid (aCSF) or a non-targeting control siRNA (NTC). As these are standard control conditions not expected to alter HTT mRNA expression, we pooled all Q111 mice for analysis.

#### 4.1.2 Tissue Processing

After transcardial perfusion and fixation, brains were sectioned coronally at 40 *µ*m thickness using a vibratome. Sections were stored free-floating in PBS at 4^°^C prior to RNAscope staining.

##### Mouse ID Mapping

Anonymized mouse IDs used in Figures 3F and 4F correspond to the following samples (Table 1):

#### 4.1.3 Staining Protocol

RNAscope™ assays were performed according to the manufacturer’s instructions (ACD Bio; detailed protocol in Supplementary Note 3, reagents in table S6). Typical probe sets comprised up to 20 ZZ pairs per target; one ZZ pair spans ∼50 bases, yielding a tiled probe region of ∼1000 bases and an estimated physical contour length of ∼260 nm (assuming ∼0.26 nm per base). Because this effective label extent approaches the lateral PSF, we acquired additional PSF calibration data with larger beads (see Section 4.3.3). For housekeeping control signals we used a ready-to-use mixture provided by the manufacturer containing three probes targeting POLR2A (channel C1), PPIB (channel C2), and UBC (channel C3).

#### 4.1.4 Slide Preparation

After staining and amplification, samples were counterstained with DAPI and mounted in ProLong− Glass Antifade Mountant (Thermo Fisher Scientific, P36982; refractive index 1.52 after curing) with a No. 1.5 high-precision coverslip (Thorlabs, CG15KH1; 170 *µ*m ±5 *µ*m thickness). Slides were cured flat at room temperature (30-60 min) and stored protected from light at 4^°^C until imaging.

### 4.2 Optical Setup & Image Acquisition

Images were acquired sequentially on a TissueFAXS SL Q tissue cytometer (TissueGnostics, TG3-970 C), illuminated by a Lumencor Spectra III solid-state light engine with nominal excitation bands at 390/22 nm (mDAPI), 475/28 nm (sFITC), 555/28 nm (sCY3), and 637/12 nm (sCY5).

Two objectives were used:

- 2.5× (NA 0.085; Zeiss EC Plan-Neofluar M27) for whole-slide overview and ROI selection (2D).
- 40× oil (NA 1.3; Zeiss Plan-Apochromat Oil DIC (UV) VIS-IR) for 3D confocal imaging.

Following 2.5× overview scans, immersion oil (Immersol™ 518F; refractive index 1.518 at 23^°^C) was applied before 40× confocal acquisitions. Emission was split using a SpectraSplit^®^ 7 filter set (S-Split Blue, Green, Orange, Farred), with central excitation/emission wavelengths (nm) of 374/425, 491/526, 543/570, and 648/688, respectively. Fluorescence was detected on a Hamamatsu ORCA-Fusion BT sCMOS camera (C15440-20UP) with a physical pixel size of 6.5 *µ*m. At 40× magnification, this yields an effective pixel size in the image plane of 6.5 *µ*m*/*40 = 162.5 nm. Confocal sectioning used a Crest V2 spinning disk (15,000 rpm; pinhole diameter 60 *µ*m, spacing 250 *µ*m). All other optics (mirrors, lenses, collimation) were standard to the TissueFAXS SL Q system.

Acquisitions were controlled with TissueFAXS SL Viewer v7.1.131. Overview scans used exposure and power settings optimized for contrast. For confocal stacks, laser power and exposure were set such that the brightest signals remained below 40,000 ADUs; per-sample settings are listed in Supplementary Table S1. Laser powers were measured in situ (Thorlabs S170C; table S2). Illumination was restricted by a circular iris to a 485 *µ*m area centered in the FOV. Each z-stack comprised 21 slices acquired at 500 nm axial spacing, providing a total imaging depth of 10 *µ*m.

The Airy unit (AU) at Alexa 546 emission is 1 AU = 1.22 *λ*_573_*/*NA ≈ 538 nm. With system magnification, the 60 *µ*m pinhole corresponds to ∼2.8 AU, maximizing signal collection at the cost of thicker optical sections. The confocal optical slice thickness is

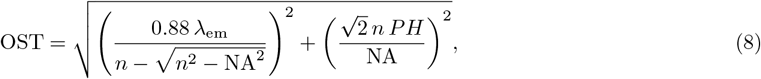

with pinhole diameter *PH* and refractive index *n* [33]. For Alexa 546 in our system, OST ≈ 2.6 *µ*m.

#### 4.2.1 Image photon calibration

We recorded 1000 dark frames in complete darkness to estimate the camera offset: 101 ± 6 ADU (mean± SD across pixels). Photon calibration was performed by acquiring 1000 out-of-focus brightfield images of fixed cells at three illumination levels spanning the 16-bit range. The response was linear up to ∼50,000 ADUs. The conversion gain (electrons/ADU) was obtained from a mean-variance fit [34], yielding 0.21 electrons/ADU (typical value ∼ 0.24 for this sensor [35]). All analysis used images converted to photon (electron) counts.

### 4.3 Imaging Model

We model the recorded intensity as the sum of an emitter signal convolved with the microscope PSF and a per-slice background term.

#### 4.3.1 Three-Dimensional Gaussian PSF Model

We model the emitter as a separable integrated 3D Gaussian, which provides a reasonable approximation for spinning-disk confocal near focus [21]:

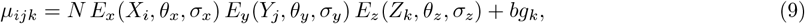

where indices *i, j* denote pixel positions in the lateral plane and *k* denotes the slice index, with parameters ***θ*** = [*θ*_*x*_, *θ*_*y*_, *θ*_*z*_, *N, bg*_1..*K*_, *σ*_*x*_, *σ*_*y*_, *σ*_*z*_]^*T*^ and *bg*_*k*_ is the background level for slice *k*. The per-axis integrated Gaussians are

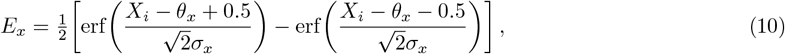

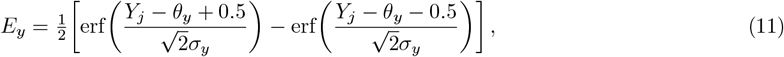

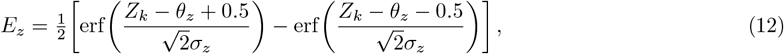

analogous to the 2D integrated model [36].

Closed-form first derivatives are used for gradient and Hessian computations. For example,

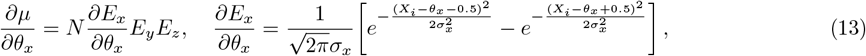

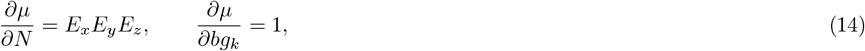

with analogous expressions for *θ*_*y*_, *θ*_*z*_ and *σ*_*x,y,z*_.

#### 4.3.2 Optimization Procedure (MLE using Levenberg-Marquardt)

Parameters are estimated by Poisson MLE. Given observed intensities *I*_*ijk*_ and predictions *µ*_*ijk*_,

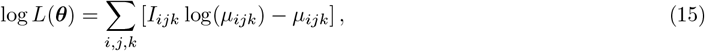

(up to an additive constant). The gradient is

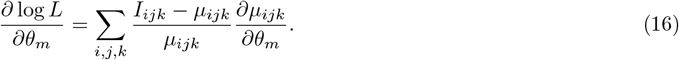

We use the Gauss-Newton approximation to the Hessian [40]:

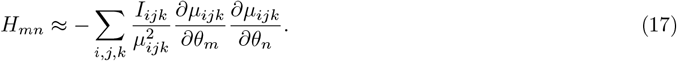

Levenberg-Marquardt updates are

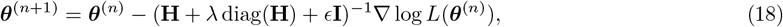

with *λ* = 10^−2^ and *ϵ* = 10^−6^. In practice, we compute the pseudo-inverse via torch.linalg.pinv [41] to ensure robustness during batch processing: if any matrix in the batch is singular, the pseudo-inverse returns a valid result rather than causing the entire batch to fail. After each step we clip parameters to physically plausible bounds: *θ*_*x,y*_ ∈ [2, 10] pixels; *θ*_*z*_ ∈ [0, 30] slices; *N* ∈ [1, 10^9^] photons; *bg*_*k*_ ∈ [1, 10^6^] photons; *σ*_*x,y*_ ∈ [0.2, 4] pixels; and *σ*_*z*_ ∈ [0.2, 10] slices. Convergence is declared if all absolute parameter changes fall below 0.01 nm or 0.01 photons, or after 60 iterations. Fits violating bounds or failing to converge within 60 iterations are discarded.

#### 4.3.3 PSF Calibration on Beads

We calibrated (*σ*_*x*_, *σ*_*y*_, *σ*_*z*_) on TetraSpeck− Fluorescent Microspheres Size Kit (Invitrogen−; 0.2 *µ*m) with emission labels at 365/430 (blue), 505/515 (green), 560/580 (orange), and 660/680 nm (dark red). Beads are pre-mounted at the coverslip in ProLong− Glass (refractive index 1.52; per manufacturer support for slides produced 2021-2025), matching the mounting conditions used for tissue samples. Across 20 fields of view per channel, we extracted > 6,000 bead volumes per channel (range 6,330-142,365) and fit 3D Gaussians following the procedure described in Section 4.3.2. The resulting *σ*_*x*_, *σ*_*y*_, and *σ*_*z*_ distributions were then fit with Gaussian functions (SciPy curve fit [43]) to obtain mean PSF widths and standard errors. These values are summarized in Supplementary Table S3; the calibrated widths are used as priors and quality bounds in tissue fits.

### 4.4 Detection of Single Spots

We detect and validate single-molecule signals via a permissive candidate step followed by statistical and biological filtering.

#### 4.4.1 Initial Detection

Candidates are detected on a maximum-intensity projection (MIP) of the z-stack using a difference-of-uniform-filters pipeline [24]: (i) apply uniform filters of width 6 and 12 pixels to the MIP and difference them; (ii) find local maxima exceeding 5 photons; (iii) enforce a 10-pixel minimum separation. The MIP provides robust 2D localization of candidates regardless of their axial position. For each detected candidate, we then extract a 12 × 12 pixel lateral ROI and a 10-slice axial sub-volume centered on the lateral center-of-mass for subsequent 3D Gaussian PSF fitting (Section 4.3.2).

#### 4.4.2 Detector False Positive Probability

After fitting, we compute a GLRT statistic between background-only (*H*_0_) and signal-plus-background (*H*_1_) [23]:

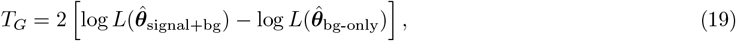

evaluated on each of 5 central slices using a 2D Gaussian PSF, as the 3D Gaussian model becomes less accurate further from focus [21]. P-values under *H*_0_ are

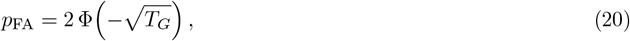

with Benjamini-Yekutieli FDR control [42]. Candidates with any slice *p*_FA_ > 0.05 are rejected.

#### 4.4.3 Biological False Positive Probability

Negative-control slides (bacterial probes) were imaged each session to profile non-specific binding and free-dye fluorescence. For each channel we computed the 95th percentile of negative-control photon counts,

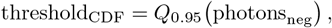

and accepted only candidates exceeding this threshold (per channel), complementing the detector-level filter and avoiding arbitrary cutoffs.

### 4.5 PSF Calibration Using RNAscope Data

#### 4.5.1 Rationale

Bead-derived PSFs may under-estimate the effective width of RNAscope signals. Unlike point-source fluorescent beads, RNAscope amplification trees consist of branched DNA structures with fluorophores distributed across the amplification tree, spanning approximately 20 ZZ probe pairs (∼ 50 base pairs each) that bind along the target mRNA. This spatial distribution of fluorophores effectively broadens the apparent PSF beyond the optical diffraction limit. We therefore refined the PSF constraints using tissue data directly.

#### 4.5.2 Two-Pass Calibration Procedure

We employed a two-pass approach to determine both the experimental PSF and the channel-specific breakpoints that distinguish diffraction-limited single-molecule signals from larger clusters (see fig. 1 for an overview):

##### Pass 1: Breakpoint determination with bead priors

1. **Initial detection and fitting**. Using bead-derived PSF widths as initial values, we performed MLE fitting on all detected candidates with *σ* bounds set to [0.2, 4] pixels laterally and [0.2, 10] slices axially, allowing the fitted *σ* to vary freely within these bounds.
2. **Signal-width analysis**. We binned fitted spots by their recovered *σ*_*x*_ values and computed the mean integrated photon signal *N* within each bin. For true single molecules with consistent fluorophore stoichiometry, *N* should remain approximately constant regardless of fitted *σ* (with variation due to noise). Instead, we observed approximately linear increases in *N* with increasing *σ* up to a channel-specific inflection point, beyond which the relationship breaks down.
3. **Breakpoint identification**. The breakpoint *σ*_bp_ was defined as the *σ* value where the signal-width curve transitions from linear growth to plateau or deterioration. Below the breakpoint, larger fitted *σ* values reflect noise or minor optical aberrations in true single molecules; above the breakpoint, large *σ* values indicate spatially extended signals (clusters) where the Gaussian model is inappropriate. We identified the breakpoint using segmented linear regression (piecewise linear fits with one knot).
4. **Experimental PSF estimation**. To determine the effective RNAscope PSF width for each channel, we fitted a parabola to the five histogram bins surrounding the breakpoint and extracted the vertex. This represents the most probable *σ* for single-molecule signals in tissue, accounting for RNAscope amplification tree geometry.

##### Pass 2: Final detection with experimental PSF

1. **Re-detection with calibrated priors**. We repeated the full detection pipeline (Section 4.4) using the experimentally determined PSF widths as initial values for MLE fitting, while still allowing *σ* to vary within bounds.
2. **Size-based rejection**. Fits were rejected from the single-molecule pipeline if any *σ* component fell outside the acceptable range: a minimum threshold of 80% of the bead-derived PSF width (to reject spuriously small fits likely caused by noise or fitting artifacts), and a maximum threshold at the channel-specific breakpoint (to reject spatially extended signals). Formally, fits were accepted only if 0.8 *σ*_bead_ ≤*σ* ≤*σ*_bp_ for each axis. Signals exceeding the upper breakpoint are captured by the CNN-based cluster detection (Section 4.6).

#### 4.5.3 Signal Preservation Through Overlapping Pipelines

Importantly, the cluster detection pipeline is intentionally designed to overlap with single-molecule detection at the boundary. Clusters that spatially overlap with validated single spots are pruned from the cluster masks (16.7% of green clusters, 26.6% of orange clusters; see fig. 2), ensuring no double-counting while preserving all detected signal. This overlap guarantees that signals near the single-molecule/cluster boundary are captured by at least one pipeline, preventing signal loss at the transition.

Recalibrated breakpoint values are listed in Supplementary Table S4.

### 4.6 mRNA Cluster Detection

#### 4.6.1 Training Data

We annotated clusters with LABKIT [25] in volumes of 512 × 512 × 21 voxels for Alexa 488 and Alexa 546. Signals visually larger than diffraction-limited spots were labeled as clusters; ambiguous cases were included to ensure comprehensive detection (with later pruning to protect single-spot counts).

#### 4.6.2 Inference and Cluster Definition

A 3D CNN [45] produced voxelwise cluster probability maps (architecture in Supplementary Section 1). Binary masks were obtained at *p* > 0.5. To avoid double counting, cluster masks overlapping validated single spots were removed. Diffraction-limited signals (single or merged sub-diffraction emitters) remained in the single-spot pipeline (Sections 4.3.1-4.3.2).

#### 4.6.3 Signal Estimation

For each cluster, we computed background-corrected photon counts on the photon-converted images:

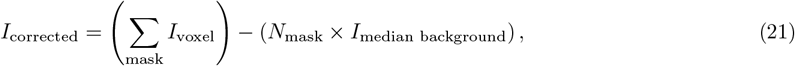

where *I*_corrected_ is the background-corrected photon count for the cluster, *I*_voxel_ is the photon count at each voxel within the cluster mask, *N*_mask_ is the number of voxels in the cluster mask, and *I*_median background_ is the median photon count in a local background shell formed by dilating the mask by 10 voxels in all dimensions and subtracting the original mask. Cluster centers-of-mass were retained for spatial analyses.

### 4.7 Fluorescent Signal with RNAscope

The RNAscope™ technique employs probes designed to target mRNA transcripts, with up to 20 individual probe pairs per mRNA molecule. Following probe hybridization, two sequential amplification steps can generate up to 8000 fluorescent labels per transcript [1]. Each probe pair requires two adjacent binding events on the target mRNA, significantly increasing specificity. However, variability introduced during the amplification process can result in significant fluctuations in fluorescence intensity per transcript. Consequently, careful consideration of how estimated signal intensities are interpreted and categorized is essential for accurate quantification.

Two custom probes were used to detect mutant huntingtin transcripts: (1) the HTT1a probe (Mm-Htt-intron1-O2, Cat. No. 575581) targeting intron 1, which detects the aberrantly spliced transcript; and (2) the fl-HTT probe (Mm-Htt-No-XHs, Cat. No. 473001) targeting the 3’ UTR (bp 9833-10798), which detects the canonical full-length transcript. Each probe consists of 20 ZZ pairs, with each ZZ pair covering approximately 40-50 nt, spanning roughly 1,000 nt of the target region. Full target sequences are provided in Supplementary Note 4 and table S5.

### 4.8 DAPI-based Nuclear Segmentation

DAPI-stained nuclei were segmented in 3D using a residual 3D U-Net architecture [45], trained on manually annotated brain tissue volumes. Labeled regions were filtered by size (minimum 5,000 voxels, maximum 500,000 voxels) to remove noise and artifacts.

### 4.9 Distance to Nuclear Border

We computed the signed Euclidean distance from each cluster center-of-mass to the nearest nuclear boundary using anisotropic distance transform. Negative values indicate nuclear localization; positive values indicate cytoplasmic localization.

### 4.10 Atlas Registration and Coordinate Assignment

Anterior-posterior coordinates were assigned to each tissue section as a whole (not independently for striatum and cortex) by manual visual comparison with the Allen Mouse Brain Atlas [32]. For each slide, a trained observer compared the 2.5× overview image to coronal reference images from the Allen Reference Atlas (available at https://mouse.brain-map.org), identifying the best-matching atlas plane based on anatomical landmarks including corpus callosum shape, striatal boundaries, and cortical layering.

Within each section, FOV positions were assigned to striatal or cortical regions based on DARPP-32 signal (for striatum) or anatomical landmarks (for cortex). Striatal regions were definitively identified by positive DARPP-32 staining marking medium spiny neurons. Cortical region boundaries were more arbitrary, lacking cell-type-specific markers; cortical FOVs were positioned to sample all cortical layers in approximately equal proportions.

Coordinates are reported following the Allen atlas convention, where reference sections are spaced at 100 *µ*m intervals. However, this visual matching procedure introduces substantial uncertainty: precise registration to the atlas coordinate system is not possible without computational image registration or fiducial markers. We estimate the uncertainty in anterior-posterior assignment to be on the order of ±200-400 *µ*m (2-4 atlas sections), depending on how distinctively the anatomical landmarks differ between adjacent reference planes. Coordinates should therefore be interpreted as approximate rostrocaudal positions rather than precise atlas locations.

The anatomical schematic showing cortical and striatal subregion labels (Fig. 3A) was adapted from the Allen Reference Atlas coronal sections.

### 4.11 Data and Code Availability

Analysis scripts were executed per slide in parallel on a high-performance computing (HPC) cluster. All analysis code is publicly available on GitHub at https://github.com/pvanvelde/rna_scope_HTT. The complete imaging dataset totals multiple terabytes. A representative subset of processed data is available at https://surfdrive.surf.nl/s/DJ8jHFSBxxRAj34. The complete dataset is available upon reasonable request.

## Acknowledgements

We thank the Sanderson Center for Optical Experimentation (SCOPE) at UMass Chan Medical School for imaging support, the Scientific Computing for Innovation (SCI) Cluster for high-performance computing resources, and the Department of Animal Medicine for mouse housing and care. We also thank the laboratory members for technical assistance and helpful discussions.

## Author contributions

P.v.V. conceptualized the study, developed the analysis software and methodology, performed formal analysis, created visualizations, and wrote the original draft. B.T. performed RNAscope experiments and imaging. S.A. performed RNAscope experiments and animal work and contributed to methodology. E.L. performed RNAscope experiments and animal work. R.F. provided feedback and contributed to discussions. A.S. performed animal work. J.B. supervised experiments and performed animal work. E.K. performed RNAscope experiments, animal work, and contributed to methodology. A.K. conceptualized the study, provided supervision, and acquired funding. D.G. conceptualized the study, provided supervision, acquired funding, and reviewed and edited the manuscript. All authors reviewed and approved the final manuscript.

## Supplementary Information

### Supplementary Tables

**Table S1:** Imaging parameters by session. Laser intensity (%) and exposure time (ms) settings for each fluorescence channel across imaging sessions. FITC (488 nm) was used for HTT1a detection; Cy3 (548 nm) was used for fl-HTT detection. The scanner is operated by imaging core staff, with settings chosen by the operator based on sample appearance. Parameter variations between sessions may partially contribute to the 2-3× variation in raw photon counts across slides, motivating per-slide intensity normalization.

| Session | DAPI |  | FITC (HTT1a) |  | Cy3 (fl-HTT) |  | Cy5 |  |
| --- | --- | --- | --- | --- | --- | --- | --- | --- |
|  | Int. (%) | Exp. (ms) | Int. (%) | Exp. (ms) | Int. (%) | Exp. (ms) | Int. (%) | Exp. (ms) |
| 1 | 50 | 150 | 50 | 500 | 50 | 50 | 50 | 100 |
| 2 | 50 | 150 | 70 | 500 | 50 | 150 | 50 | 500 |
| 3 | 50 | 100 | 80 | 500 | 50 | 100 | 50 | 100 |

**Table S2:**
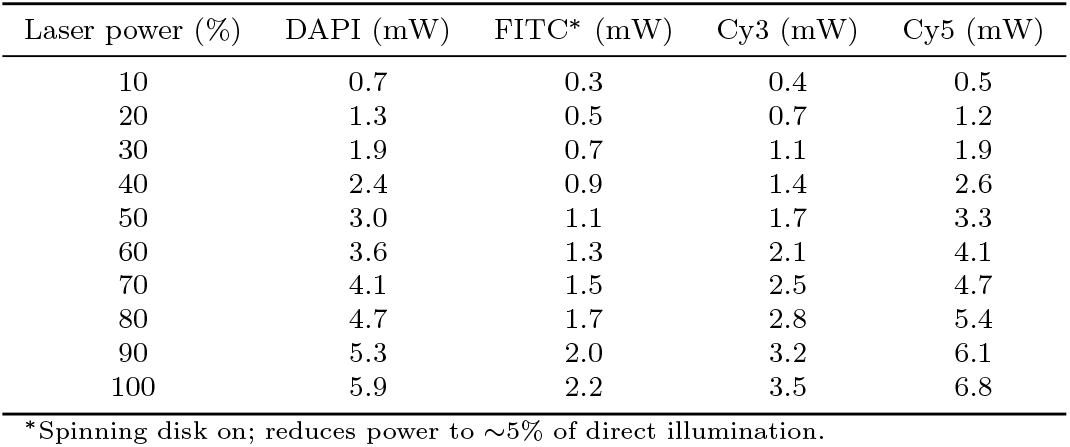
Laser power calibration. Measured laser power (mW) at the sample plane as a function of software-controlled laser intensity (%) for each fluorescence channel. Measurements performed on TissueFAXS SL Q tissue cytometer (TissueGnostics, TG3-970 C) with Plan-Apochromat 40×/1.3 Oil DIC objective. Power measured using Thorlabs S170C photodiode sensor (S/N: 7010096) with Zeiss Immersol 518 F immersion oil. Illumination area restricted by circular iris to 485 *µ*m diameter, centered at FOV center. All imaging data were acquired with the spinning disk active. FITC power was recorded with spinning disk on, showing ∼5% transmission relative to direct illumination. Other channels exhibited similar reduction with spinning disk but were measured without it. Power measurements were performed once during the imaging period (∼3 months); long-term stability and measurement uncertainty were not characterized.

**Table S3:**
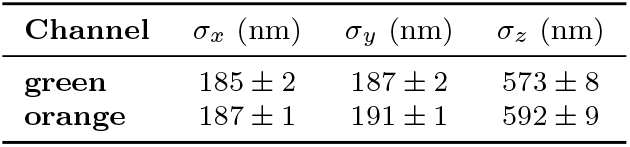
Bead-based PSF calibration. Mean *±* SE of *σ*_*x,y,z*_ per channel from 0.2 *µ*m TetraSpeck microspheres.

| Channel | $\sigma_x$ (nm) | $\sigma_y$ (nm) | $\sigma_z$ (nm) |
| --- | --- | --- | --- |
| green | 185 ± 2 | 187 ± 2 | 573 ± 8 |
| orange | 187 ± 1 | 191 ± 1 | 592 ± 9 |

**Table S4:** Empirically-determined breakpoints. Size thresholds separating single-molecule regime from cluster regime (see figs. S2 and S3).

| Channel | $\sigma_{x,\text{break}}$ (nm) | $\sigma_{y,\text{break}}$ (nm) | $\sigma_{z,\text{break}}$ (nm) |
| --- | --- | --- | --- |
| green | 290.8 | 291.1 | 698.7 |
| orange | 285.1 | 285.3 | 700.7 |

**Table S5:** RNAscope probe target specifications. Target region information for custom RNAscope probes used in this study. The HTT1a probe targets the 5’ region of intron 1 of mouse *Htt*, which is retained in the aberrantly spliced Htt1a transcript. This region contains the cryptic polyadenylation sites used in incomplete splicing of mutant huntingtin mRNA. The fl-HTT probe targets the 3’ UTR to detect the canonical full-length transcript. Full target sequences for both probes are provided in Supplementary Note 4 to document the exact genomic coordinates used for probe design, as reference genome annotations may change over time.

| Probe | Target Region | ZZ Pairs | Notes |
| --- | --- | --- | --- |
| Mm-Htt-intron1-O2 (HTT1a) | Mouse *Htt* intron 1 (bp 41-1190, 1,150 bp) | 20 | Detects aberrantly spliced Htt1a; target sequence in Supplementary Note 4 |
| Mm-Htt-No-XHs (fl-HTT) | Mouse *Htt* 3' UTR (bp 9833-10798, 966 bp) | 20 | Detects canonical full-length transcript; target sequence in Supplementary Note 4 |

### Supplementary Figures

**Fig. S1:**
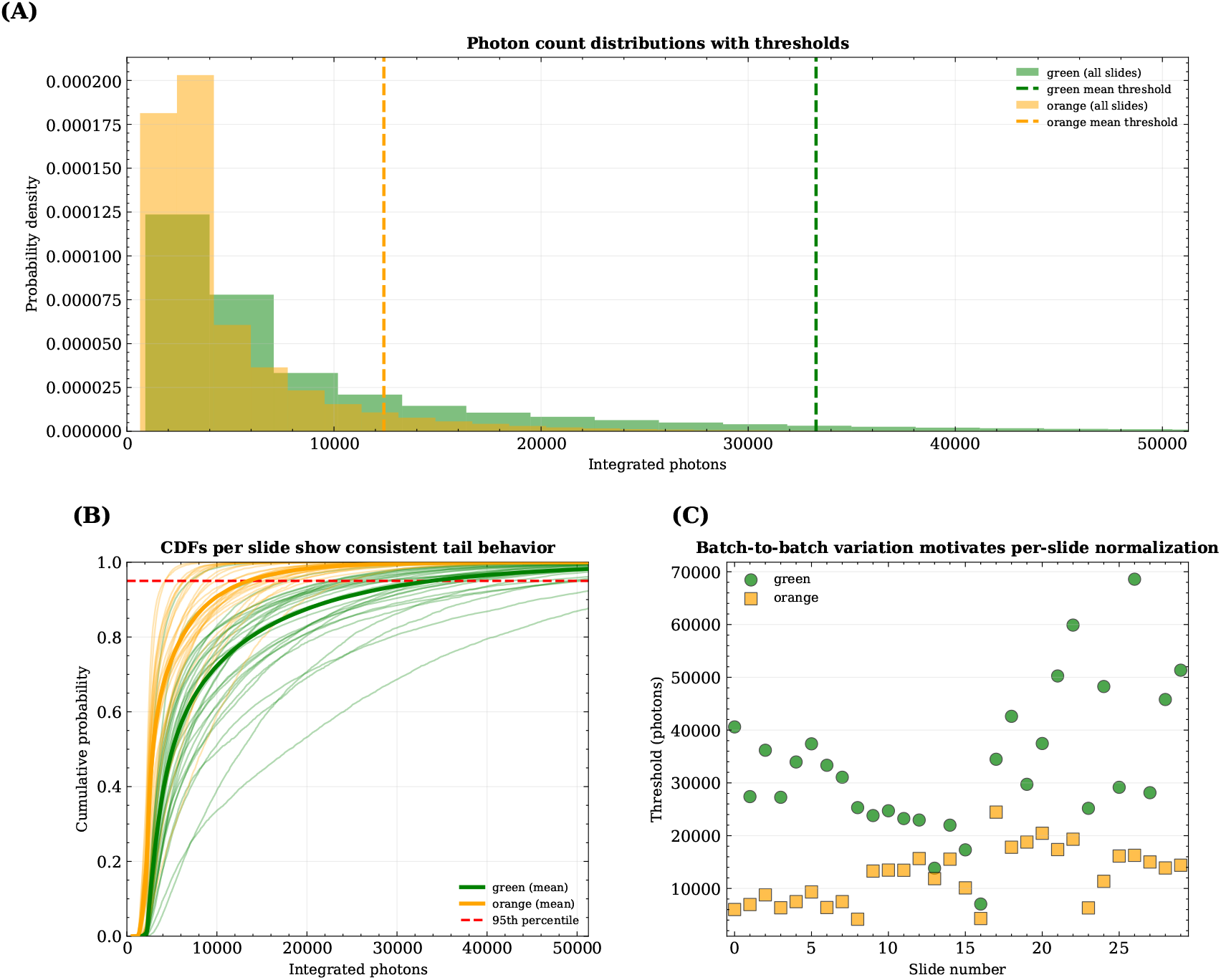
Negative controls establish per-slide detection thresholds. Analysis of 727,329 negative control spots (bacterial *DapB* probe, absent in mammalian tissue) across 30 slides (mean ± SD: 24,244 ± 7,748 spots/slide; 13,546 ± 7,748 green, 10,698 ± 6,113 orange). **(A)** Placeholder for example FOVs from negative-control sections showing sparse false-positive detections from autofluorescence and non-specific binding. **(B)** Distributions of integrated photon counts from all negative-control detections (combined across slides). Mean ± SD of 95th percentile thresholds: 33,266 ± 13,253 photons (green); 12,305 ± 5,211 photons (orange). **(C)** Cumulative distribution functions (CDFs) per slide (thin lines) and mean CDF across all slides (bold) demonstrate consistent tail behavior despite variation in median background. Horizontal line at 0.95 marks the quantile used for thresholding. **(D)** Threshold values versus slide number (circles: green channel, squares: orange channel) reveal substantial slide-to-slide variation: CV_green_ = 39.8% (33,266 ± 13,253 photons); CV_orange_ = 42.4% (12,305 ± 5,211 photons). This motivates per-slide normalization to account for batch-to-batch variation in autofluorescence and staining efficiency. All values reported as mean ± SD across slides.

**Fig. S2:**
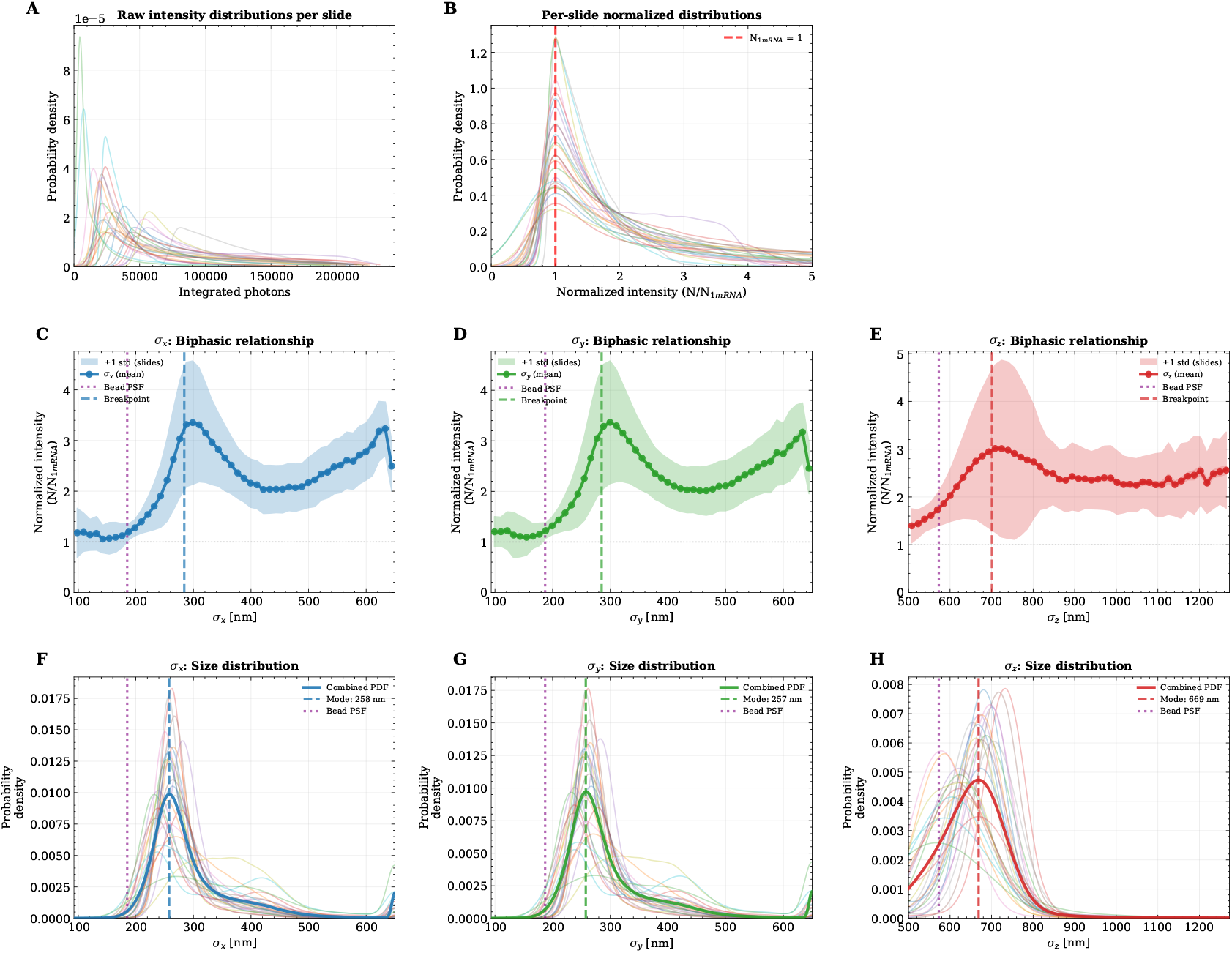
Breakpoint analysis for green channel (HTT1a). Analysis of 1,822,764 spots across 26 slides from Q111 mice (PFA < 1e-04). Median spots per slide: 70,106. **(A)** Raw intensity distributions per slide before normalization, showing 2-3× slide-to-slide variation in photon yield due to technical factors (tissue autofluorescence, fixation quality, probe hybridization efficiency, imaging conditions). **(B)** Per-slide normalized intensity distributions converging around *N*_1mRNA_ = 1 after modal normalization via KDE (Scott’s rule bandwidth [43, 44]). Successful removal of technical variation while preserving biological signal (cluster tail). **(C-E)** Size-intensity relationships for *σ*_*x*_, *σ*_*y*_, *σ*_*z*_. Bold line: mean from all pooled spots; light shaded region: ±1 SD across per-slide means; narrower dark bands: ±1 SEM. Purple dotted line: bead PSF (*σ*_*x*_ = 185 nm, *σ*_*y*_ = 187 nm, *σ*_*z*_ = 573 nm). Colored dashed line: empirically-determined breakpoint (*σ*_*x*_ = 284.1 nm, 53.6% larger than bead PSF; *σ*_*y*_ = 284.9 nm, 52.4% larger; *σ*_*z*_ = 700.8 nm, 22.3% larger). Piecewise linear regression: *σ*_*x*_ slope1 = 0.027 (single-molecule regime), slope2 = −0.005 (cluster regime). **(F-H)** Spot size distributions. Faint lines: per-slide KDE; thick line: combined PDF. Mode values from combined distributions (colored dashed): *σ*_*x*_ = 257.9 nm (39.4% larger than bead PSF), *σ*_*y*_ = 257.4 nm (37.7%), *σ*_*z*_ = 668.6 nm (16.7%). Size statistics: *σ*_*x*_ mean = 299.6 nm, median = 273.1 nm, SD = 90.9 nm. Modes define tissue-calibrated refined PSF values that supersede bead measurements for downstream analysis.

**Fig. S3:**
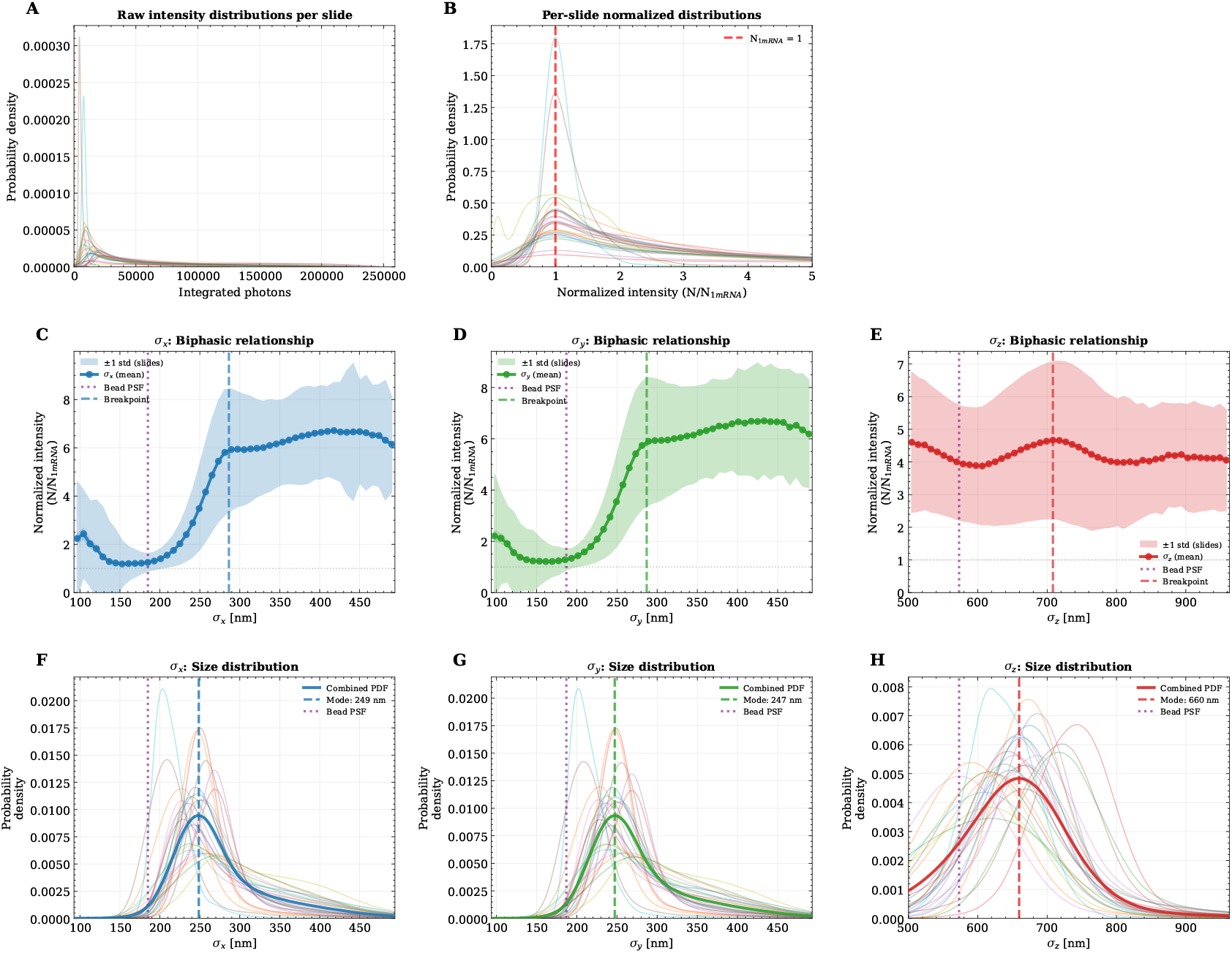
Breakpoint analysis for orange channel (fl-HTT). Analysis of 4,460,914 spots across 26 slides from Q111 mice (PFA < 1e-04). Median spots per slide: 171,573. **(A)** Raw intensity distributions per slide showing 2-3× slide-to-slide variation due to technical factors. **(B)** Normalized intensity distributions converging around *N*_1mRNA_ = 1 after per-slide modal normalization via KDE. **(C-E)** Size-intensity relationships. Breakpoints: *σ*_*x*_ = 286.1 nm (54.7% larger than bead PSF), *σ*_*y*_ = 286.8 nm (53.4% larger), *σ*_*z*_ = 708.4 nm (23.6% larger). Piecewise linear regression: *σ*_*x*_ slope1 = 0.060 (single-molecule regime), slope2 = 0.005 (cluster regime). **(F-H)** Spot size distributions. Mode values: *σ*_*x*_ = 248.6 nm (34.4% larger than bead PSF), *σ*_*y*_ = 247.2 nm (32.2%), *σ*_*z*_ = 659.6 nm (15.1%). Size statistics: *σ*_*x*_ mean = 281.2 nm, median = 262.1 nm, SD = 76.1 nm. Consistent breakpoint values between channels (∼285 nm lateral, ∼705 nm axial) validate the quantitative framework and reflect the physical extent of RNAscope probe clusters (∼20 probe pairs) bound to target mRNA.

**Fig. S4:**
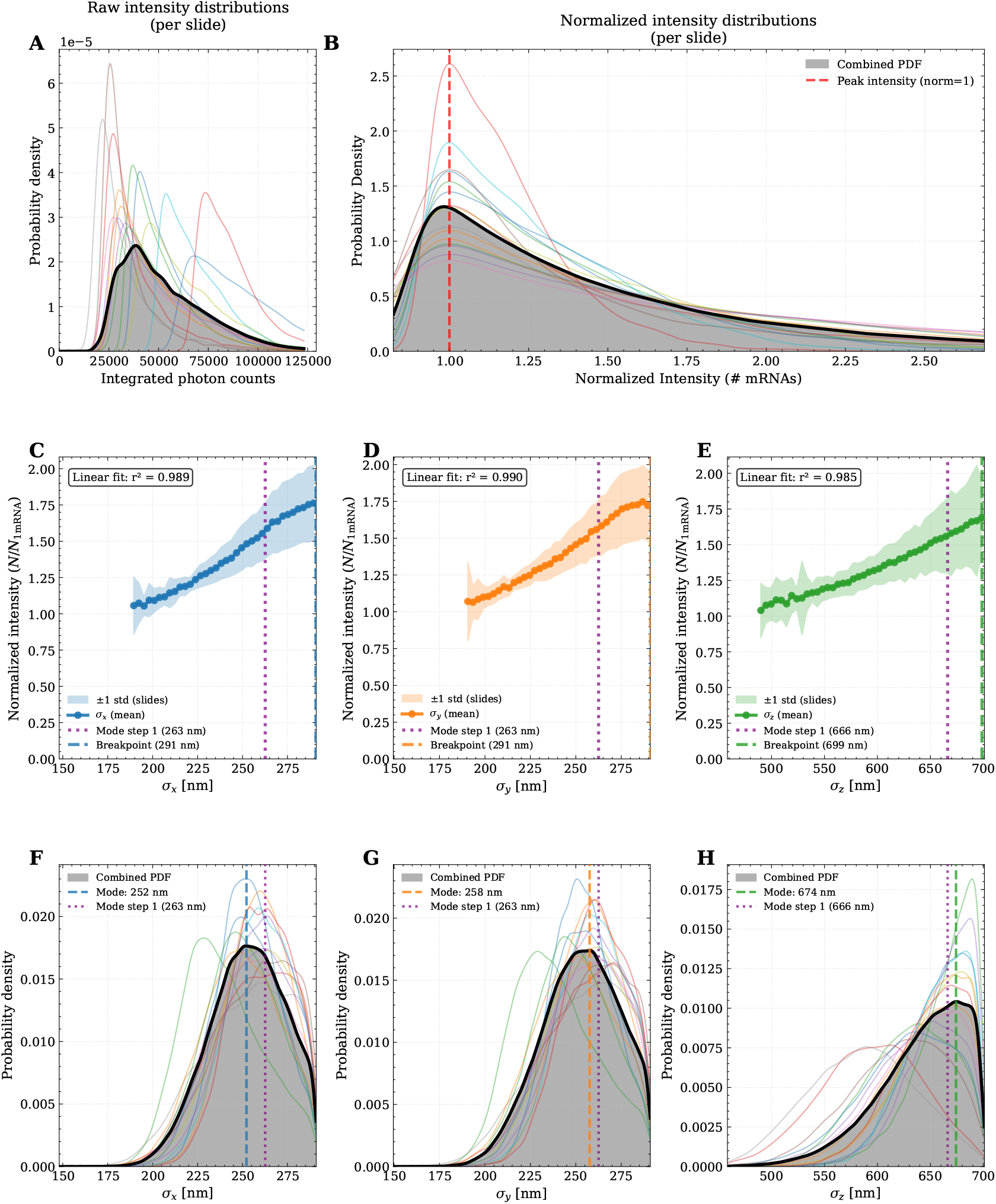
Validation of single-molecule filtering (green channel, HTT1a). All panels show data from the green channel (488 nm) corresponding to the HTT1a probe in Q111 transgenic mouse tissue. Analysis of orange channel (548 nm, fl-HTT) data yields similar results (fig. S5). **(A-B)** Per-slide intensity normalization demonstrates successful removal of technical variation. **(A)** Raw integrated photon counts for 15 slides (faint colored lines, 178,070 total spots) showing 2-3× variation; thick black line = combined PDF. **(B)** After per-slide normalization, all distributions converge around *N*_1mRNA_ = 1 (red dashed line) via KDE with Scott’s bandwidth [43, 44]. **(C-E)** Biphasic relationships between normalized intensity and spot size after filtering validate the single-molecule regime. For each dimension, the bold line shows mean normalized intensity across 15 slides binned into 50 size bins (only bins with ≥100 spots shown). Purple dotted lines: bead-derived PSF (*σ*_*x*_ = 263 nm, *σ*_*y*_ = 263 nm, *σ*_*z*_ = 666 nm). Colored dashed lines: empirically-determined breakpoints (*σ*_*x*_ = 290.8 nm, *σ*_*y*_ = 291.1 nm, *σ*_*z*_ = 698.6 nm; 11%, 11%, 5% larger than bead PSF). Strong linear relationships (Pearson *r*^2^ > 0.90) confirm single-molecule regime. **(F-H)** Size distributions after filtering show tight clustering around modal values with clear truncation at breakpoints. Colored dashed lines mark modes defining tissue-calibrated refined PSF. **Quality filters:** (i) PFA < 5e-02, (ii) *σ*_*i*_ < breakpoint for all dimensions, (iii) intensity > slide-specific negative-control threshold, (iv) cluster CV ≥ 0.5. **Slides used (n=15):** m1a4, m1a5, m1b1, m2a2, m2a3, m2a4, m2a8, m2b1, m2b7, m3a1, m3a2, m3a3, m3a5, m3b2, m3b3. **Slides excluded (n=14):** poor tissue integrity or low positive control expression.

**Fig. S5:**
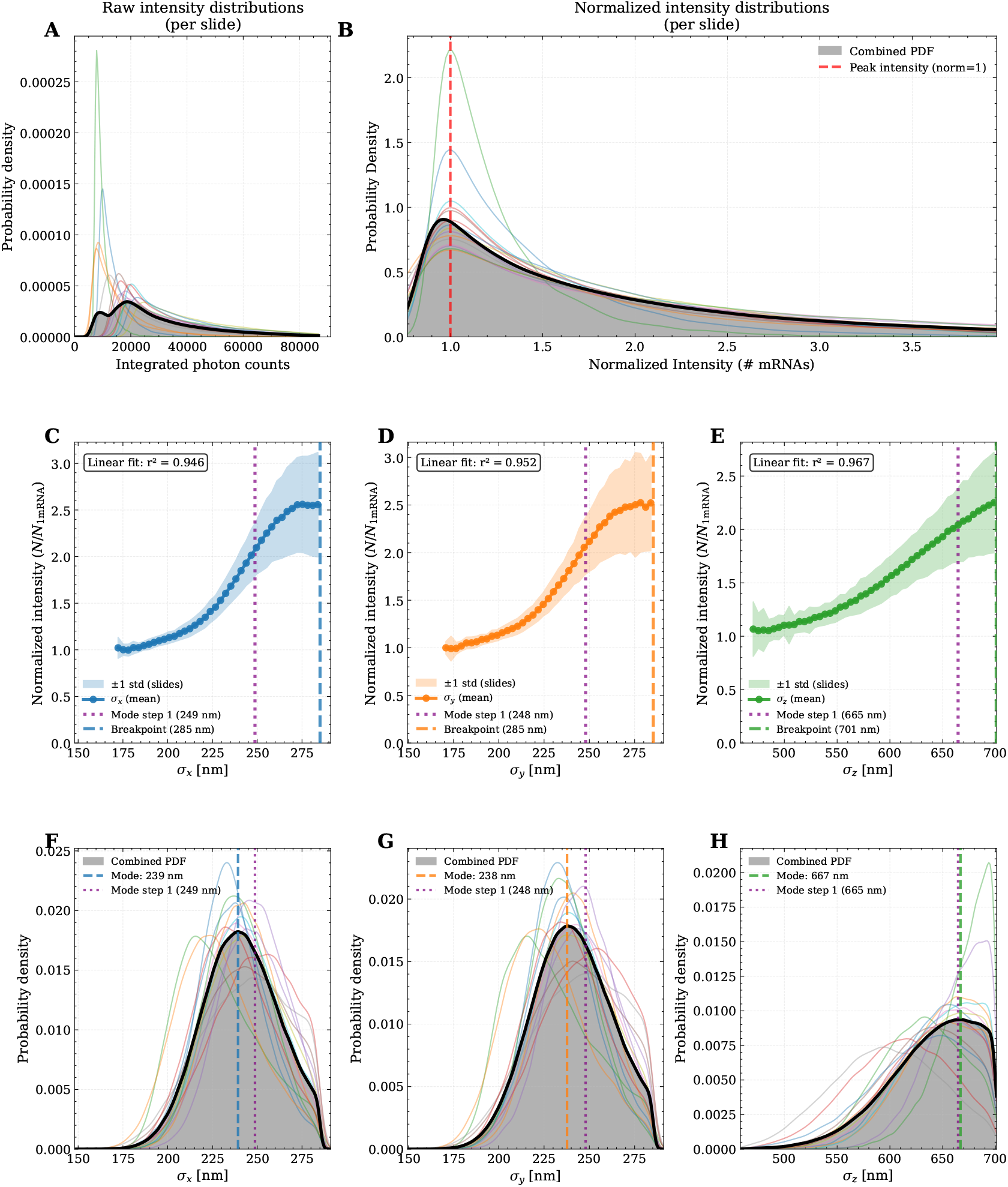
Validation of single-molecule filtering (orange channel, fl-HTT). All panels show data from the orange channel (548 nm) corresponding to the fl-HTT probe in Q111 transgenic mouse tissue. Analysis of green channel (488 nm, HTT1a) data yields similar results (fig. S4). **(A-B)** Per-slide intensity normalization demonstrates successful removal of technical variation. **(A)** Raw integrated photon counts for 15 slides (faint colored lines, 746,805 total spots) showing 2-3× variation; thick black line = combined PDF. **(B)** After per-slide normalization, all distributions converge around *N*_1mRNA_ = 1 (red dashed line) via KDE with Scott’s bandwidth [43, 44]. **(C-E)** Biphasic relationships between normalized intensity and spot size after filtering validate the single-molecule regime. For each dimension, the bold line shows mean normalized intensity across 15 slides binned into 50 size bins (only bins with ≥100 spots shown). Purple dotted lines: bead-derived PSF (*σ*_*x*_ = 249 nm, *σ*_*y*_ = 248 nm, *σ*_*z*_ = 665 nm). Colored dashed lines: empirically-determined breakpoints (*σ*_*x*_ = 285.1 nm, *σ*_*y*_ = 285.3 nm, *σ*_*z*_ = 700.7 nm; 15%, 15%, 5% larger than bead PSF). Strong linear relationships (Pearson *r*^2^ > 0.90) confirm single-molecule regime. **(F-H)** Size distributions after filtering show tight clustering around modal values with clear truncation at breakpoints. Colored dashed lines mark modes defining tissue-calibrated refined PSF. **Quality filters:** (i) PFA < 5e-02, (ii) *σ*_*i*_ < breakpoint for all dimensions, (iii) intensity > slide-specific negative-control threshold, (iv) cluster CV ≥ 0.5. **Slides used (n=15):** m1a4, m1a5, m1b1, m2a2, m2a3, m2a4, m2a8, m2b1, m2b7, m3a1, m3a2, m3a3, m3a5, m3b2, m3b3. **Slides excluded (n=14):** poor tissue integrity or low positive control expression.

**Fig. S6:**
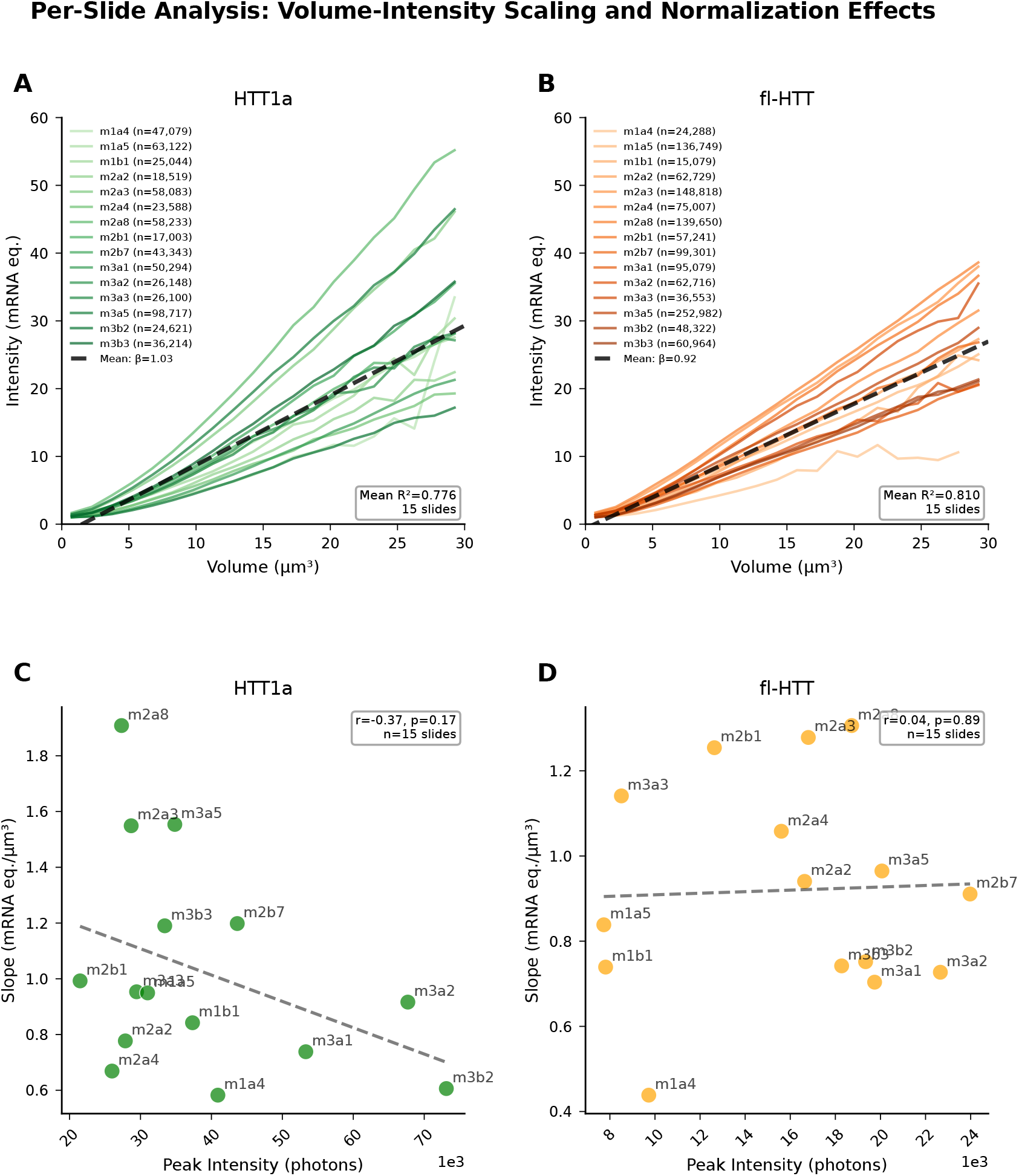
Per-slide analysis of volume-intensity scaling validates normalization and reveals biological variability in mRNA packing density. This figure examines consistency of the volume-intensity relationship across individual slides and evaluates whether per-slide normalization introduces systematic biases. *Key question:* Is inter-slide variability in slopes driven by technical factors (normalization errors) or biological differences (mRNA packing density)? **(A)** Green channel (488 nm, mHTT1a): Per-slide binned mean intensity vs. cluster volume. Each colored line = one slide (n = 15 slides, 616,100 clusters within 0 to 30 *µ*m^3^). Dashed black line: mean slope *β* = 1.03 mRNA eq./*µ*m^3^. Bins shown have ≥5 clusters. **(B)** Orange channel (548 nm, fl-mHTT): Same analysis. n = 15 slides, 1,314,992 clusters. Mean slope *β* = 0.92 mRNA eq./*µ*m^3^. **(C)** Green channel: Peak intensity (photons) vs. slope correlation. Each point = one slide. Pearson *r* = −0.37, *p* = 0.17 (not significant). Peak intensity = modal intensity of single-molecule spots (*N*_1mRNA_) used for normalization. **(D)** Orange channel: Peak intensity vs. slope correlation. Pearson *r* = 0.04, *p* = 0.89 (not significant). **Slope statistics:** Green: mean = 1.03, SD = 0.37 mRNA eq./*µ*m^3^, CV = 36%, range = 0.58-1.91. Orange: mean = 0.92, SD = 0.24 mRNA eq./*µ*m^3^, CV = 26%, range = 0.44-1.31. **Per-slide** *R*^2^: Green: mean = 0.78, range = 0.37-0.87. Orange: mean = 0.81, range = 0.43-0.89. **Key findings:** (1) All slides show positive linear relationships, confirming the volume-intensity scaling is robust. (2) *No significant correlation* between peak intensity and slope for either channel validates that normalization does not introduce systematic bias: if errors in *N*_1mRNA_ estimation were driving slope variability, we would observe strong correlations. (3) The observed inter-slide variability (CV = 26-36%) reflects genuine biological differences: variation in mRNA packing density within clusters across different animals and tissue sections, regional heterogeneity, and disease progression state. (4) High within-slide *R*^2^ values (>0.78 mean) confirm the linear model is appropriate. **Interpretation:** The per-slide normalization successfully removes technical variation (2-3× raw photon count differences between slides) while preserving biologically meaningful differences in mRNA density.

**Fig. S7:**
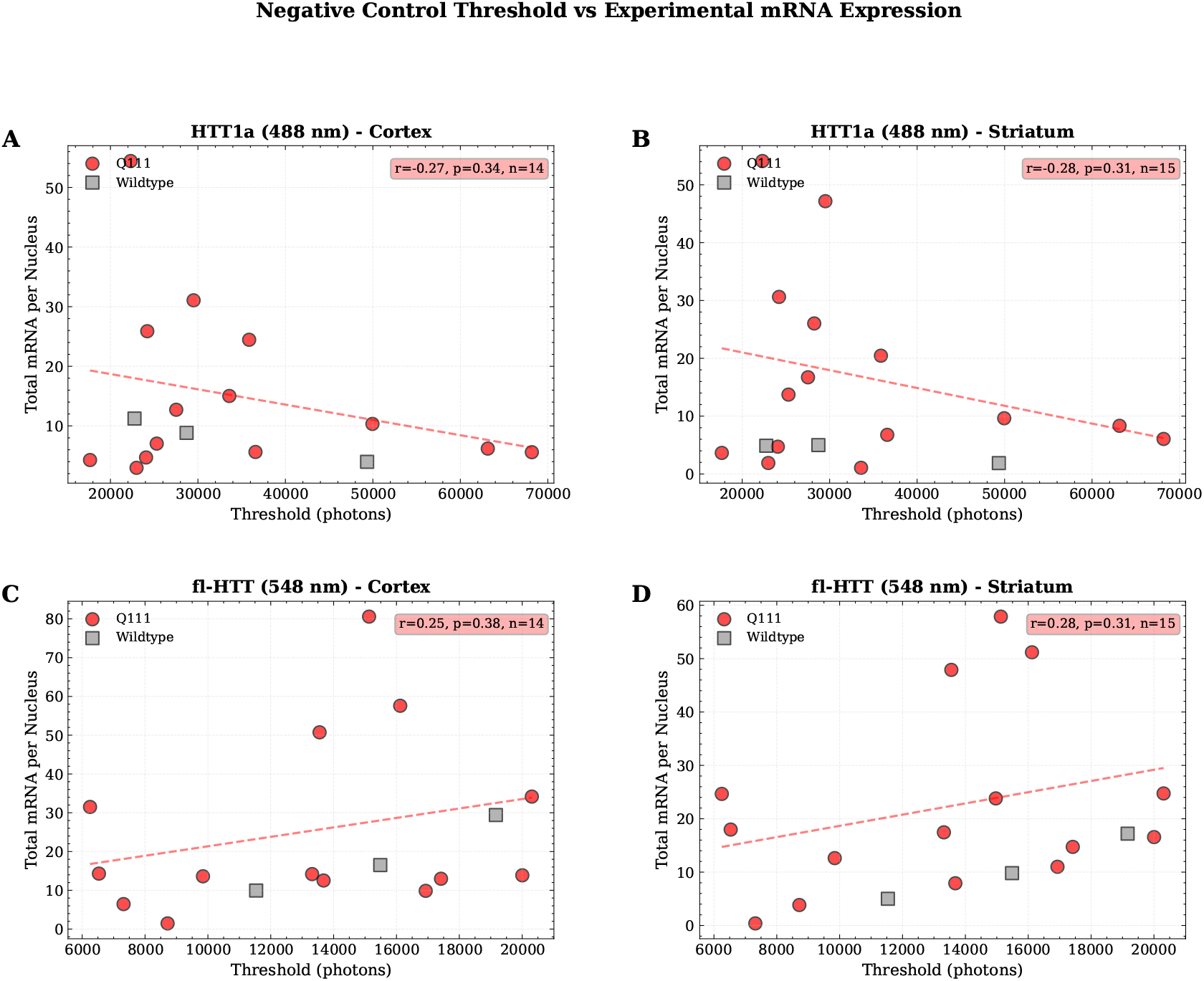
Negative control thresholds show no correlation with experimental mRNA expression. Validation that negative control background thresholds do not correlate with experimental mRNA expression levels. A *lack* of correlation validates that experimental signal represents true biological mRNA, not technical artifacts influenced by background. **Dataset:** 18 unique slides (Q111 transgenic: 15, Wildtype: 3). Channels: HTT1a (488 nm, exon 1) and fl-HTT (548 nm). Regions: Cortex and Striatum. Merged threshold = average of Cortex and Striatum thresholds per slide (based on 95th percentile of DapB negative control intensities). **(A) HTT1a - Cortex:** Q111 n=14, Wildtype n=3. Q111 mRNA: 15 ± 15 mRNA/nucleus; Wildtype: 8 ± 4 mRNA/nucleus. Q111 correlation: *r* = −0.273, *p* = 0.345 (weak/no correlation). **(B) HTT1a - Striatum:** Q111 n=15, Wildtype n=3. Q111 mRNA: 17 ± 17 mRNA/nucleus; Wildtype: 4 ± 2 mRNA/nucleus. Q111 correlation: *r* = −0.283, *p* = 0.307. **(C) fl-HTT - Cortex:** Q111 n=14, Wildtype n=3. Q111 mRNA: 25 ± 23 mRNA/nucleus; Wildtype: 18 ± 10 mRNA/nucleus. Q111 correlation: *r* = +0.255, *p* = 0.379. **(D) fl-HTT - Striatum:** Q111 n=15, Wildtype n=3. Q111 mRNA: 22 ± 18 mRNA/nucleus; Wildtype: 10 ± 6 mRNA/nucleus. Q111 correlation: *r* = +0.284, *p* = 0.305. **Nuclear volume normalization:** Mean nuclear volume = 716 *µ*m^3^; mRNA counts normalized per nucleus to account for 3D sectioning of nuclei in tissue sections. **Key findings:** All 4 channel-region combinations show weak correlation (|*r*| < 0.3, mean |*r*| = 0.274, all *p* > 0.3). Wildtype n=3 insufficient for reliable correlation analysis. **Interpretation:** Weak/no correlation validates that mRNA quantification is independent of background threshold: observed mRNA levels reflect true biological expression, not threshold-dependent bias.

**Fig. S8:**
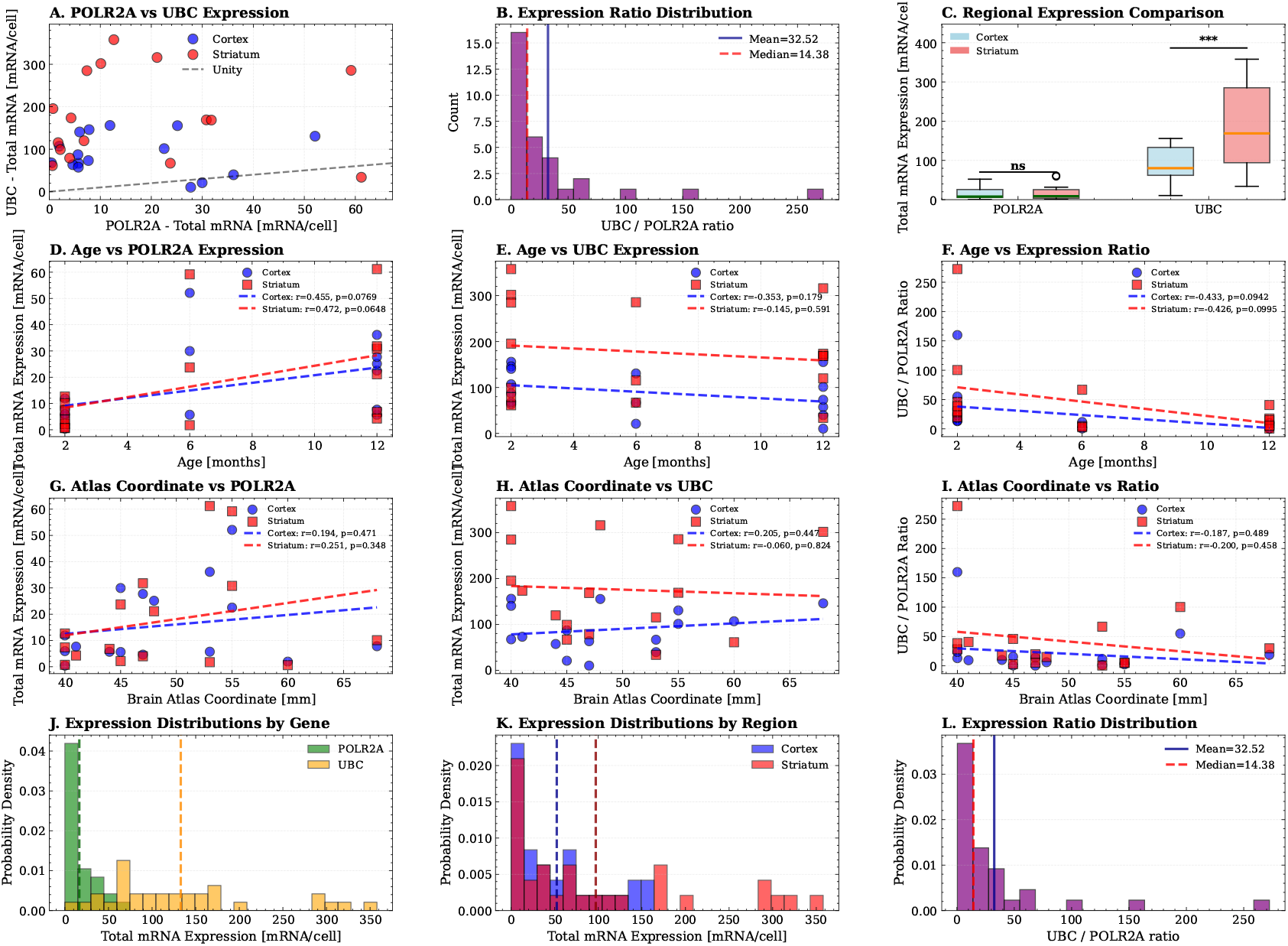
Positive control analysis validates assay dynamic range. Analysis of POLR2A (low-expression housekeeping gene, green channel) and UBC (high-expression housekeeping gene, orange channel) in positive control samples from Q111 and Wildtype mice. **Slides analyzed (n=16):** m1a4, m1b1, m2a2, m2a3, m2a4, m2a6, m2a8, m2b1, m2b6, m2b7, m3a1, m3a3, m3a4, m3a5, m3b2, m3b3. **Slides excluded (n=13):** poor tissue integrity. **Total FOVs:** 764; **Paired samples (slides with both Cortex and Striatum):** 16; **Age range:** 2-12 months; **Atlas coordinate range:** 40-68 (sections spaced at 100 *µ*m intervals). **Row 1 (A-C):** Gene expression comparisons. (A) POLR2A vs UBC scatter plot colored by region. (B) UBC/POLR2A ratio histogram. (C) Regional comparison box plots. POLR2A: Cortex 15.78 ± 14.76 mRNA/cell (n=16 slides), Striatum 17.47 ± 19.49 mRNA/cell; paired t-test *t* = −0.909, *p* = 0.38. UBC: Cortex 89.34 ± 47.37 mRNA/− cell, Striatum 177.28 ± 104.08 mRNA/cell; *t* = −4.809, *p* = 0.00023. UBC/POLR2A ratio across both regions combined: mean 32.89 ± 57.05, median 14.10 (Cortex alone: 21.93; Striatum alone: 43.84). Note: the striatum-only ratio (43.84) is used in fig. S11 for comparison with striatum-derived RNA-seq data. **Row 2 (D-F):** Age dependencies for POLR2A, UBC, and UBC/POLR2A ratio with separate Cortex/Striatum regression lines. Non-significant correlations suggest robust normalization. **Row 3 (G-I):** Brain atlas coordinate dependencies with regional regressions. Tests for anterior-posterior expression gradients. **Row 4 (J-L):** Expression distributions by gene and region. UBC shows higher expression than POLR2A, as expected for a high-abundance housekeeping gene.

**Fig. S9:**
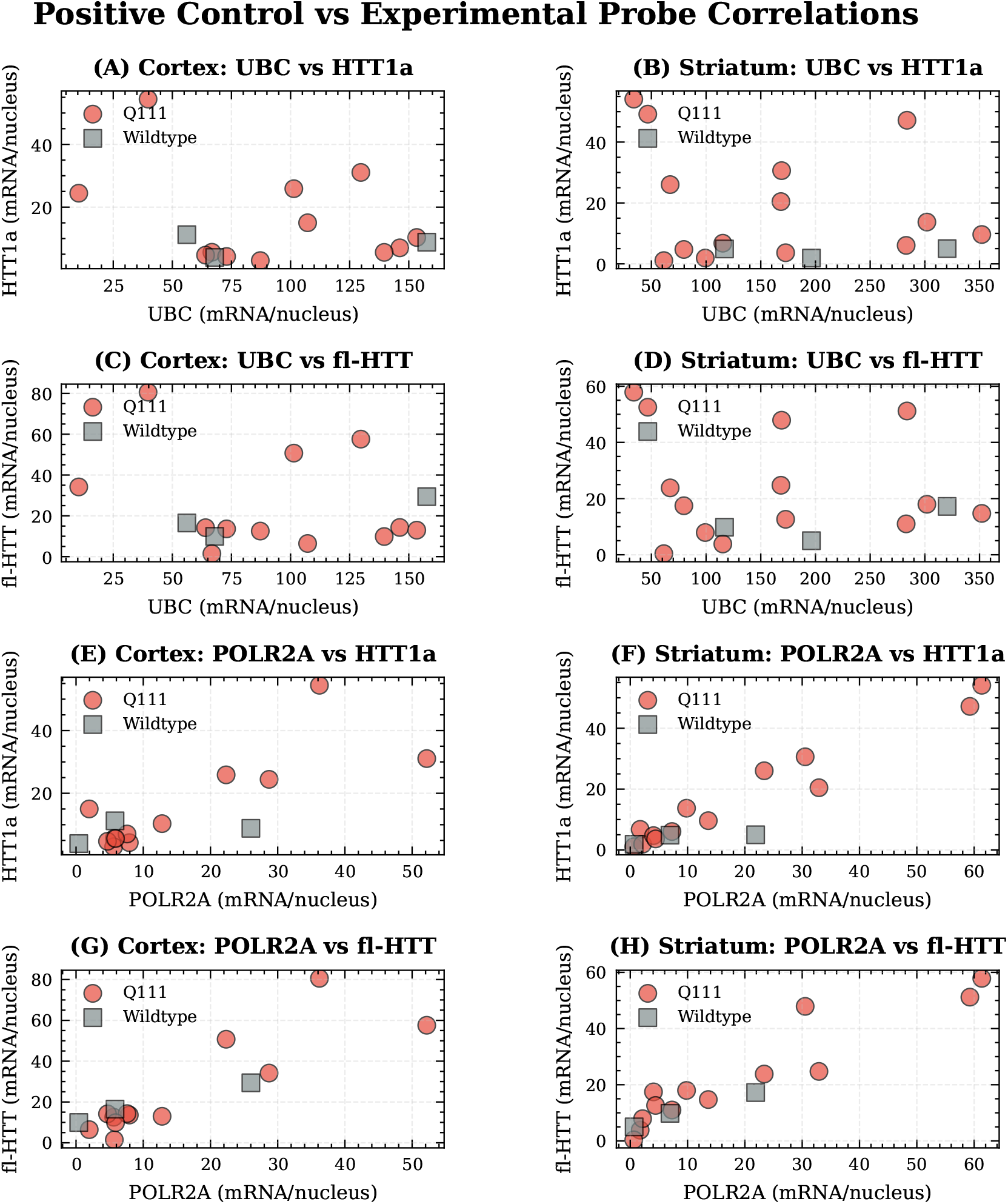
Positive control correlations with experimental probes. Tests whether variability in mHTT probes is driven by technical factors (tissue quality, RNA preservation) or true biological variation. Each point = one slide (mean expression across all FOVs). Red circles: Q111; gray squares: wildtype. Sample sizes: Cortex n=12, Striatum n=13. **(A-B)** UBC vs mHTT1a. Cortex Q111: *r* = −0.34, *p* = 0.29; Striatum Q111: *r* = −0.05, *p* = 0.86. Weak correlations. **(C-D)** UBC vs fl-mHTT. Cortex Q111: *r* = −0.28, *p* = 0.37; Striatum Q111: *r* = −0.00, *p* = 0.99. Weak correlations. **(E-F)** POLR2A vs mHTT1a. Cortex Q111: *r* = 0.81, *p* = 0.001**; Striatum Q111: *r* = 0.98, *p* < 0.001***. Strong correlations. **(G-H)** POLR2A vs fl-mHTT. Cortex Q111: *r* = 0.87, *p* < 0.001***; Striatum Q111: *r* = 0.94, *p* < 0.001***. Strong correlations. **Interpretation:** UBC shows weak correlations with mHTT probes, indicating experimental variability is not driven by UBC-related tissue quality. POLR2A shows strong correlations; whether this reflects RNA quality confounding or true biological coupling is addressed in fig. S10.

**Fig. S10:**
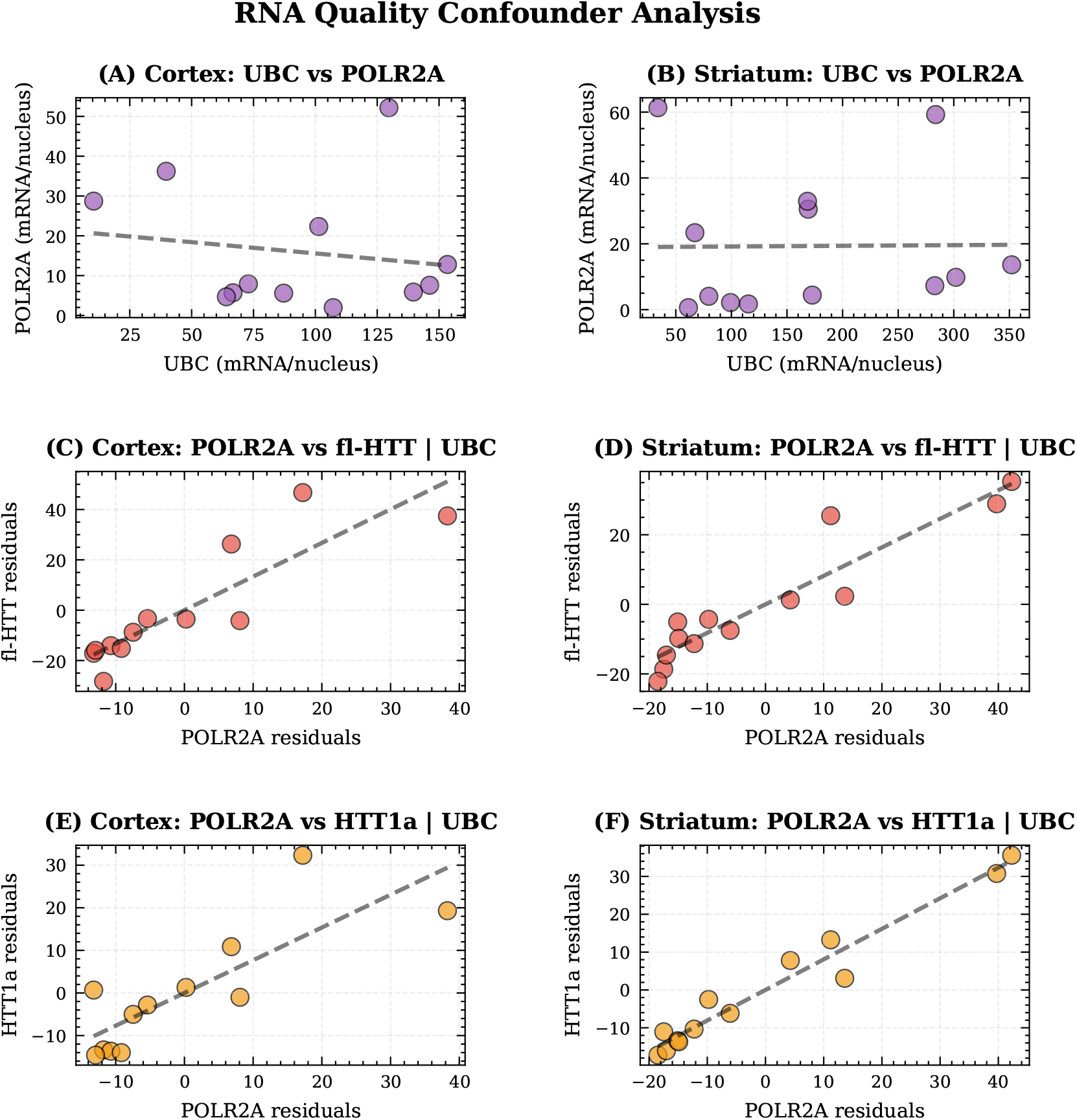
Partial correlation analysis rules out RNA quality as major confounder. Addresses whether strong POLR2A-mHTT correlations are spurious (driven by RNA quality) or reflect true biological coupling. Uses partial correlation controlling for UBC as a tissue quality proxy. Sample sizes: Cortex n=12, Striatum n=13. **(A-B)** UBC vs POLR2A correlation quantifies shared technical variance. Cortex: *r* = −0.161, *p* = 0.62; Striatum: *r* = 0.010, *p* = 0.97. Low correlations indicate UBC and POLR2A respond differently to tissue quality. **(C-D)** Partial correlation: POLR2A vs fl-mHTT | UBC. Cortex: original *r* = 0.872, partial *r* = 0.873, *p* < 0.001*** (+0.1%); Striatum: original *r* = 0.939, partial *r* = 0.939, *p* < 0.001*** (0.0%). **(E-F)** Partial correlation: POLR2A vs mHTT1a | UBC. Cortex: original *r* = 0.809, partial *r* = 0.812, *p* = 0.001** (+0.4%); Striatum: original *r* = 0.975, partial *r* = 0.977, *p* < 0.001*** (+0.2%). **Conclusions:** (1) RNA quality is not a major confounder: partial correlations remain nearly identical after controlling for UBC. (2) Biological coupling is supported: correlations persist after removing tissue quality effects, suggesting co-transcriptional regulation or shared transcriptional machinery. (3) If correlations were spurious, we would expect high UBC-POLR2A correlation and large reductions in partial correlations; neither is observed.

**Fig. S11:**
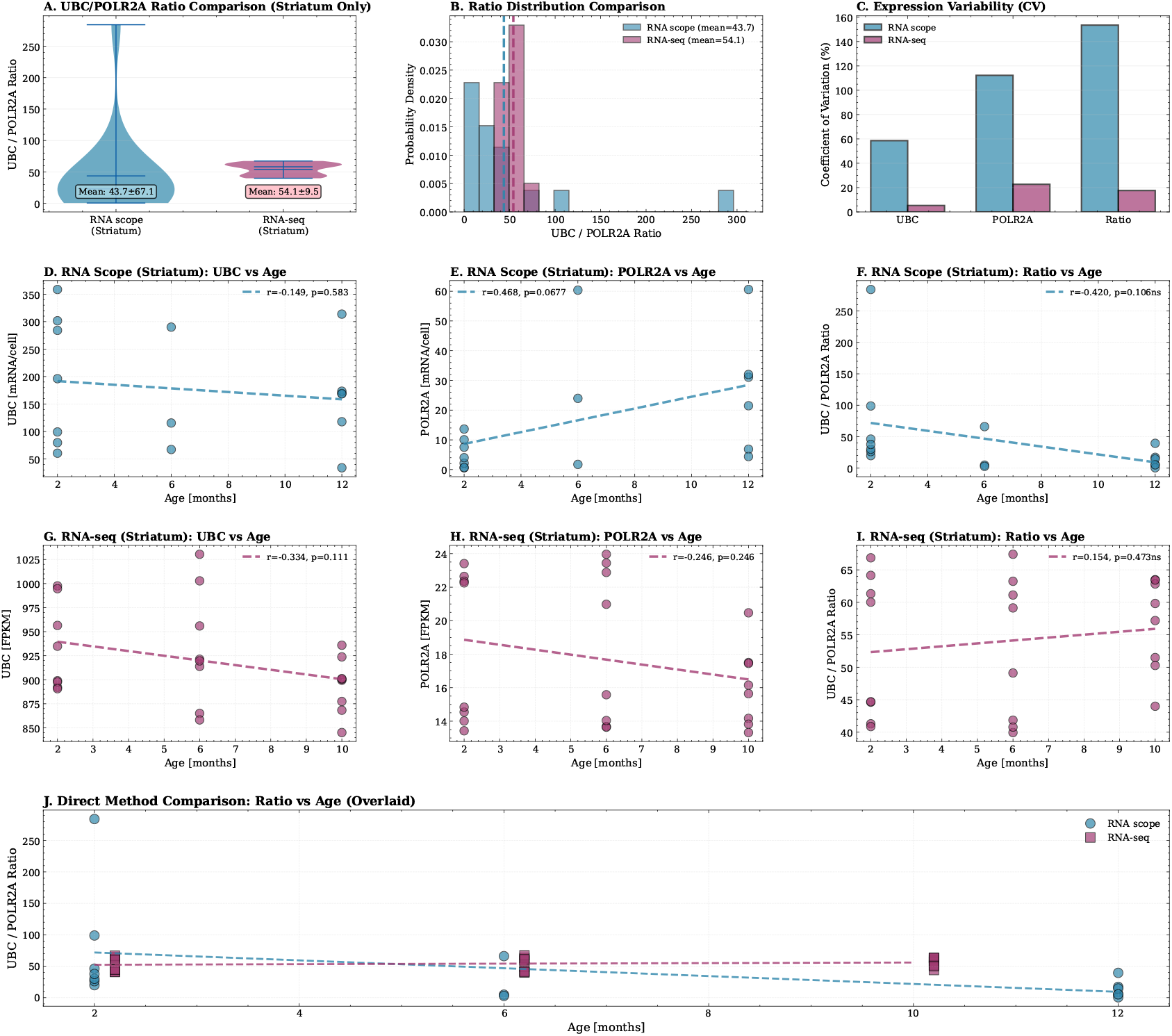
Comparison of RNAscope and bulk RNA-seq validates housekeeping gene quantification. Comparison between single-molecule RNAscope (this study; *n* = 16 slides from Q111 mice, ages 2-12 months, striatum only) and bulk RNA-seq (GSE65774 [10]; *n* = 24 Q111 striatum samples). **(A)** UBC/POLR2A ratio comparison (violin plots). This unit-independent ratio enables direct comparison despite different quantification approaches (mRNA/nucleus vs FPKM). RNAscope (striatum only): 44 ± 68; RNA-seq: 54 ± 10; fold difference 0.81× (*t* = −0.72, *p* = 0.48, not significant). Note: The RNAscope ratio here (43.8) is higher than the combined Cortex+Striatum ratio in fig. S8 (32.89) because this comparison uses striatum data only to match the RNA-seq dataset. **(B)** Overlaid histograms showing full distribution of UBC/POLR2A ratios; dashed lines indicate means. **(C)** Coefficient of variation comparison: UBC CV (RNAscope 58%, RNA-seq 5%), POLR2A CV (112% vs 23%), Ratio CV (154% vs 18%). Higher CV in RNAscope reflects biological heterogeneity at single-cell level that bulk RNA-seq averages out [9]. **(D-F)** RNAscope age correlations (striatum): UBC vs age (*r* = −0.14, *p* = 0.60), POLR2A vs age (*r* = 0.48, *p* = 0.06), ratio vs age (*r* = −0.42, *p* = 0.11). **(G-I)** RNA-seq age correlations: UBC vs age (*r* = −0.33, *p* = 0.11), POLR2A vs age (*r* = −0.25, *p* = 0.25), ratio vs age (*r* = 0.15, *p* = 0.47). **(J)** Direct method overlay showing both datasets on same axes. **Key findings:** (1) UBC/POLR2A ratios are comparable between methods (*p* = 0.49), validating RNAscope quantification. (2) Higher variance in RNAscope reflects preserved cell-to-cell heterogeneity. (3) Age correlations are qualitatively similar, supporting biological validity. Sampling note: RNAscope examines thin tissue sections (40 *µ*m) from specific anatomical locations, while RNA-seq uses homogenized bulk striatum; exact dissection boundaries for the RNA-seq dataset are not specified but likely represent larger tissue volumes.

**Fig. S12:**
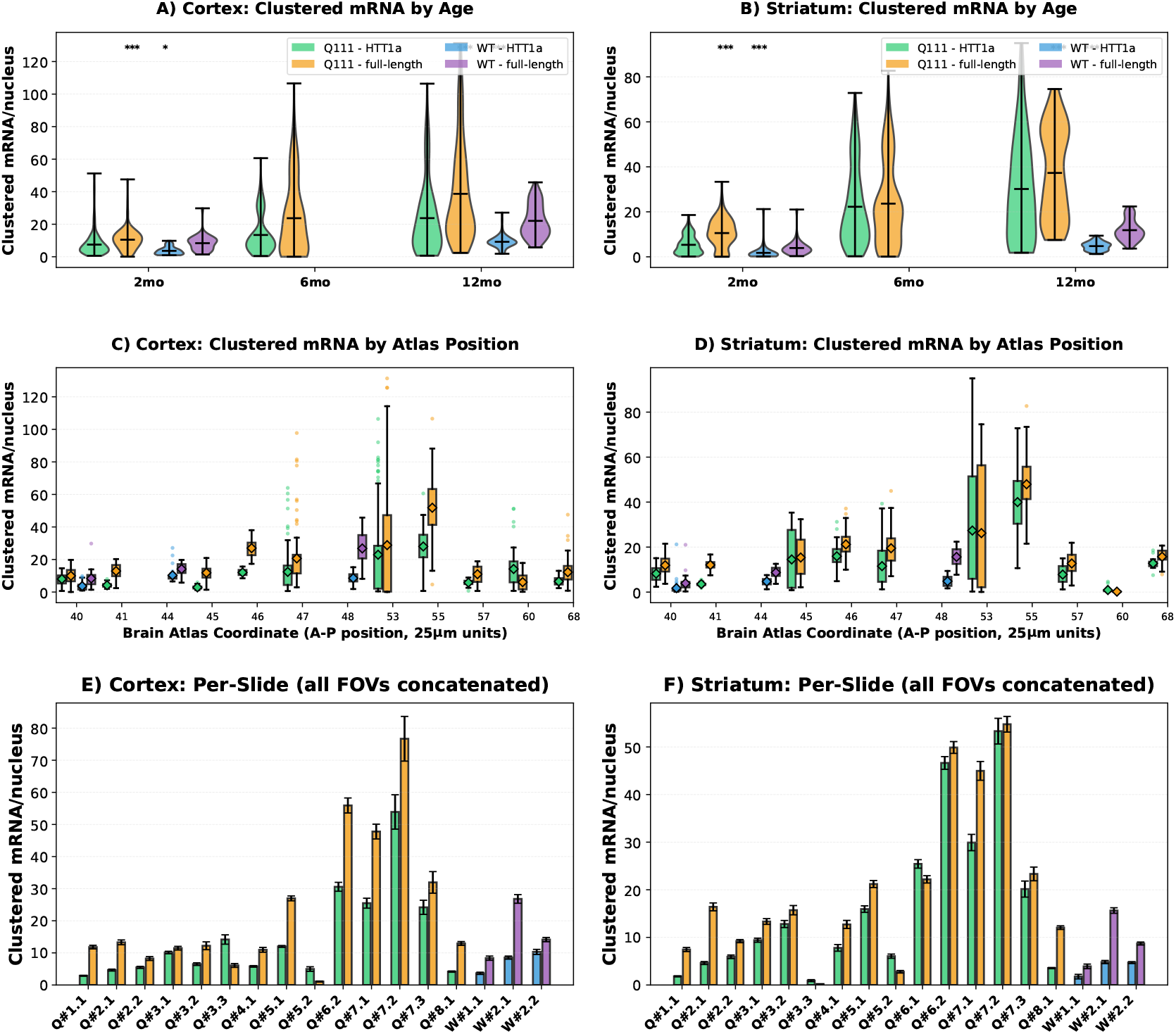
Clustered mRNA expression dominates total signal. Clustered mRNA represents clustered transcripts, quantified as (*I*_cluster_*/I*_peak_)*/N*_nuclei_. Dataset: 3,234 FOVs (Q111: 2,668; WT: 566). Cortex: 1,662 FOVs; Striatum: 1,572 FOVs. **(A-B) Age trends**. Violin plots showing clustered mRNA distribution across ages. Cortex: 2mo Q111 7.50 ± 6.79 mRNA/cell (n=335), 6mo 13.18 ± 11.28 (n=193), 12mo Q111 23.86 ± 22.17 (n=169); 12mo WT 9.19 ± 3.89 (n=83). Striatum: 2mo Q111 5.33 ± 4.19 (n=180), 6mo 21.85 ± 16.82 (n=284), 12mo Q111 30.24 ± 22.76 (n=173); 12mo WT 4.75 ± 1.88 (n=104). **Statistical tests (Q111 vs WT):** Striatum 2mo HTT1a: *t* = 5.27, *p* < 0.0001; Striatum 2mo fl-HTT: *t* = 6.81, *p* < 0.0001; Striatum 12mo HTT1a: *t* = 11.39, *p* < 0.0001; Striatum 12mo fl-HTT: *t* = 13.04, *p* < 0.0001. **(C-D)** Box plots showing expression across atlas coordinates (40-68, sections spaced at 100 *µ*m intervals). Q111 HTT1a Striatum: mean 19.46 ± 19.02 mRNA/cell (n=637 FOVs); WT: 3.86 ± 2.74 (n=149 FOVs). **(E-F)** Per-slide bar plots showing mean±SEM expression, revealing inter-individual variability.

**Fig. S13:**
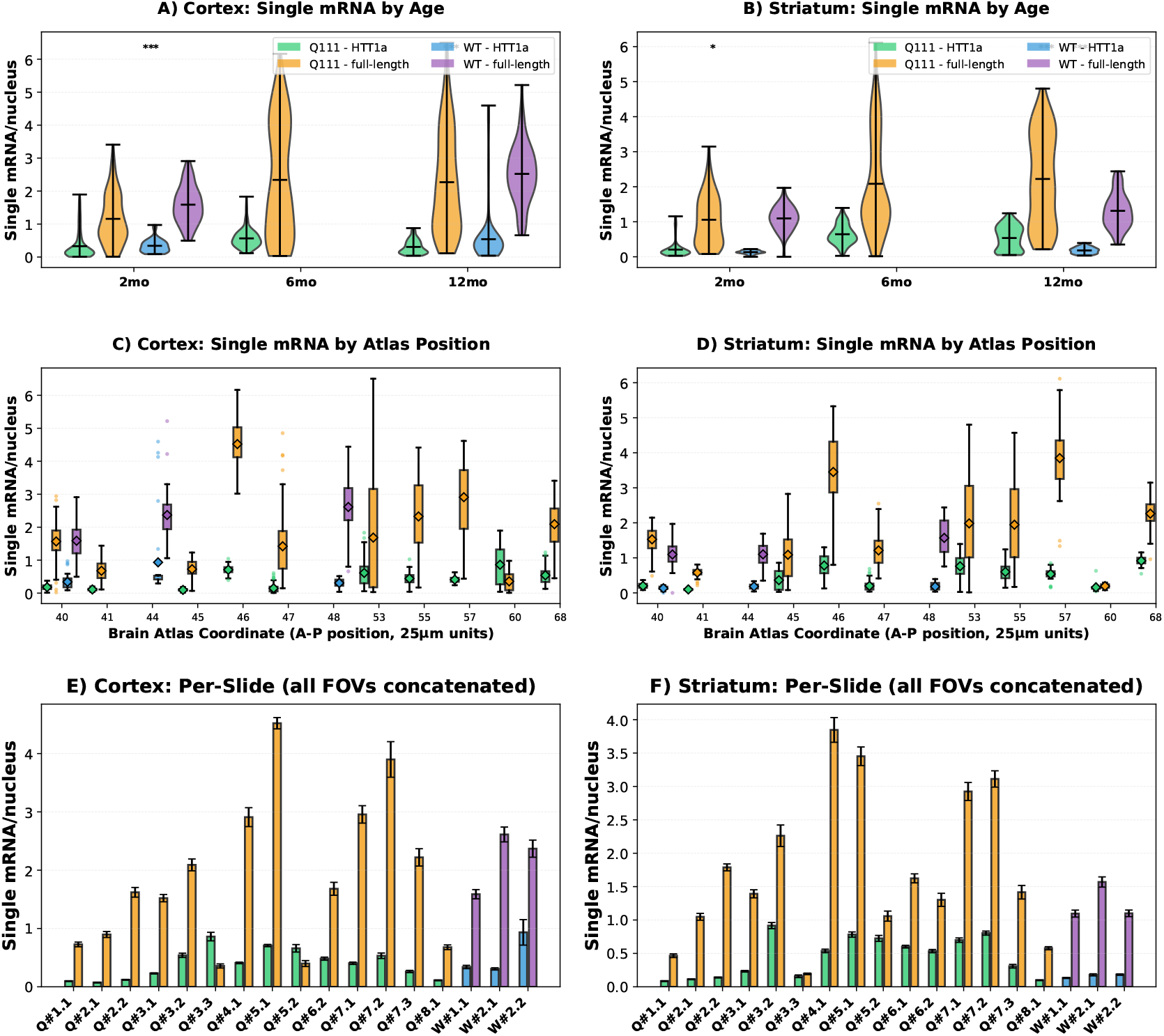
Single mRNA expression analysis. Single mRNA represents individual diffraction-limited spots, quantified as *N*_spots_*/N*_nuclei_ per nucleus. Dataset: 3,234 FOVs (Q111: 2,668; WT: 566). **(A-B) Age trends**. Violin plots showing distribution of single mRNA expression across ages. Cortex: 2mo Q111 HTT1a 0.33 ± 0.42 (n=335), WT 0.36 ± 0.21 (n=51); 6mo Q111 0.57 ± 0.29 (n=193); 12mo Q111 0.31 ± 0.21 (n=169), WT 0.54 ± 0.80 (n=83). Striatum: 2mo Q111 0.21 ± 0.23 (n=180), WT 0.14 ± 0.05 (n=45); 6mo Q111 0.65 ± 0.29 (n=284); 12mo Q111 0.54 ± 0.34 (n=173), WT 0.18 ± 0.09 (n=104). **Statistical tests (Q111 vs WT):** Striatum 2mo HTT1a: *t* = 1.92, *p* = 0.056 (ns); Striatum 2mo fl-HTT: *t* = −0.05, *p* = 0.96 (ns); Striatum 12mo HTT1a: *t* = 10.49, *p* < 0.0001; Striatum 12mo fl-HTT: *t* = 7.34, *p* < 0.0001. **(C-D)** Atlas coordinate trends. Box plots showing expression across approximate anterior-posterior coordinates (atlas sections spaced at 100 *µ*m intervals). Q111 HTT1a: cortex 0.39 ± 0.36 (n=697), striatum 0.49 ± 0.34 (n=637). WT HTT1a: cortex 0.47 ± 0.65 (n=134), striatum 0.17 ± 0.08 (n=149). **(E-F)** Per-slide breakdown showing mean ± SEM for each slide (all FOVs concatenated per slide).

**Fig. S14:**
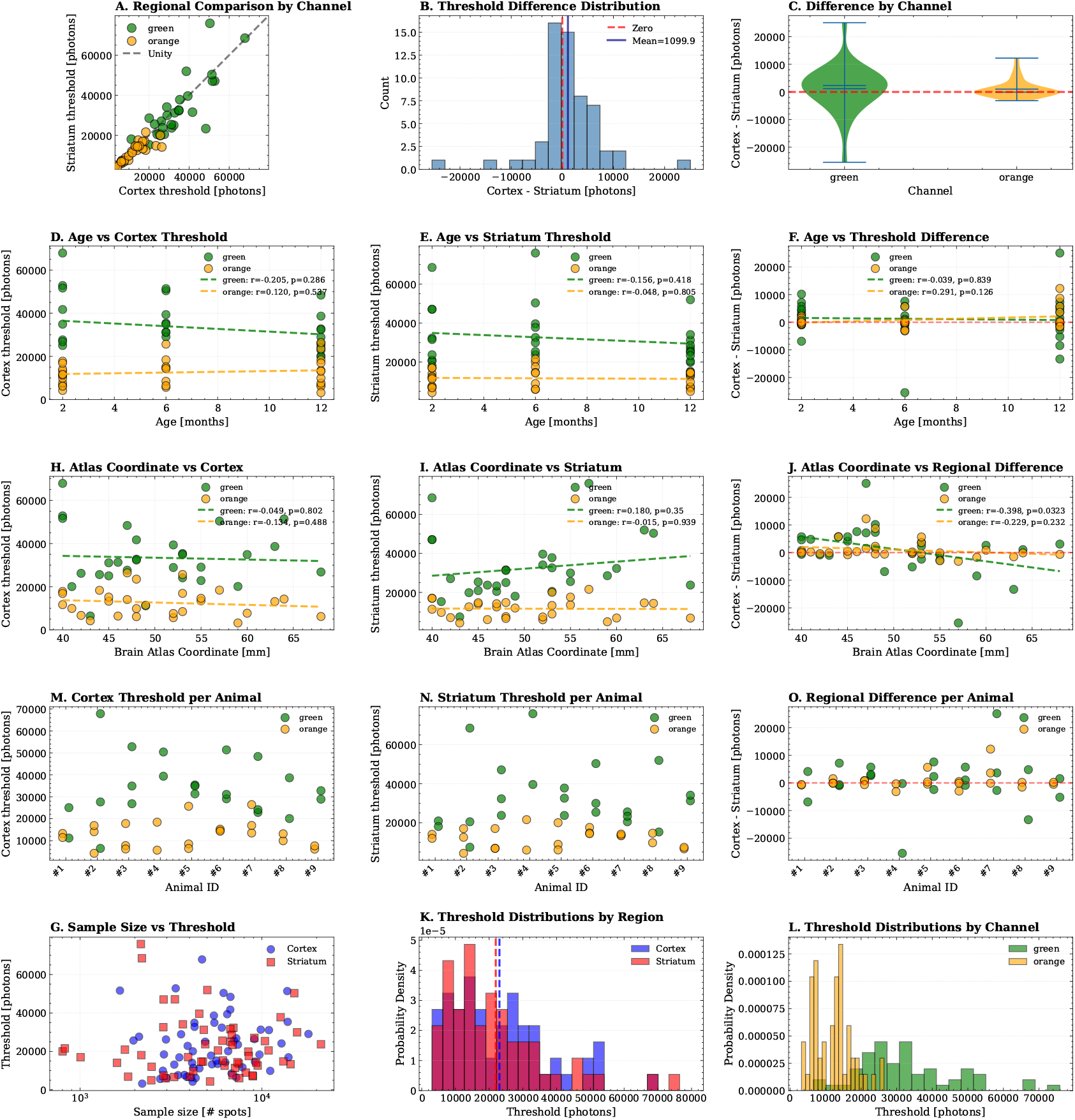
Comprehensive negative control background threshold analysis across brain regions. Analysis of regional differences in negative control thresholds (Cortex vs Striatum) and their relationships with age, sample size, and brain atlas coordinates. Negative control thresholds define the intensity cutoff above which spots are considered true mRNA. **Dataset:** 58 paired samples (slide-channel combinations), 29 unique slides. Channels: green (488 nm, HTT1a) and orange (548 nm, fl-HTT). Age range: 2-12 months. Brain atlas coordinates: 40-68 (sections spaced at 100 *µ*m intervals). **Overall regional comparison:** Cortex threshold: 23, 037 ± 14, 661 photons; Striatum: 21, 938 ± 15, 334 photons. Mean difference (Cortex − Striatum): +1, 099 ± 6, 469 photons. Paired *t*-test: *t* = 1.294, *p* = 0.201 (*no significant regional difference*). **Per-channel statistics:** Green (n=29): Cortex 33, 404 ± 13, 292, Striatum 32, 262 ± 15, 369 photons (*p* = 0.481). Orange (n=29): Cortex 12, 670 ± 6, 183, Striatum 11, 614 ± 4, 655 photons (*p* = 0.100). **Age correlations:** Age vs Cortex threshold: *r* = −0.067, *p* = 0.617; Age vs Striatum: *r* = −0.085, *p* = 0.528; Age vs regional difference: *r* = +0.048, *p* = 0.719 (*no significant age effects*). **Rostrocaudal position:** Atlas coordinate vs Cortex threshold: *r* = −0.052, *p* = 0.70; Atlas coordinate vs Striatum threshold: *r* = +0.087, *p* = 0.52 (both negligible). The regional *difference* shows a modest correlation with atlas position (*r* = −0.323, *p* = 0.013), but this reflects small opposing trends that largely cancel when regions are merged. **Row 1 (A-C):** Regional comparison scatter, difference histogram, and violin plots by channel. **Row 2 (D-F):** Age dependencies for Cortex, Striatum, and their difference. **Row 3 (H-J):** Brain atlas coordinate dependencies. **Row 4 (M-O):** Per-animal analysis. Box plots showing threshold distributions for each animal (n=9), enabling assessment of inter-animal vs intra-animal variability. Panel M: Cortex threshold per animal. Panel N: Striatum threshold per animal. Panel O: Regional difference (Cortex − Striatum) per animal. **Row 5 (G, K-L):** Sample size effects and threshold distributions. **Conclusion:** No significant regional difference (*p* = 0.201), no age effect, and negligible rostrocaudal correlations for individual regions. The precise anatomical placement of the negative control imaging region within a slide does not meaningfully affect threshold determination. Per-slide calibration accounts for slide-to-slide technical variation; green channel requires higher thresholds due to tissue autofluorescence at 488 nm.

**Fig. S15:**
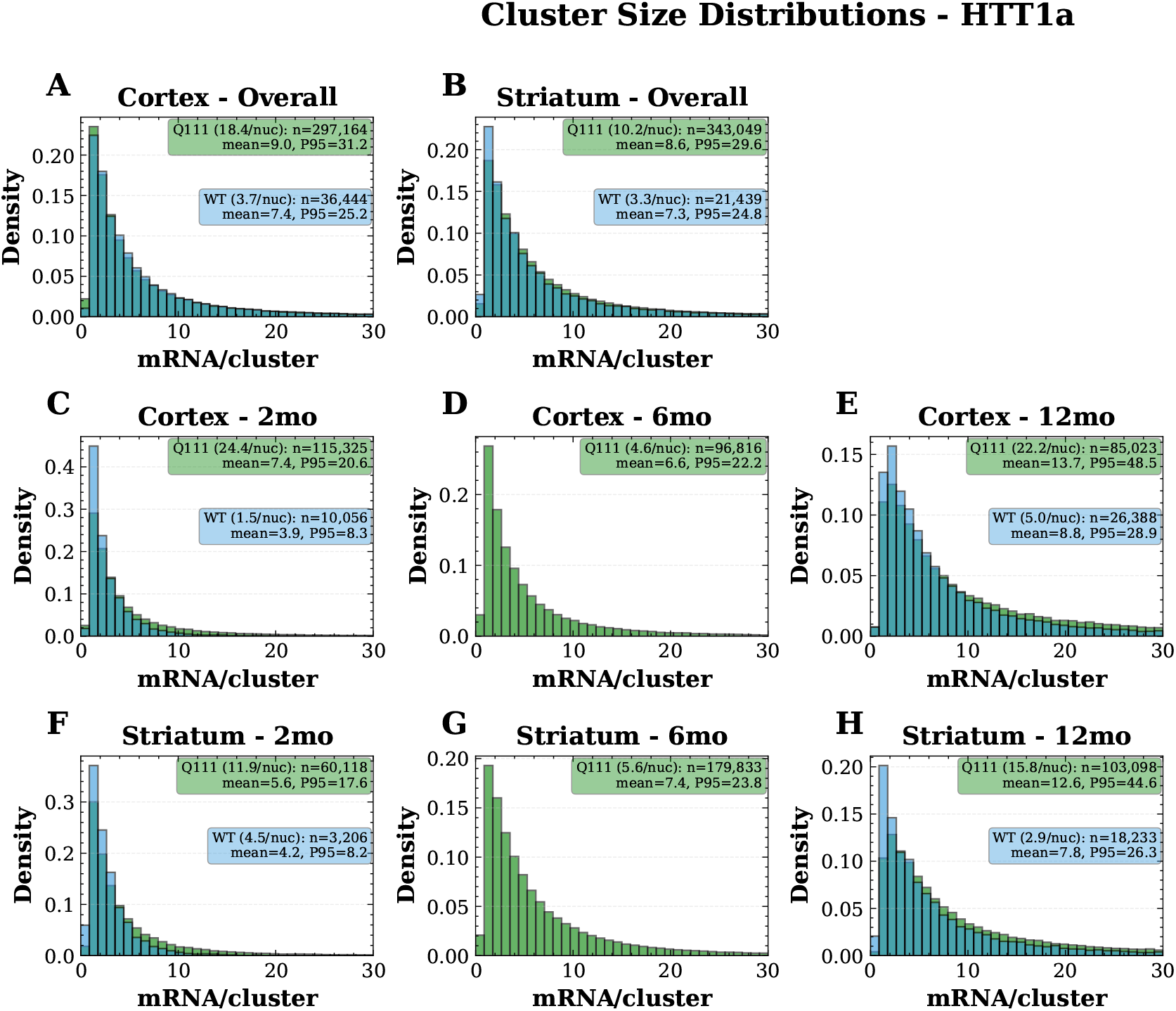
Cluster size distributions for HTT1a. Analysis of 698,169 clusters: Q111 640,320, WT 57,849. Cortex: 333,511 clusters; Striatum: 364,658 clusters. 13 slides excluded for technical failures. **(A-B) Overall distributions**. Cortex Q111: n=297,099 clusters, mean 8.96 mRNA equiv., median 3.87, P95 31.21, 18.42 clusters/nucleus; WT: n=36,412, mean 7.40, median 3.90, P95 25.06, 3.71 clusters/nucleus. Striatum Q111: n=342,971, mean 8.64, median 4.54, P95 29.57, 10.19 clusters/nucleus; WT: n=21,437, mean 7.26, median 3.98, P95 24.78, 3.35 clusters/nucleus. **(C-H) Age stratification**. Cortex: 2mo Q111 n=115,329, mean 7.44, 24.37 cl/nucleus; 6mo n=96,749, mean 6.59, 4.62 cl/nucleus; 12mo Q111 n=85,021, mean 13.74, 22.16 cl/nucleus. Striatum: 2mo Q111 n=60,096, mean 5.64, 11.94 cl/nucleus; 6mo n=179,781, mean 7.40, 5.64 cl/nucleus; 12mo Q111 n=103,094, mean 12.54, 15.79 cl/nucleus. **(I-P)** Per-mouse distributions revealing inter-individual variability. Cluster size = cluster intensity / slide-specific single-spot peak intensity (mRNA equivalents). CV threshold ≥0.5; intensity threshold >95th percentile of negative control.

**Fig. S16:**
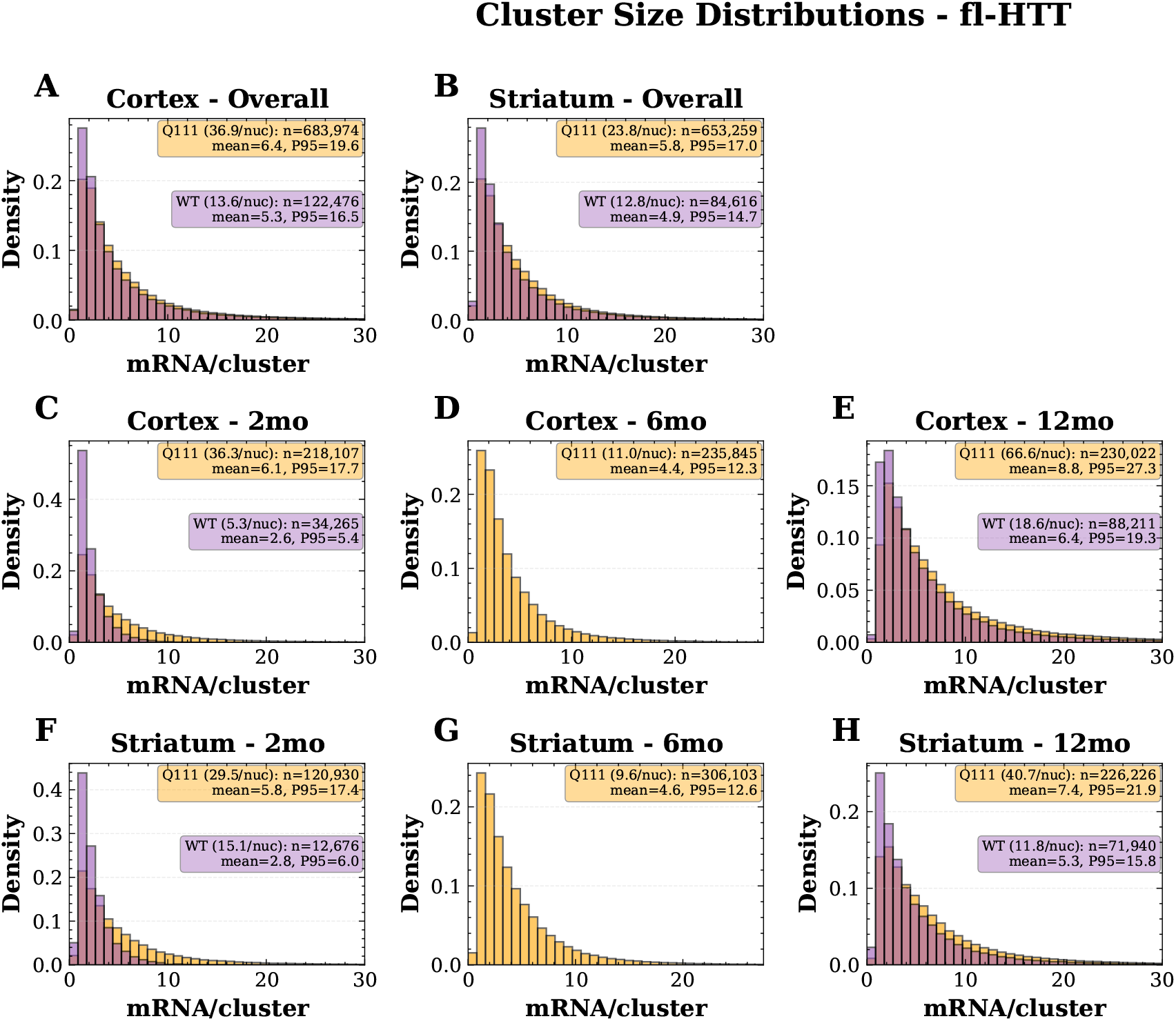
Cluster size distributions for fl-HTT. Analysis of 1,544,983 clusters: Q111 1,337,908, WT 207,075. Cortex: 806,405 clusters; Striatum: 738,578 clusters. 13 slides excluded for technical failures. **(A-B) Overall distributions**. Cortex Q111: n=683,947 clusters, mean 6.42 mRNA equiv., median 3.77, P95 19.59, 36.85 clusters/nucleus; WT: n=122,458, mean 5.30, median 3.10, P95 16.47, 13.56 clusters/nucleus. Striatum Q111: n=653,230, mean 5.78, median 3.75, P95 17.02, 23.75 clusters/nucleus; WT: n=84,617, mean 4.88, median 3.03, P95 14.68, 12.83 clusters/nucleus. **(C-H) Age stratification**. Cortex: 2mo Q111 n=218,032, mean 6.14, 36.24 cl/nucleus; 6mo n=235,892, mean 4.38, 11.03 cl/nucleus; 12mo Q111 n=230,023, mean 8.79, 66.57 cl/nucleus. Striatum: 2mo Q111 n=120,905, mean 5.83, 29.48 cl/nucleus; 6mo n=306,100, mean 4.57, 9.63 cl/nucleus; 12mo Q111 n=226,225, mean 7.39, 40.68 cl/nucleus. **(I-P)** Per-mouse distributions revealing inter-individual variability. Cluster size = cluster intensity / slide-specific single-spot peak intensity (mRNA equivalents). CV threshold ≥0.5; intensity threshold >95th percentile of negative control.

**Fig. S17:**
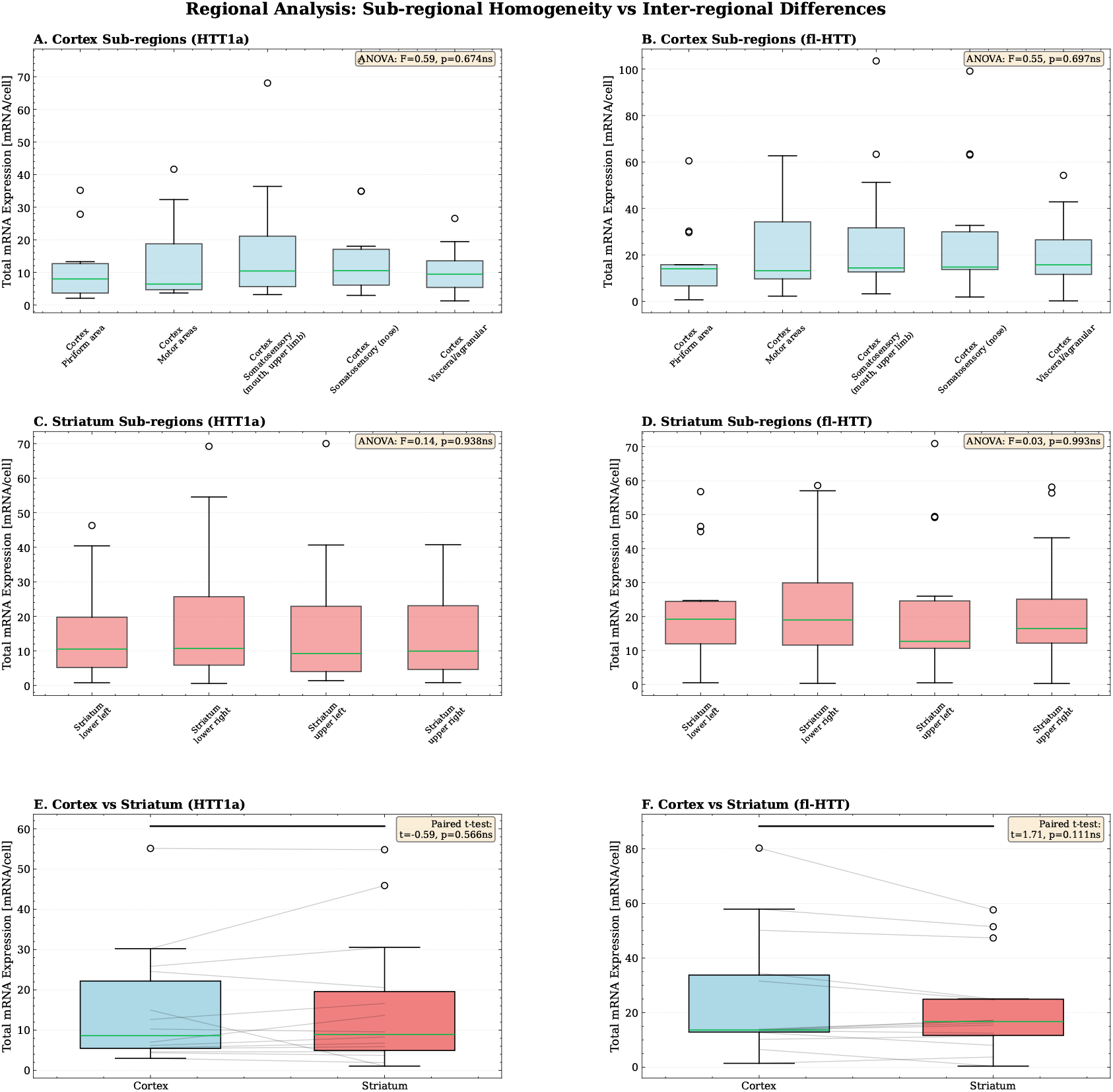
Detailed regional expression analysis. Sub-regional expression patterns analyzed using one-way ANOVA with Tukey HSD post-hoc tests (*α* = 0.05). **Dataset:** Q111 experimental samples, 15 slides, 2,668 FOVs total (Cortex: 1,394; Striatum: 1,274). Age range: 2-12 months; atlas coordinate range: 40-68 (sections spaced at 100 *µ*m intervals). **Cortex subregions (n=5):** Piriform area, Primary/secondary motor areas, Primary somatosensory (mouth/upper limb), Supplemental/primary somatosensory (nose), Visceral/gustatory/agranular areas. **Striatum subregions (n=4):** Lower left, lower right, upper left, upper right quadrants. **Row 1 (A-B):** Cortex sub-regional analysis. **ANOVA, HTT1a:** *F* = 0.579, *p* = 0.679 (ns). **ANOVA, fl-HTT:** *F* = 0.548, *p* = 0.702 (ns). **Row 2 (C-D):** Striatum sub-regional analysis. **ANOVA, HTT1a:** *F* = 0.135, *p* = 0.939 (ns). **ANOVA, fl-HTT:** *F* = 0.033, *p* = 0.992 (ns). **Row 3 (E-F):** Inter-regional comparison with paired t-test and connecting lines. HTT1a: Cortex 14.91 ± 13.77 vs Striatum 15.95 ± 15.90 mRNA/cell, *t* = −0.605, *p* = 0.555 (ns, n=14 paired samples). Full-length: Cortex 25.30 ± 21.98 vs Striatum 22.08 ± 17.11 mRNA/cell, *t* = 1.698, *p* = 0.113 (ns). **Conclusion:** No significant differences between subregions within cortex or striatum (all Tukey HSD *p* > 0.7), confirming uniform expression patterns within major brain regions. This validates pooling FOVs across subregions for region-level analyses.

**Fig. S18:**
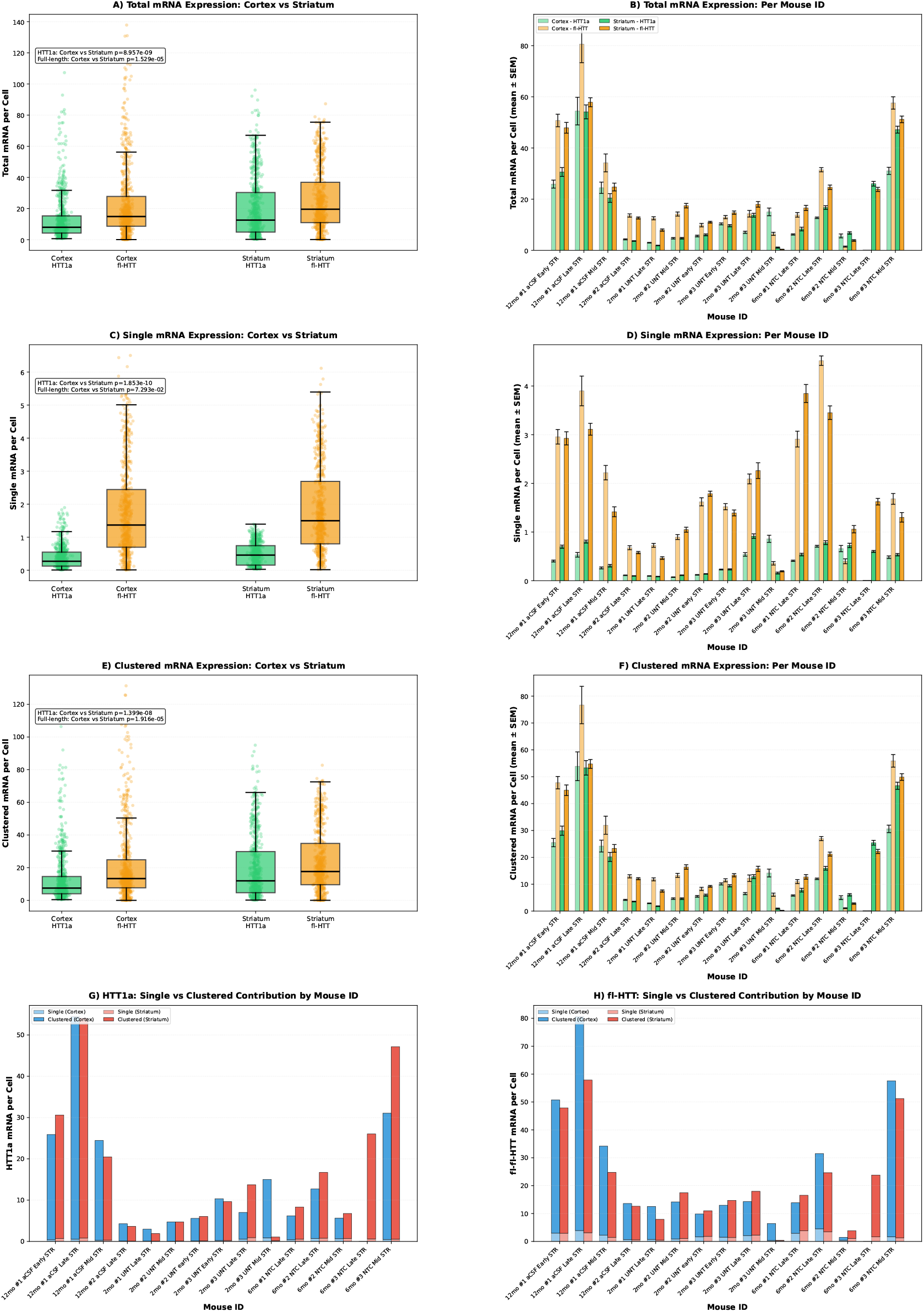
Expression analysis across Q111 mice. Detailed analysis of 2,668 FOVs from 15 Q111 slides. Cortex: 1,394 FOVs; Striatum: 1,274 FOVs. FOVs per mouse range from 86 to 250. **(A-B) Total mRNA**. Cortex: HTT1a mean 13.41 ± 14.88, median 7.98 mRNA/cell (n=697 FOVs); fl-HTT 22.80 ± 23.40, median 14.96. Striatum: HTT1a 19.96 ± 19.25, median 12.62; fl-HTT 25.37 ± 19.76, median 19.81. **(C-D) Single mRNA**. Cortex: HTT1a 0.39 ± 0.35, median 0.28; fl-HTT 1.77 ± 1.41, median 1.39. Striatum: HTT1a 0.49 ± 0.34, median 0.46; fl-HTT 1.84 ± 1.31, median 1.49. **(E-F) Clustered mRNA**. Cortex: HTT1a 13.02 ± 14.74, median 7.63; fl-HTT 21.03 ± 22.48, median 13.52. Striatum: HTT1a 19.48 ± 19.07, median 11.99; fl-HTT 23.53 ± 19.09, median 17.80. **Quality control:** 13 slides excluded for poor tissue integrity; CV≥0.5 for cluster filtering; minimum 40 nuclei per FOV.

**Fig. S19:**
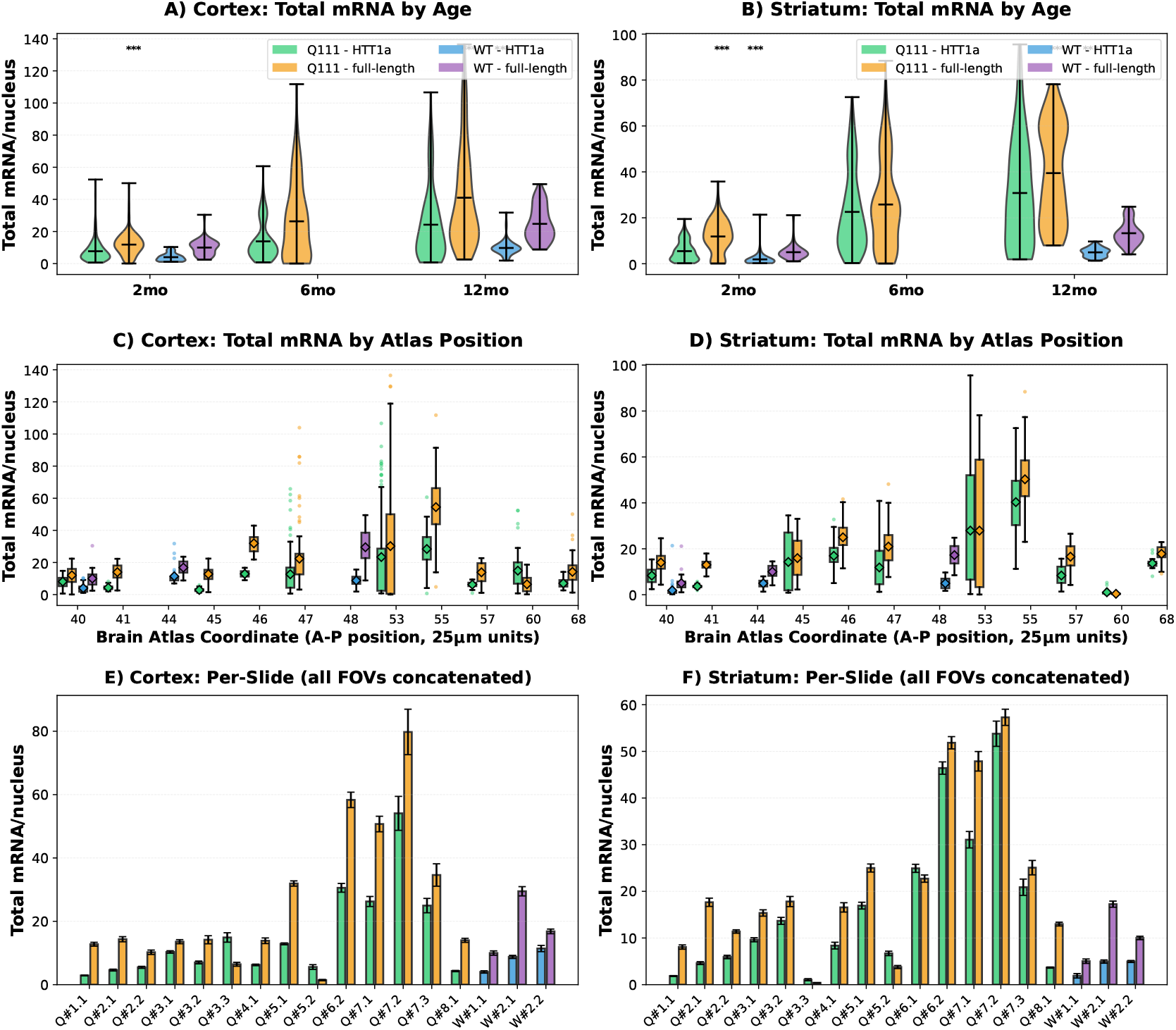
Comprehensive total mRNA expression analysis. Total mRNA combines single spots and clustered signal: (*N*_spots_ + *I*_cluster_*/I*_peak_)*/N*_nuclei_. Dataset: 3,234 FOVs (Q111: 2,668; WT: 566). **(A-B) Age trends**. Violin plots showing distribution of total mRNA expression. Cortex: 2mo Q111 HTT1a 7.83 ± 7.08 (n=335), WT 4.09 ± 2.62 (n=51); 6mo Q111 13.75 ± 11.31 (n=193); 12mo Q111 24.17 ± 22.36 (n=169), WT 9.73 ± 4.46 (n=83). Striatum: 2mo Q111 5.53 ± 4.35 (n=180), WT 1.93 ± 3.24 (n=45); 6mo Q111 22.50 ± 16.81 (n=284); 12mo Q111 30.78 ± 23.06 (n=173), WT 4.93 ± 1.96 (n=104). **FOV-level statistical tests (Q111 vs WT):** Striatum 2mo HTT1a: *t* = 5.20, *p* < 0.0001; Striatum 2mo fl-HTT: *t* = 6.36, *p* < 0.0001; Striatum 12mo HTT1a: *t* = 11.40, *p* < 0.0001; Striatum 12mo fl-HTT: *t* = 12.73, *p* < 0.0001. *Note:* FOV-level tests are more sensitive than per-slide tests but violate independence assumptions. **(C-D)** Atlas coordinate trends. Box plots showing expression across approximate anterior-posterior coordinates (atlas sections spaced at 100 *µ*m intervals). Q111 HTT1a: cortex 13.43 ± 14.94 (n=697), striatum 19.95 ± 19.20 (n=637). WT HTT1a: cortex 7.58 ± 4.73 (n=134), striatum 4.03 ± 2.78 (n=149). **(E-F)** Per-slide breakdown showing mean ± SEM for each slide (all FOVs concatenated per slide). **Power note:** With n = 4-6 mice per age group, per-slide analyses may be underpowered (see main text).

**Fig. S20:**
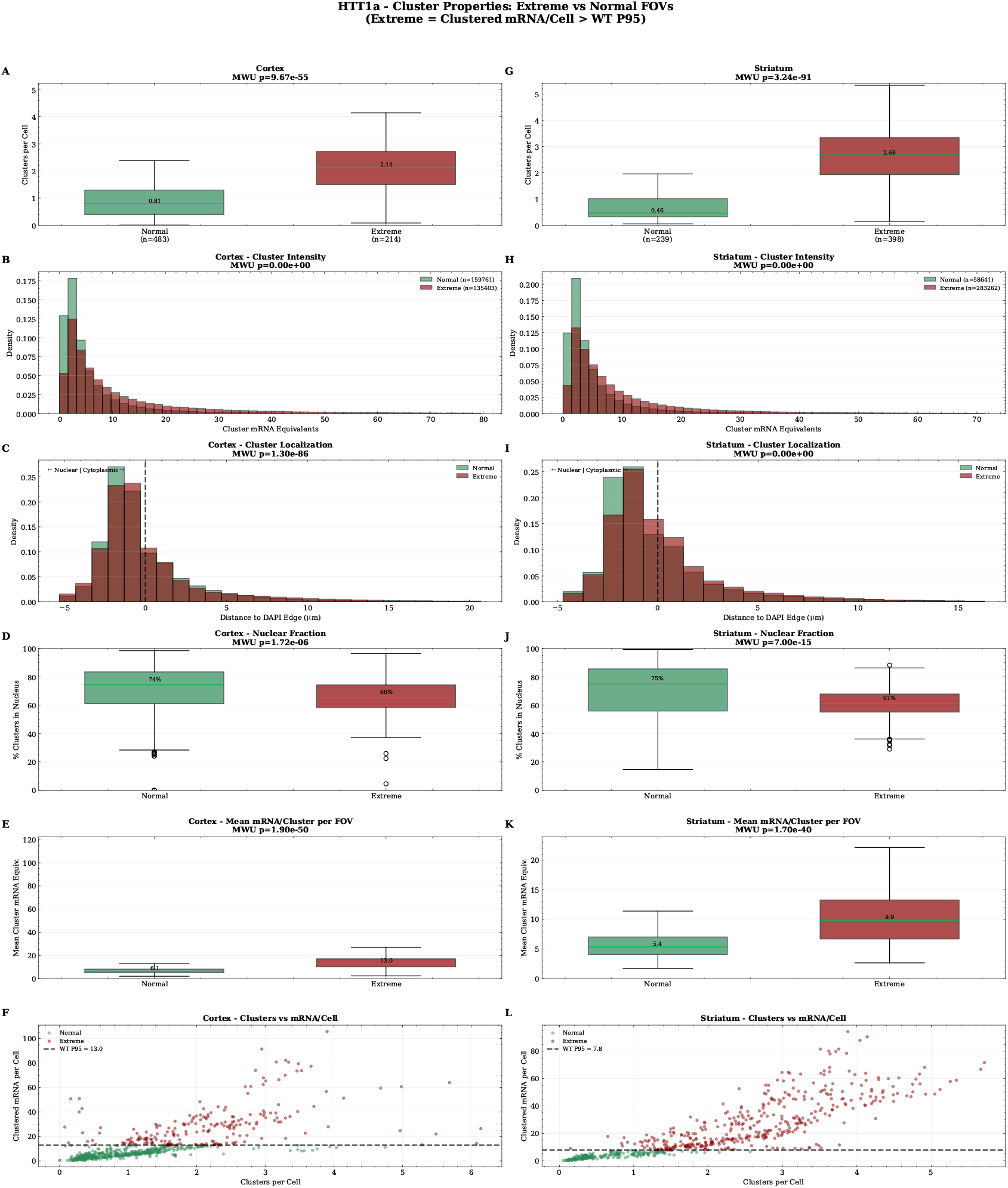
Cluster properties for HTT1a in extreme vs normal FOVs. Extreme FOVs defined as exceeding WT P95 threshold (Q111 tissue only; WT data only used to define threshold). **Quality control:** FOVs with <40 nuclei were excluded to ensure reliable per-nucleus normalization, removing 3,253 clusters (0.5%) relative to the unfiltered dataset in fig. S15. **Dataset:** Cortex: WT P95 threshold 12.87 mRNA/cell, Extreme n=217, Normal n=480. Striatum: threshold 7.77 mRNA/cell, Extreme n=399, Normal n=238. Total clusters after filtering: 637,067. Color scheme: Sea Green = Normal, Dark Red = Extreme. **(A-B) Clusters per cell**. Box plots comparing cluster abundance. Cortex: Extreme median 2.14 (IQR 1.24), Normal 0.81 (IQR 0.89), fold change 2.6 ×, Mann-Whitney *p* = 4.54 × 10^−54^. Striatum: Extreme 2.68 (IQR 1.39), Normal 0.46 (IQR 0.69), 5.9 ×, *p* = 2.07 × 10^−90^. **(C-D) Cluster mRNA equivalents**. Histograms of individual cluster sizes. Cortex: Extreme n=136,123 clusters, median 6.2 mRNA equiv.; Normal n=158,490, median 3.3; *p* < 10^−300^. Striatum: Extreme n=284,529, median 5.8; Normal n=57,992, median 3.0; *p* < 10^−300^. **(E-F) Nuclear localization**. Distribution of signed distance to DAPI edge (negative = nuclear). Cortex: Extreme median −0.91 *µ*m, 65.8% nuclear; Normal −1.06 *µ*m, 68.1% nuclear; *p* = 1.64 × 10^−81^. Striatum: Extreme −0.68 *µ*m, 61.6%; Normal −1.05 *µ*m, 67.7%; *p* < 10^−300^. **(G-H) Fraction nuclear per FOV**. Cortex: Extreme median 65.5%, Normal 74.2%, *p* = 2.70 × 10^−6^. Striatum: Extreme 61.2%, Normal 75.1%, *p* = 5.19 × 10^−15^. **(I-J) Mean cluster mRNA**. Cortex: Extreme 12.9, Normal 6.1, *p* = 5.63 × 10^−51^. Striatum: Extreme 9.8, Normal 5.4, *p* = 8.86 × 10^−41^. **(K-L)** Scatter plots: clusters/cell vs mRNA/cell with WT P95 threshold (dashed line). **Cluster detection:** DBSCAN (eps=0.75 *µ*m, min samples=3), intensity > 2.5× peak.

**Fig. S21:**
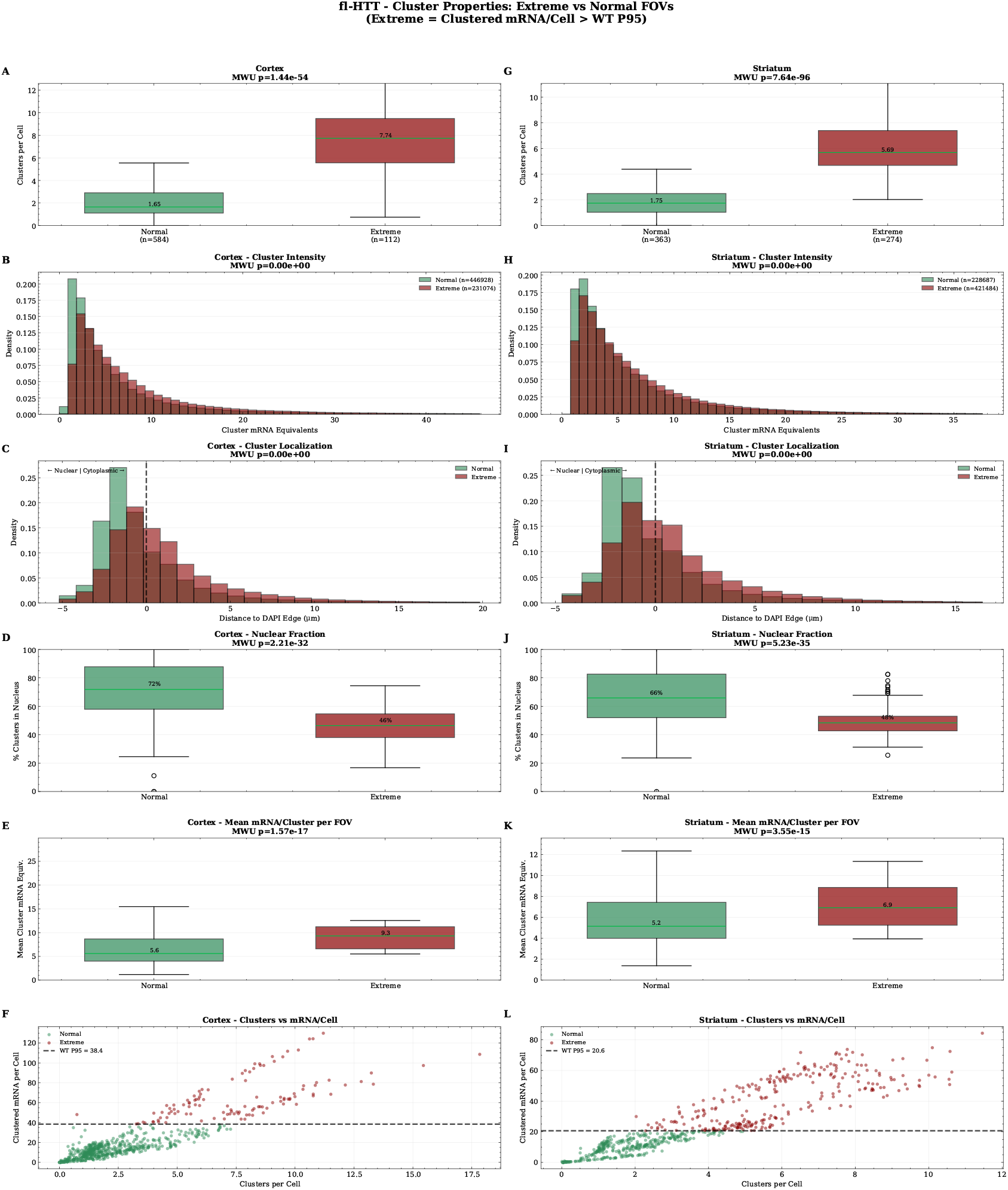
Cluster properties for fl-HTT in extreme vs normal FOVs. Quality control: FOVs with <40 nuclei were excluded to ensure reliable per-nucleus normalization, removing 9,735 clusters (0.7%) relative to the unfiltered dataset in fig. S16. **Dataset:** Cortex: WT P95 threshold 38.00 mRNA/cell, Extreme n=112, Normal n=585. Striatum: threshold 20.41 mRNA/cell, Extreme n=278, Normal n=359. Total clusters after filtering: 1,328,173. **(A-B) Clusters per cell**. Cortex: Extreme median 7.67, Normal 1.64, 4.7×, *p* = 2.24 × 10^−55^. Striatum: Extreme 5.69, Normal 1.76, 3.2×, *p* = 2.80 × 10^−97^. **(C-D) Cluster mRNA equivalents**. Cortex: Extreme n=232,925 clusters, median 5.4; Normal n=445,343, median 3.8. Striatum: Extreme n=424,127, median 4.7; Normal n=225,177, median 3.9. **(E-F) Nuclear localization (notably stronger shift)**. Cortex: Extreme median +0.34 *µ*m (cytoplasmic), 45.4% nuclear; Normal −1.12 *µ*m, 67.8% nuclear; *p* < 10^−300^. Striatum: Extreme +0.12 *µ*m, 48.7%; Normal −1.08 *µ*m, 68.0%. **(G-H) Fraction nuclear per FOV**. Cortex: Extreme 46.4%, Normal 71.8%, *p* = 1.69 × 10^−32^ (25.4 pp difference). Striatum: Extreme 48.5%, Normal 66.0%, *p* = 2.12 × 10^−34^. **(I-J) Mean cluster mRNA**. Cortex: Extreme 9.5, Normal 5.6, *p* = 1.68 × 10^−17^. Striatum: Extreme 6.7, Normal 5.1. **Key finding:** Extreme FOVs show marked cytoplasmic accumulation (∼50% nuclear vs ∼68% in normal), consistent with impaired nuclear export or cytoplasmic aggregation.

**Fig. S22:**
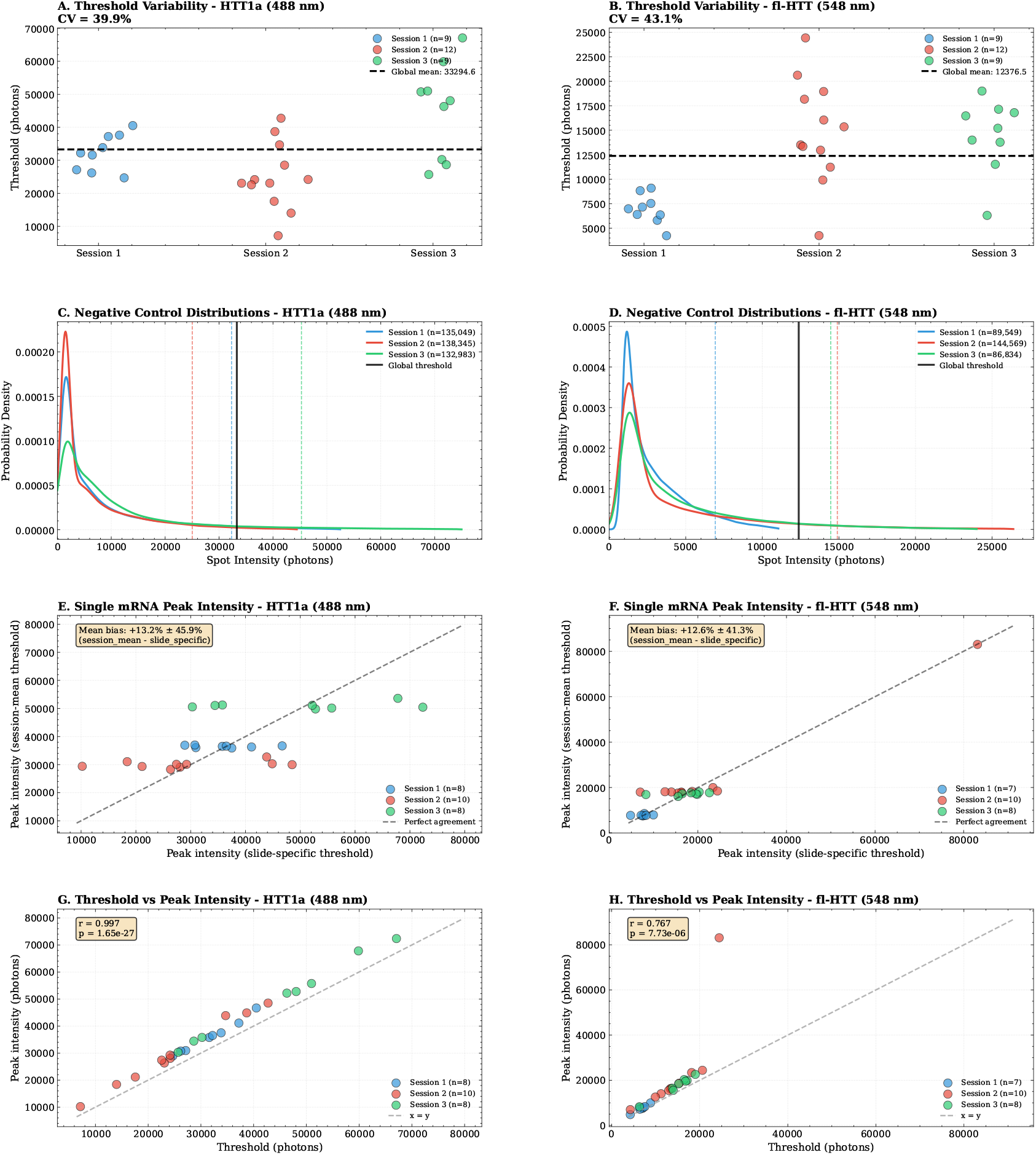
Importance of per-slide negative control calibration. Analysis across 3 imaging sessions, 30 slides with negative control, 51 slides total. **Threshold variability:** HTT1a (488 nm): global threshold 33,233 ± 13,474 A.U. (CV = 40.5%); Session 1: 32,143 ± 5,005 (n=9), Session 2: 24,961 ± 10,002 (n=12), Session 3: 45,352 ± 15,045 (n=9). fl-HTT (548 nm): 12,352 ± 5,301 A.U. (CV = 42.9%). **Row 1 (A-B):** Per-slide thresholds by session. Each point = one slide’s 95^th^ percentile of negative control. Dashed line = global mean. **Row 2 (C-D):** KDE of negative control spot intensities by session. **Row 3 (E-F):** Single mRNA peak intensity: slide-specific vs pooled threshold. Mean difference (pooled − slide-specific): HTT1a +13.6% ± 46.7%; fl-HTT +12.2% ± 40.6%. Session 2 shows largest difference (+23.1% for HTT1a). **Row 4 (G-H):** Threshold vs peak intensity correlation showing relationship between background and signal levels. **Conclusion:** Per-slide calibration essential for accurate quantification; pooled thresholds introduce variability in peak intensity estimation.

**Fig. S23:**
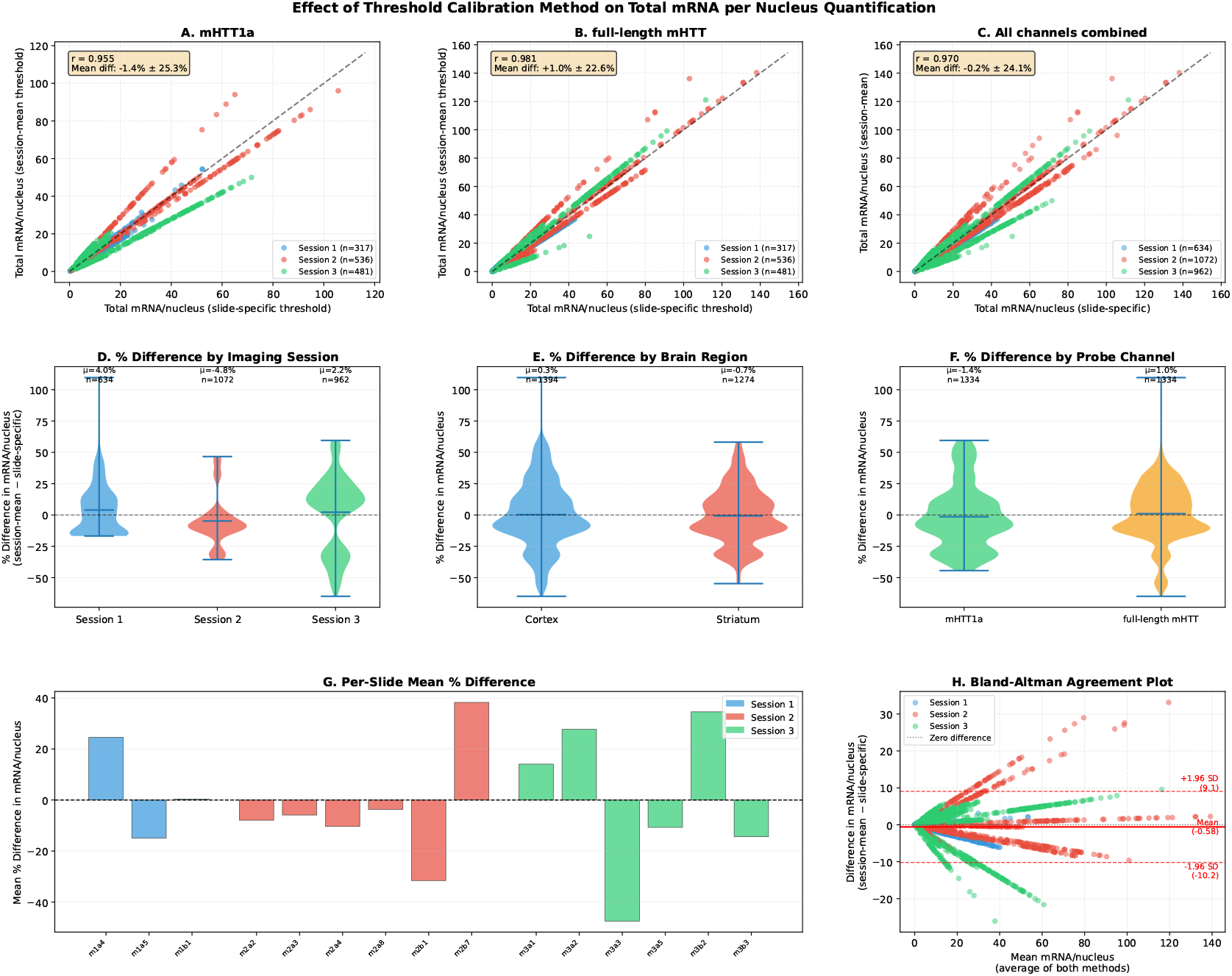
Effect of threshold calibration method on total mRNA quantification. Comparison of slide-specific (95th percentile per slide) vs pooled (average across slides within imaging batch) thresholds. **Dataset:** 2,668 FOVs (Session 1: n=634, Session 2: n=1,072, Session 3: n=962); Brain regions: Cortex n=1,394, Striatum n=1,274; Channels: HTT1a n=1,334, full-length n=1,334. 13 slides excluded for poor tissue integrity. **Agreement:** Pearson *r* = 0.9667 (*p* < 10^−100^), indicating near-perfect linear agreement; mean % difference 0%±25%; median −2.55% (IQR 30.08%). Paired t-test: *t* = 6.391, *p* = 1.94 × 10^−10^; Cohen’s *d* = −0.003 (negligible effect size). **Bland-Altman:** Mean difference −0.647 mRNA/nucleus; 95% limits of agreement [−10.89, +9.59] mRNA/nucleus. **Individual variability (CRITICAL):** 69.1% of FOVs differ >10%; 35.9% differ >20%; 5.2% differ >50%. Maximum individual difference: 98.2% (range −65.1% to +98.2%). 5th-95th percentile range: −33.9% to +45.3%. **Per-session breakdown:** Session 1: +4% ± 19%; Session 2: −5% ± 20%; Session 3: +2% ± 31%. **Per-slide variability:** 11/15 slides (73.3%) show >10% mean bias; 6/15 (40.0%) show >20% mean bias; max per-slide bias 48.2% (slide m3a3). **Row 1 (A-C):** Scatter plots comparing slide-specific (x) vs pooled (y) mRNA quantification by channel. Dashed line = perfect agreement. **Row 2 (D-F):** Violin plots of % differences by session, region, and channel. **Row 3 (G-H):** Per-slide mean difference (bar plot) and Bland-Altman plot (agreement analysis). **Conclusion:** While mean difference is negligible (−0.07%), individual FOVs show substantial variability (up to 98% difference); slide-specific calibration strongly recommended for publication-quality data, especially when comparing samples across imaging batches.

**Fig. S24:**
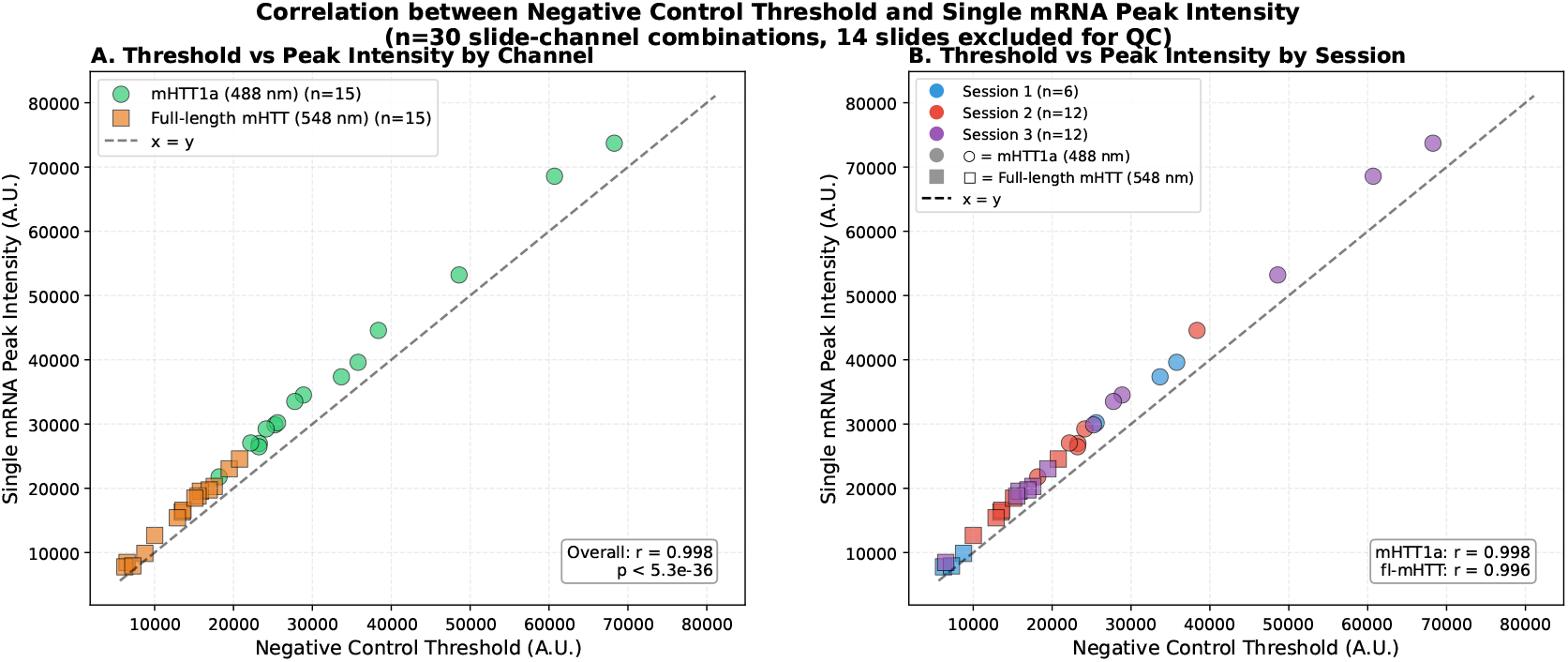
Correlation between threshold and peak intensity. Technical validation showing relationship between negative control thresholds (95th percentile of DapB intensities) and single-molecule peak intensities (KDE mode of experimental spot intensities) across slides. These metrics are independently derived from different probe populations (negative control vs experimental), providing internal consistency check. **Interpretation:** Positive correlation indicates imaging conditions that elevate background (autofluorescence, detector settings) also elevate experimental signal. This is expected for technical factors affecting both uniformly (laser power, objective alignment, tissue autofluorescence). **Implication:** Per-slide normalization accounts for correlated technical variation in both threshold and peak intensity, improving accuracy of mRNA quantification. Session-level or global normalization fails to capture slide-specific technical effects.

**Fig. S25:**
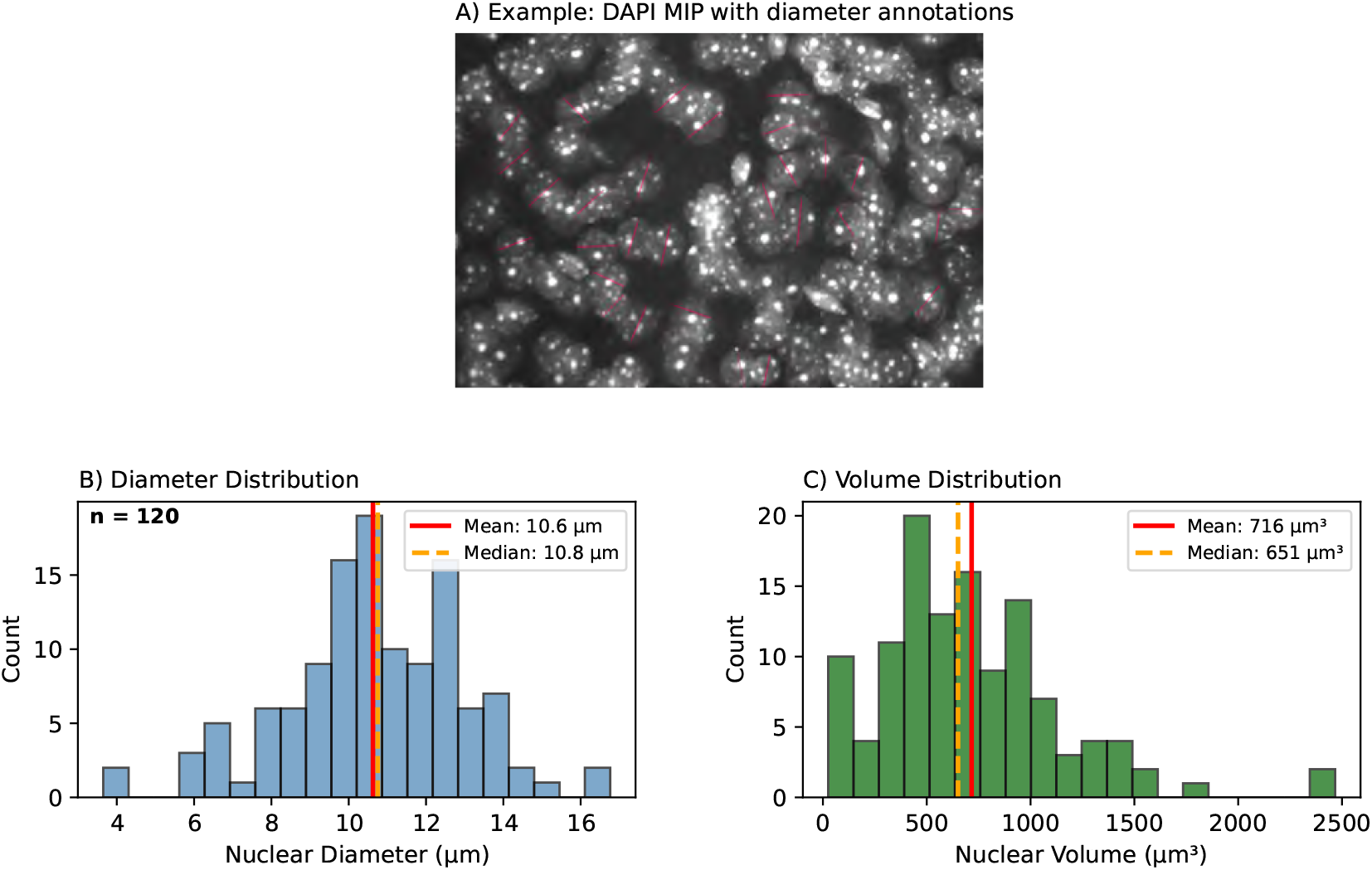
Nuclear diameter calibration for per-nucleus mRNA quantification. Manual measurement of nuclear diameters from DAPI-stained mouse brain tissue to establish mean nuclear volume for cell count estimation. **(A)** Representative maximum intensity projection (MIP) of DAPI-stained tissue with manual diameter annotations (red lines). Nuclei were selected to span cortical and striatal regions across multiple FOVs. **(B)** Distribution of measured nuclear diameters (n = 120 nuclei). Mean ± SEM = 10.6 ± 0.2 *µ*m (SD = 2.3 *µ*m); median = 10.8 *µ*m. Range spans 3.6-16.8 *µ*m, reflecting natural variation in nuclear size across cell types and tissue regions. **(C)** Derived nuclear volume distribution assuming spherical geometry 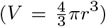. Mean ± SEM = 716 ± 39 *µ*m^3^ (SD = 432 *µ*m^3^); median = 651 *µ*m^3^. The mean value (716 *µ*m^3^) is used throughout this study for converting total DAPI volume to estimated nuclear counts. *Rationale:* The spherical approximation, while simplifying actual ellipsoidal nuclear morphology, provides consistent estimates across tissue regions. Deviations from sphericity average out across the many nuclei per FOV (typically 50-200), and the resulting per-nucleus mRNA values show strong correlation with direct nuclear counts (*r* = 0.94; see Supplementary Note 2).

### Supplementary Note 1: Per-slide versus pooled negative control calibration

Including a dedicated negative control section on every experimental slide increases reagent costs and uses tissue sections that could otherwise be used for experimental imaging. A practical alternative is to image negative controls on only a subset of slides and apply a pooled threshold to all slides. Here we evaluate the trade-offs of this approach. In our experimental design, each slide contains tissue sections from a single animal, with the negative control (DapB probe) applied to a separate section on the same slide as the experimental probes. Imaging proceeds slide-by-slide, with all fields of view from one slide acquired before moving to the next. This design means that within-slide variation reflects local factors such as tissue properties and staining uniformity, while between-slide variation could reflect either technical factors (imaging drift, illumination changes) or biological differences between animals. We compared per-slide thresholds (95th percentile of DapB spot intensities) against pooled thresholds for 30 slides spanning 3 imaging sessions on separate days. Between-session variation was substantial: Session 3 thresholds were ∼1.8× higher than Session 2 for the green channel (fig. S22). This magnitude of difference clearly reflects technical variation and demonstrates that per-session calibration is essential. Within-session variation was also considerable (CV ∼15-33%), but we cannot definitively determine whether this reflects technical drift during the session or biological animal-to-animal differences in tissue autofluorescence. We lack ground truth to distinguish these sources. Figure S14 (panels M-O) shows per-animal threshold distributions. The absence of significant regional differences within slides (panels A-C, *p* = 0.22) and lack of age effects (panels D-F) suggests the threshold variation occurs at the slide level rather than reflecting systematic biological gradients.

The negative control threshold directly affects the single-molecule peak intensity estimate: after applying the threshold, only spots above this cutoff contribute to the intensity distribution, and the mode of this distribution defines the single-molecule reference intensity. As shown in fig. S24 (panels A-C), there is a positive linear correlation between threshold and peak intensity across slides. This relationship is expected–a higher threshold excludes more dim spots, shifting the intensity distribution toward brighter values. When using pooled thresholds instead of per-slide values, individual slides show mean differences of +13.6% ± 46.7% in peak intensity for mHTT1a and +12.2% ± 40.6% for fl-mHTT (fig. S22, rows E-F).

Figure S23 (panels A-C) directly compares total mRNA per nucleus calculated using slide-specific versus pooled thresholds across 2,668 FOVs. The scatter plots show a linear relationship (*r* = 0.967) with negligible mean difference ( −0.07%), indicating that for population-level comparisons, the two calibration methods yield similar results. However, individual FOV-level variability is substantial (36% differ by >20%), and Bland-Altman analysis (panel H) shows 95% limits of agreement spanning ±10 mRNA/nucleus.

We cannot prove that per-slide calibration is more accurate than session-pooled calibration. If the within-session threshold variation is primarily biological, then per-slide calibration might over-correct by attributing biological variation to technical noise. Conversely, if the variation is primarily technical, per-slide calibration appropriately accounts for slide-to-slide drift. For the present study, we used per-slide calibration as a conservative approach: per-session calibration is clearly required given the 1.8 × between-session differences, and per-slide calibration cannot make results worse if the variation is technical. However, for applications where absolute quantification matters less than relative comparisons within a single session, pooled calibration may be adequate and offers cost savings by reducing the number of negative control sections required.

### Supplementary Methods: Model Architecture and Training

The identification of mRNA clusters in the 3D microscopy images was performed using a deep learning approach, specifically employing a residual 3D U-Net architecture based on an existing, publicly available implementation [45]. The selected model, *ResidualUNet3D*, extends the classical 3D U-Net by integrating residual connections within its convolutional blocks, enabling deeper architectures and improved convergence. The network architecture is characterized by:

- An encoder-decoder structure with skip connections to preserve spatial context.
- Residual blocks consisting of Group Normalization, Convolution, and ReLU activations arranged in the order: GroupNorm → Conv3D → ReLU.
- A sequence of four hierarchical layers with increasing feature channels (32, 64, 128, 256) to capture multi-scale spatial features.
- A sigmoid activation applied at the output layer to produce voxel-wise probability maps of mRNA clusters.

Model training was configured as follows:

- **Loss Function**: Binary cross-entropy loss with logits (*BCEWithLogitsLoss*).
- **Optimizer**: Adam optimizer with a learning rate of 2 ×10^−4^ and weight decay of 10^−5^.
- **Learning Rate Scheduler**: A plateau-based scheduler (*ReduceLROnPlateau*) was employed, reducing the learning rate by a factor of 0.2 after no improvement over eight validation cycles.
- **Training Protocol**: Training was performed for a maximum of 800 epochs (15,000 iterations), with validation every 500 iterations and TensorBoard logging every 30 iterations.

#### Data Preprocessing and Augmentation

The input training data consisted of volumetric patches (8 × 256 × 256 voxels), which were normalized (standardization) and augmented to enhance generalizability. Augmentation included:

- Random flipping and 90-degree rotations.
- Random rotations ( ±30^°^), specifically in the ZY-plane, to make the model robust for different spacing between z-slices.
- Elastic deformation for realistic shape variability.

#### Inference and Post-processing

At inference, the trained residual U-Net produced probability maps highlighting potential mRNA clusters. The predicted clusters were thresholded (set at a probability of 0.5) to yield binary masks.

### Supplementary Note 2: Mean nuclear volume estimation

Converting total mRNA expression to per-nucleus values requires estimating the number of nuclei per field of view. Direct counting of individual nuclei via instance segmentation is challenging in brain tissue due to the densely packed cellular architecture: overlapping and touching nuclei (particularly prevalent in striatum and deep cortical layers) cause standard segmentation algorithms (watershed, deep learning-based instance segmentation) to either merge adjacent nuclei into single objects or fragment single nuclei into multiple objects. Rather than attempting unreliable direct counting, we estimate nuclear number by dividing the total DAPI-stained volume by a mean nuclear volume derived from manual diameter measurements (fig. S25).

We manually measured nuclear diameters from DAPI maximum intensity projections, selecting well-separated nuclei across cortical and striatal regions (n = 120 nuclei). The measured diameter distribution showed mean ±SEM of 10.6 ± 0.2 *µ*m (SD = 2.3 *µ*m; median = 10.8 *µ*m; range 3.6-16.8 *µ*m; fig. S25B). Assuming spherical geometry 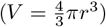, this yields a mean nuclear volume of 716 ± 39 *µ*m^3^ (SD = 432 *µ*m^3^; median = 651 *µ*m^3^; fig. S25C). We use the mean value (716 *µ*m^3^) throughout this study for converting total DAPI volume to estimated nuclear counts. This nuclear diameter is consistent with published measurements for mouse striatal neurons. Medium spiny neurons (MSNs), which constitute >95% of striatal neurons, have nuclear diameters of 10-11 *µ*m in adult mice [51]. We assume spherical nuclear geometry for volume estimation. Other cell types (interneurons, glia) may have different nuclear sizes, making this volume-based approach less accurate than individual nuclear segmentation. However, since we apply the same mean nuclear volume across all conditions, relative comparisons between genotypes and brain regions remain valid.

**Table S6:** RNAscope reagent list. Complete list of reagents used for RNAscope™ experiments, including reference numbers and storage conditions.

| Reagent | Ref./Cat. No. | Qty | Comments |
| --- | --- | --- | --- |
| *Stored at 4° C* |  |  |  |
| RNAscope H<sub>2</sub>O<sub>2</sub> and Protease Reagents | 322381 | 2 | 3 mL squeeze-dropper bottles |
| Hydrogen Peroxide | 322335 | 2 | - |
| Protease III | 322337 | 2 | For prefixed tissues |
| Protease IV | 322336 | 2 | For fresh frozen tissues |
| *RNAscope Probes* |  |  |  |
| 3-plex Positive Control Probe - Mm | 320881 | 1 | C1: ~29 μL/slice |
| 3-plex Negative Control Probe | 320871 | 1 | - |
| Mm-Htt-intron1-O2 (HTT1a) | 575581 | 1 | Mouse HTT intron 1 |
| Mm-Htt-No-XHs-C3 (fl-HTT) | 473001-C3 | 3 | Full-length mouse HTT |
| Mm-Pp1r1b-C2 (DARPP-32) | 405901-C2 | 1 | Striatal marker |
| Hs-HTT-intron1-No-XMm-O1-C2 | 561441-C2 | 3 | Human HTT intron 1 |
| HS-HTT-no-X-MmOa-ver2 | 420231 | 3 | Full-length human HTT |
| *RNAscope V2 Fluorophores* |  |  |  |
| Alexa Fluor 488 tyramide reagent | B40953 | 2 | - |
| Alexa Fluor 546 tyramide reagent | B40955 | 2 | - |
| Alexa Fluor 647 tyramide reagent | B40958 | 2 | - |
| RNAscope Multiplex TSA buffer | 322809 | 1 | - |
| *RNAscope Multiplex Fluorescent Detection v2* |  |  |  |
| Detection Reagents v2 (Box) | 323110 | 4 | 3 mL squeeze-dropper bottles |
| AMP 1 | 323101 | 1 | - |
| AMP 2 | 323102 | 1 | - |
| AMP 3 | 323103 | 1 | - |
| HRP C1 | 323104 | 1 | - |
| HRP C2 | 323105 | 1 | - |
| HRP C3 | 323106 | 1 | - |
| HRP Blocker | 323107 | 3 | - |
| DAPI | 323108 | 1 | - |
| *Stored at Room Temperature* |  |  |  |
| RNAscope Wash Buffer (Box) | 310091 | 4 | 4 bottles per box |
| RNAscope Target Retrieval (Box) | 322000 | 8 | 4 bottles per box |
| ProLong Glass Antifade Mountant | P36984 | 2 | - |
| DMSO | J66650.AD | - | - |
| Ethanol | BP2818-4 | - | - |
| SSC Buffer 20× | S6639-1L | - | - |
| 10% Neutral Buffered Formalin | - | - | - |
| *Other Materials* |  |  |  |
| High precision coverslips #1.5 | CG15KH1 | 1 | - |
| Hydrophobic barrier pen | H-4000 | 1 | - |

### Supplementary Note 3: RNAscope Protocol for Prefixed Mouse Brain Tissues

The following protocol was used for RNAscope™ staining of mouse brain tissues that had been prefixed for more than 48 hours in PFA/NBF.

#### Section Preparation

- Set the oven to 60^°^C.
- Prepare a petri dish with deionized water (DI water) and thin brushes for transferring brain slices.
- Using a thin brush, move the brain slice from the plate into the petri dish. Gently unfold the brain slice as much as possible.
- Pipette 100 *µ*L of DI water onto the slide for mounting. Treat the slide as three sections: top, middle, and bottom, large enough to fit 3 brain samples.
- Carefully transfer the brain slice using the thin brush to move it into the water droplet on the slide.
- Continue to unfold the brain slice while gently wiping excess water with a Kimwipe until the sample is completely flat. Avoid directly touching the sample with Kimwipe.
- Using the thin brush, orient the brain slices in the same manner with image processing purposes in mind.
- Bake the slides at 60^°^C for 30 min in the humidity tray (maximum 20 slides) with the tray lid off and no wet paper towels. After incubation, adjust the temperature to 40^°^C.
- During this incubation, prepare 1× wash buffer by mixing 60 mL of 50× RNAscope wash buffer (Ref. No. 320058) with 2.94 L milli-Q water.

#### Fixation and Dehydration

1. Place prefixed tissues into the slide rack and immerse slides in 10% Neutral Buffer Formalin (NBF). Place the container on ice for 15 min.
2. Prepare ethanol standards:
3. Immerse slides in 50% EtOH for 5 min at room temperature (RT).
4. Immerse slides in 70% EtOH for 5 min at RT.
5. Immerse slides in 100% EtOH for 5 min at RT. Repeat with fresh 100% EtOH.
6. Remove excess liquid by gently flicking slides and wiping around the sample. Allow to dry for 5 min at RT.

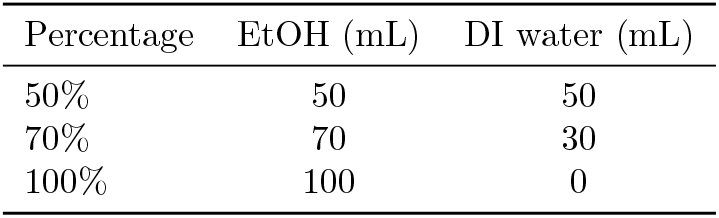

#### Pretreatment

1. Add 5 drops of hydrogen peroxide (Ref. No. 322335) to cover each sample and incubate at RT for 10 min.
2. Prepare 1 ×Target Retrieval Buffer by mixing 70 mL of 10× Target Retrieval Buffer (Ref. No. 322001) with 700 mL DI water in a 1 L glass beaker and boil on a hot plate.
3. Wash slides in DI water by moving the slide rack up and down 5 times. Repeat with fresh DI water.

#### Target Retrieval

1. Submerge slides in boiling liquid for 5 min, ensuring the temperature is >99^°^C and <105^°^C.
2. Immediately transfer hot slide rack into RT DI water.
3. Wash slides in DI water (5 times). Repeat with fresh DI water.
4. Wash slides in 100% EtOH (5 times).
5. Remove excess liquid and allow to dry for 5 min at RT.
6. Draw 1-2 circles with hydrophobic barrier pen (Cat. No. H-4000) around each sample without touching it. Allow to dry for 5 min.
7. Insert slides into humidity tray.
8. Add 1 drop of Protease III to cover each section.
9. Incubate at 40^°^C for 30 min.
10. During incubation, dilute probes to RT before use:
11. Wash slides in DI water (5 times). Repeat with fresh DI water.

| # Experiments | C1 (μL) | C2 (μL) | C3 (μL) |
| --- | --- | --- | --- |
| 6 | 240 | 5 | 5 |
| 10 | 480 | 10 | 10 |
| 20 | 960 | 20 | 20 |

#### Probe Hybridization

1. Remove excess liquid and insert slides into humidity tray.
2. For negative control probe (Ref. No. 320871) and positive control probe (Ref. No. 320881), add 1 drop to cover corresponding samples. For experimental probe mixture, pipette 30 *µ*L to cover samples.
3. Incubate in humidity tray for 2 hours at 40^°^C.
4. Submerge in wash buffer for 2 min at RT, agitating slightly. Repeat with fresh wash buffer.
5. Transfer slides to slide rack and submerge in 5× SSC (Ref. No. S6639-1L) overnight. Prepare by adding 25 mL of 20× SSC to 75 mL DI water.

#### Signal Amplification (Amp 1-3)

1. Line humidity tray with paper towels, fill with DI water, pour out excess, and place into oven at 40^°^C.
2. Transfer slides to humidity tray.
3. Wash off 5× SSC by submerging in wash buffer for 2 min at RT. Repeat.
4. Remove excess liquid.
5. Add 1 drop of Amp 1 (Ref. No. 323101) to cover each sample and incubate at 40^°^C for 30 min.
6. Wash in wash buffer for 2 min. Repeat.
7. Add 1 drop of Amp 2 (Ref. No. 323102) and incubate at 40^°^C for 30 min.
8. Wash in wash buffer for 2 min. Repeat.
9. Add 1 drop of Amp 3 (Ref. No. 323103) and incubate at 40^°^C for 15 min.
10. Wash in wash buffer for 2 min. Repeat.

#### HRP-C1 Signal Development

1. Add 1 drop of HRP-C1 (Ref. No. 323104) to cover each sample.
2. Incubate at 40^°^C for 15 min.
3. During incubation, prepare fluorophores: resuspend Alexa Fluor reagents (dry pellet) in 150 *µ*L DMSO (Ref. No. J66650.AD), then dilute 1:100 in TSA buffer (Ref. No. 322809):
4. Wash in wash buffer for 2 min. Repeat.
5. Add 30 *µ*L of corresponding diluted Alexa dye to cover each sample.
6. Incubate at 40^°^C for 30 min.
7. Wash in wash buffer for 2 min. Repeat.
8. Add 1 drop of v2 HRP blocker (Ref. No. 323107) to cover each sample.
9. Incubate at 40^°^C for 15 min.
10. Wash in wash buffer for 2 min. Repeat.

| # Slides | Alexa Fluor (μL) | TSA Buffer (mL) |
| --- | --- | --- |
| 6 | 10 | 0.99 |
| 10 | 20 | 1.98 |
| 20 | 40 | 3.96 |

#### HRP-C2 and HRP-C3 Signal Development

Repeat the HRP-C1 development steps for HRP-C2 (Ref. No. 323105) and HRP-C3 (Ref. No. 323106), using the corresponding Alexa dyes for each channel. For HRP-C3, retrieve ProLong Glass Antifade reagent (Ref. No. P36984) and thaw at RT for 1 hour before use to prevent bubbles.

#### Mounting

1. Add 1 drop of DAPI (Ref. No. 323108) to cover each sample.
2. Incubate for 10-60 min at RT (finalized time TBD).
3. Wash slides in 1 × PBS (5 times). Repeat with fresh PBS.
4. Remove excess liquid and allow samples to dry for 5 min at RT.
5. Set pipette to 60 *µ*L and slowly draw up ProLong Glass Antifade reagent (Cat. No. P36984). Slowly add reagent on each sample, avoiding bubbles.
6. Carefully place a #1.5 coverslip (Ref. No. CG15KH1) on each slide while avoiding bubbles.
7. Allow mounting reagent to harden overnight in the dark at RT, then seal with nail polish.
8. Store at 4^°^C until imaging. Samples are most stable for 2 weeks.

### Supplementary Note 4: RNAscope Probe Target Sequences

Custom RNAscope probes were designed by Advanced Cell Diagnostics (ACD Bio) for detection of mouse huntingtin (*Htt*) transcripts. Each probe consists of 20 ZZ pairs, with each ZZ pair covering approximately 40-50 nt, spanning roughly 1,000 nt total in the target region.

### fl-HTT Probe: Mm-Htt-No-XHs (Cat. No. 473001)

This probe targets the 3’ untranslated region (3’ UTR) of the canonical full-length mouse *Htt* mRNA. It consists of 20 ZZ pairs targeting region 9833-10798 of NM 010414.2 (966 bp).

#### Target sequence

>NM 010414.2:9833-10798 Mus musculus huntingtin (Htt), mRNA CCTGGTATGTGGATCAGAAGTCCTAGCTCTTGCCAGATGGTTCTGAGCCCGCCTGCTCCACTGGGCTGGAGAGCTCCCTCCCACATTTACCCAGTAGGCATACCTGCCACACCAGTGTCTGGACACAAAATGAATGGTGTGTGGGGCTGGGAACTGGGGCTGCCAGGTGTCCAGCACCATTTTCCTTTCTGTGTTTTCTTCTCAGGAGTTAAAATTTAATTATATCAGTAAAGAGATTAATTTTAATGTAACTTTTCCTATGCCCGTGTAAAGTGTGTGACTTGGCAAGGCCTGTGCTGCATGTGACAAAGTTTATGGAAGTGGAGGGGCCTTCTGGCCGCCACTCCCTCTCCTGTAGCTACTCAGTCTAGTCGGGCAGGTCCCTCCTGTAGCCCTCCCAACACCCTGTGGCACTTGCACTTCATACAGCTCCCTTTTCTTATGCATTCCATTAAGCCAGCACAGAGAGAGGTGTTGGTATTGACTGCCTGTGTGAGAATCCTGCCTGTGGCCTAACTGAGGAACTGAAAAACTGACTTCCACTGTTAGAGTTATAAGGAGGCTTGCCCTGTGGCAGCTGCCCTCCTCTCCCCTTCCCAGGCATGACTGTCAAGCTATCTCCTCCCTGGTGTTGATGCACTCTCCTAGTCTCTCAGCCTGGGTAGAAACAGCATCTGCTGGACCCAAAGTGGCTATCCCAATAACCTCATCCCTGGTTGTGGCTGACCTGCACTGTAGCCTGCCCACACACCAGCTGACCATTGTGGATGCTGTCTGTCCCTTTGTATCTTCTGCATGGTTGGGACCTGAGAAGTGCTGACCTGATTACCCCAAAGGTGTCTCTGAGCTATGGTTTGTTGGTTTGTCTCAGTTTCTCATAGTCAAGGGAAAGCTTGGTGTCCTAGCAACAGTTAAGAATGGACCCAGAGCCTCTTTTGCCCCTTCCCATCTTGCCTTCTGTCAGCCC

### HTT1a Probe: Mm-Htt-intron1-O2 (Cat. No. 575581)

This probe targets the 5’ region of intron 1 of mouse *Htt*, which is retained in the aberrantly spliced Htt1a transcript. This region contains the cryptic polyadenylation sites used in incomplete splicing of mutant huntingtin mRNA. The target sequence is retained in the Htt1a transcript but is absent from the canonical full-length Htt mRNA. The probe consists of 20 ZZ pairs targeting region 34919494-34920643 of NC 000071.7 (1,150 bp).

#### Target sequence

>NC 000071.7:34919494-34920643 Mus musculus strain C57BL/6J chromosome 5, GRCm39 CCTGGCCTGCGTGCTGGGCATGGCCAACACTGTTCCCTGTCCAGAGGGTCGCGGTACCTCCCTGAGGCCAGGCTTTCCCGGCCCGGGCCCTCGTCTTGCGGGGTCTCTGGCCTCCCTCAGAGGAGACAGAGCCGGGTCAGGCCAGCCAGGGACTCGCTGAGGGGCGTCACGACTCCAGTGCCTTCGCCGTTCCCAGTTTGCGAAGTTAGGGAACGAACTTGTTTCTCTCTTCTGGAGAAACTGGGGCGGTGGCGCACATGACTGTTGTGAAGAGAACTTGGAGAGGCAGAGATCTCTAGGGTTACCTCCTCATCAGGCCTAAGAGCTGGGAGTGCAGGACAGCGTGAGAGATGTGCGGGTAGTGGATGACATAATGCTTTTAGGAGGTCTCGGCGGGAGTGCTGAGGGCGGGGGAGTGTGAACGCATCCAATGGGATATTCTTTTTCCAAGTGACACTTGAAGCAGCCTGTGACTCGAGGCACTTCGTACTCTCCTGGCGTTTCATTTAGTTTGTGGTGTAGTGTAGTTAAACCAGGTTTTAAGCATAGCCAGAGAGGTGTGCTTCTGTGTGTCTGCAGGCAGTTGGATGAGTTGTATTTGTCAAGTACATGGTGAGTTACTTAGGTGTGATTATTAATAAAAAACTATATGTGTGCATATATATGAAAGAGTCGACTTATACTTAACTGCCTATCGATTTTTTGTTCTATATAAAACGGATACATTGGTGGTGCTCAGTTTTCACCGGGGAATGAATTTTACTAGTGTTGCAGACAGGCTTGTTTTAGAACATAGGCCACTCTGACTCTGACTTTGTGCCAGTAAAAGTTCCTGTTTAGTTCTTTGCTGACATCTTATAGATCTTTGGAAGCTAGCTGCTTGTGACTGGAGAGAATATTGAAACAGAAGAGAGACCATGAGTCACAGTGCTCTAAGAGAAAAGAGACGCTCAAAACATTTCCTGGAAATCCATGCTGAGTGTTGAGCCCTGTGCTCTCTTGCAGCTCAGTCCTTTCTCTCAACTCTGGGCATTTTATTTCTAATCTGGATTTGTATAATTAATAAGGAGAACTTTTGGGAACAACCTACTAAAGAATGTCATCATTAAAACTCACTTAGAAAATA AGTGTTCTGGTGATATCATTGAGC

